# Independent and repeated acquisition of endosymbiotic bacteria across the diversification of feather lice

**DOI:** 10.1101/2025.01.17.633592

**Authors:** Juliana Soto-Patiño, Kimberly K. O. Walden, Jorge Doña, Lorenzo D’ Alessio, Sarah E. Bush, Dale H. Clayton, Colin Dale, Kevin P. Johnson

## Abstract

Many parasitic insects, including lice, form close relationships with endosymbiotic bacteria that are crucial for their survival. In this study, we used genomic sequencing to investigate the distribution and evolutionary history of the bacterial genus *Sodalis* across a broad range of feather louse species spanning 156 genera. Phylogenomic analysis revealed significant diversity among *Sodalis* lineages in feather lice, and robust evidence for their independent and repeated acquisition by different louse clades throughout their radiation. Among the 1,020 louse genomes analyzed, at least 22% contained *Sodalis*, distributed across 57 louse genera. Cophylogenetic analyses between the *Sodalis* and feather louse phylogenies indicated considerable mismatch. This phylogenetic incongruence between lice and *Sodalis*, along with the presence of distantly related *Sodalis* lineages in otherwise closely related louse species, strongly indicates repeated independent acquisition of this endosymbiont. Additionally, evidence of cospeciation among a few closely related louse species, coupled with frequent acquisition of these endosymbionts from free-living bacteria, further highlights the diverse evolutionary processes shaping *Sodalis* endosymbiosis in feather lice.

## Introduction

Throughout evolutionary history, many insects have established associations with intracellular, heritable bacteria (Kikuchi, 2009). These endosymbionts often inhabit specialized host cells, providing benefits such as nutritional provisioning, enhanced digestion, and protection against pathogens and environmental stresses (Sudakaran et al. 2017, McCutcheon et al. 2019). These symbiotic associations not only benefit individual insects but can facilitate specialization of lineages into diverse dietary niches (Douglas, 2009), ultimately shaping insect diversification (Cornwallis et al. 2023).

Despite growing recognition of the significance of insect-endosymbiont associations in evolutionary research (Wernegreen 2015, Provorov and Onishchuk, 2018), significant gaps remain in our understanding. Specifically, patterns of evolutionary diversification of endosymbiotic bacteria and mechanisms of acquisition by insect hosts require further investigation. Such studies can offer valuable insights into the fundamental processes of host-symbiont coevolution.

The acquisition of novel endosymbionts in insect lineages is primarily explained by two models: host-switching (horizontal transfer from other insects) and acquisition from free-living bacteria (Russell et al. 2003; Toju et al. 2013; Sudakaran et al. 2017). Host-switching involves the transfer of endosymbiotic bacteria between host lineages, resulting in a phylogenetic tree topology in which the recipient lineage becomes united with the donor lineage on a long, well-supported branch (Russell et al. 2003).

Alternatively, insects may acquire endosymbionts from free-living bacteria present in their environment. In this scenario, a free-living ancestral bacterium, characterized by a large effective population size (Moran 1966, McCutcheon et al. 2009), colonizes an insect and becomes an endosymbiont, sometimes replacing a pre-existing endosymbiont lineage (Sudakaran et al. 2017). Because free-living bacterium evolve slowly due to strong purifying selection, their acquisition by insects leads to an ancestral endosymbiont with a starting genome similar to this free-living ancestor, in a sense resetting the molecular evolutionary clock (Boyd et al. 2024). However, once a free-living bacterium transitions to an endosymbiotic lifestyle, the relaxed selection imposed on the maintenance of many genes, along with a reduced effective population size and the loss of DNA repair functions, lead to an accelerated rate of molecular evolution (Moran, 1996; Woolfit & Bromham, 2003; Boyd et al. 2024). This accelerated rate can be detected by long branches of endosymbiont lineages compared to short branches of free-living lineages in a bacterial phylogenetic tree. Repeated acquisition of the same (or similar) free-living ancestral bacteria across different insect lineages results in endosymbionts that begin from a very similar ancestral state. As these endosymbionts evolve independently within their respective hosts, they diverge into distinct endosymbiont lineages. This process results in a star-like phylogenetic tree topology, with long, independently evolving branches for each endosymbiont lineage (Smith et al. 2013; Boyd et al. 2024).

Instances of host-switching by bacteria between insect lineages have been discovered in certain psyllids and aphids, where symbionts have independently colonized both unrelated and closely related insect hosts (Thao et al. 2000; Sandström et al. 2001; Russell et al. 2003, Hall et al. 2016). For example, Russell et al (2003) conducted a phylogenetic analysis of symbionts in aphids and psyllids, revealing well-supported clades within the bacterial sequences that united aphid symbionts with those of psyllids, suggesting horizontal transfer between these insect groups. This finding reveals that the genetic similarity of endosymbiotic bacteria across diverse hosts is the result of host-switching, highlighting the significant role of horizontal transfer in shaping symbiont distributions among both distantly and closely related insect lineages.

In addition to host-switching, cases of acquisition of novel endosymbionts from free-living bacterial lineages are also known. For example, the analysis of host-symbiont phylogenetic congruence between psyllids and their endosymbionts provided compelling evidence of both bacterial host-switching and replacements by novel free-living bacterial lineages within this system (Hall et al. 2016). The replacement phenomenon occurs when one lineage of endosymbiont is replaced by another within the host insect population, with each replacement event representing the independent acquisition of a novel endosymbiont (Sudakaran et al. 2017). Previous studies have uncovered instances of endosymbiont replacement or novel acquisitions in diverse insect groups (e.g. McCutcheon et al. 2009; Koga et al. 2013; Smith et al. 2013; Šochová et al. 2017; Sudakaran et al. 2017; Chong and Moran 2018; Duron and Gottlieb 2020). For example, related groups of parasitic feather lice (Phthiraptera: Ischnocera) harbor bacterial endosymbionts from distantly related genera, suggesting the replacement of existing bacterial symbionts by free-living progenitors (Boyd et al. 2016; Smith et al. 2013; Boyd et al. 2024).

Due to their relatively simple lifecycle and specialized dietary habits, parasitic lice (Insecta: Phthiraptera) offer an outstanding system for investigating endosymbiont acquisition. As permanent parasites of birds and mammals, lice complete their entire life cycle on the host, primarily transmitting by physical contact between hosts (Clayton et al. 2016). Notably, these lice display high host specificity, with the majority of louse species being specific to only one species of host (Price et al. 2003; Johnson et al.

2011). Many species of lice have highly specialized diets, feeding exclusively on host blood or feathers, which lack essential vitamins for louse development (Sterkel et al. 2017; Alickovic et al. 2021). Consequently, lice with such specialized diets depend on heritable endosymbiotic bacteria capable of synthesizing vitamins that are lacking in their diet (Ries 1931; Buchner 1953, 1965; Puchta, 1955; Boyd et al. 2016, 2024; Říhová et al. 2017, 2022). Feather-feeding lice (Phthiraptera: Ischnocera) comprise over 3,000 described species across over 150 genera (Price et al. 2003; Smith 2001). Despite this diversity, little research has focused on the endosymbiotic bacteria that most feather lice seem to possess.

One bacterial genus that has been documented as an endosymbiont of feather lice is *Sodalis* (Fukatsu et al. 2007). Members of the genus *Sodalis* include both free-living species and insect endosymbionts, with endosymbionts found in a wide range of insect groups, including feather lice, stink bugs, mealybugs, psyllids, grain weevils, and hippoboscid flies (McCutcheon et al. 2019; Renoz et al. 2023). Among feather lice, the *Sodalis* phylogeny from the dove-louse genus *Columbicola* is star-like, characterized by weak internal node support and long terminal branches. This pattern is indicative of repeated acquisitions from a free-living ancestor, perhaps something similar to *Sodalis praecaptivus*, which is a free-living species of *Sodalis* with a notably short branch length in the phylogenetic tree (Smith et al. 2013; Boyd et al. 2024). Initially isolated from a human infection, *S. praecaptivus* provides insight into the process of endosymbiont acquisition. The frequent transitions within *Sodalis* to endosymbiosis across insects indicates a potential predisposition to repeatedly colonize diverse hosts (Clayton et al. 2012; Chari et al. 2015). This possibility makes *Sodalis* an important model for studying endosymbiont acquisition and evolution in insects.

To date, *Sodalis* endosymbiotic bacteria have been documented in a handful genera of feather feeding lice: dove lice (*Columbicola*), songbird lice (*Guimaraesiella*), and shorebird lice (*Carduiceps*, *Lunaceps*, *Quadraceps*, and *Saemundssonia*) (Smith et al. 2013; Grossi et al. 2024a; Grossi et al. 2024b; Boyd et al. 2024). However, the distribution of *Sodalis* across the diversity of feather lice (Ischnocera) is unknown.

Here, we employ genome-resolved metagenomic approaches to examine the presence of *Sodalis* across the diversity of feather lice, analyzing data from over 1,000 louse samples representing 156 feather louse genera. We use these data to reconstruct the phylogeny of *Sodalis* to understand the process of acquisition of these endosymbionts across the diversity of feather lice.

## Methods

The workflow pipeline of this project (detailed below) leverages whole genome sequencing, using metagenomic and phylogenomic techniques to resolve the cophylogenetic history of lice and their endosymbiotic bacteria.

### Taxon sampling

Samples of 1,020 chewing lice belonging to the parvorder Ischnocera (Johnson et al. 2018a) representing 156 feather louse genera (following classification of Price et al. 2003 with modifications of Gustafsson and Bush, 2017) were selected for genomic sequencing or available from previously published data (Johnson et al. 2018, de Moya et al. 2019, Virrueta Herrera et al. 2020, Johnson et al. 2021, Johnson et al. 2022, Doña and Johnson 2023, Johnson and Doña 2024, Boyd et al. 2024, Sweet et al. 2024) (supplementary material, table S1). These samples form the basis from which to explore whether *Sodalis* is present in a given louse species, and then to build a phylogeny from the resulting *Sodalis* genome sequences that were obtained (below).

### Genome sequencing

Genome sequencing, louse gene assembly, and phylogenetic analysis follow the methods described in Johnson et al. (2021). Lice were preserved in 95% ethanol and stored at -80°C. Individual lice were selected for extraction, and a photograph was taken and digitally deposited (see Data accessibility). Before extraction, individual lice were washed in a 1.5 ml vial of 100% ethanol. Total genomic DNA was extracted by first removing the louse from the vial and allowing the ethanol to evaporate. The louse specimen was then ground using a plastic pestle within a 1.5 ml tube. For the DNA extraction, a Qiagen QIAamp DNA Micro Kit (Qiagen, Valencia, CA, USA) was employed. The manufacturer’s protocol was followed, but modified by using an initial 48-hour incubation at 55 °C in buffer ATL containing proteinase K. The resulting purified and filtered DNA was finally disposed in 50ul buffer AE. The quantification of the total DNA content was performed using a high sensitive kit with a Qubit 2.0 Fluorometer (Invitrogen, Carlsbad, CA, USA).

Genomic libraries were prepared using a Hyper library construction kit from Kapa Biosystems. The libraries were sequenced using Illumina NovaSeq 6000 with S4 reagents to obtain 150 bp paired-end reads. A set of dual-end adaptors was utilized for tagging the libraries, and they were multiplexed at 48 libraries per lane, with the aim of achieving approximately 30-60X coverage of the louse nuclear genome. These reads also typically contain similar coverage of the endosymbiont genome (Boyd et al. 2024). Lastly, adapters were trimmed, and files were demultiplexed using bcl2fastq version 2.20, resulting in the generation of fastq files. For each library, the raw reads were deposited in NCBI SRA (supplementary material, table S1).

### Assembly and annotation of Sodalis sequences and phylogenomic analysis

The goal of this study was to reveal the distribution of *Sodalis* across the diversity of feather lice by analyzing 1,020 feather louse genomes (supplementary material, table S1). Therefore, we employed the reference based assembly approach Mine Your Symbiont (MinYS) (Guyomar et al. 2020) to assemble the bacterial genomes, which allowed for comparatively rapid assembly times. The assembled contigs were annotated using the Microbial Genomes Atlas (MiGA) database (Rodriguez-R et al. 2018) and tentative identification from this database was performed. MinYS uses a reference genome, in this case *Sodalis praecaptivus*, to assemble a particular genome of interest from metagenomic data. This reference-guided assembler creates initial contigs from a subset of reads, which then are fine-tuned using all metagenomic reads in a de *novo* approach. The result is a genome graph that identifies strains with possible structural variations in the samples (Guyomar et al. 2020). This approach performs well when the target genome is closely related to the reference (Guyomar et al. 2020). Thus, given that we were specifically targeting *Sodalis* endosymbionts, this approach was ideal for the current study.

In the MinYS pipeline more specifically, FASTQ reads from the louse sequencing libraries were mapped to the *Sodalis praecaptivus* reference genome (NCBI: GCF_000517425.1) using the BWA aligner. Recruited reads were then assembled into contigs with the Minia short-read assembler in MinYS. Gaps between contigs were then filled with the genome-finishing mode of MindTheGap software (Rizk et al. 2014). The final pipeline step simplified the GFA-format assembly and converted it into FASTA output.

Following assembly, we annotated the assembled contigs using the Microbial Genomes Atlas (MiGA) database (Rodriguez-R et al. 2018) to identify the closest available genomes and determine their taxonomic classification. The MiGA webserver allows the classification of unknown prokaryotic genome sequences based on the genome-aggregate Average Nucleotide and Amino Acid Identity (ANI/AAI) calculated against genomes available in two database options: ProK containing non-redundant complete and chromosomal-level assemblies in NCBI vs. TypeMat containing type material from draft and complete genomes in NCBI. We analyzed each set of MinYS-assembled contigs (minimum assembly sum >20kb) against the more complete TypeMat database using the “Popgenome” option.

To determine the presence of putative *Sodalis* contigs, we first checked the MiGA RDP Classifier output which identifies 16S rRNA sequences and assigns a confidence estimate to a particular taxon. For contigs with *Sodalis* 16S ribosomal RNA confidence >70%, we determined which sets of contigs represented intermediate to high quality genomes with low levels of contamination based on gene prediction output by Prodigal within MiGA. MiGA also leverages a Ruby script from the enveomics collection

(Rodriguez and Konstantinidis 2016) to identify 106 conserved (“essential”) genes, typically present in single-copy, across Bacteria and Archaea. For *Sodalis* contig sets containing predictions for >55 essential gene orthologs, the amino acid predictions were retrieved, as well as the genome gene prediction sets, in both amino acid and nucleotide formats. Sequences of these 106 conserved gene sets were used in phylogenetic reconstruction for *Sodalis* (below).

### Essential single-copy gene (ESS) file processing, phylogenetic matrix preparation, and tree inference

Based on the bacterial phylogeny of *Sodalis* and relatives in McCutcheon et al. (2019), we selected published genomes of 30 species from 20 bacterial genera as the outgroups for the phylogenetic analysis, with the species *Pragia fontium* and *Budvicia aquatica* used to root the phylogenetic analysis (supplementary material, table S1). These sequences were also annotated using MiGA to retrieve the same 106 essential gene set as for our novel *Sodalis* genomes. The nucleotide sequences for the essential genes were translated into amino acids using a custom python script, and multiple alignments for each gene were produced with MAFFT v7.490 using the options “—auto, -- preservecase, --adjustdirection, --amino”. The amino-acid alignments were back-translated to nucleotide sequences with a custom Phyton script. Alignment gaps were trimmed using trimAl v1.4.rev15 setting the gap threshold to “gt 0.4”. Individual gene trees were constructed using IQ-TREE v2.1.3 (-m MFP) and visualized in FigTree v1.4.4 to identify any non-*Sodalis* sequences. Sequences that were clearly not *Sodalis*, such as those identified as *Burkholderia* in a few samples, were removed to prevent the inclusion of contaminated or chimeric data in further analyses. A concatenated gene set in FASTA format was then created using the AMAS concat function with a partitions file in nexus format (Borowiec 2016). The final concatenated gene set tree was built in IQ-TREE v2.1.3 using the General Time Reversible model, the Discrete Gamma model with 4 categories for rate heterogeneity, and 1000 ultra-fast bootstrap replicates.

### Gene assembly and phylogenomic analysis for lice

#### Louse sequence assemblies and phylogenomic analysis

In our study, we sought to reconstruct the cophylogenetic relationships between *Sodalis* bacteria and their feather louse hosts. Thus, we also needed a tree for the lice that contained a *Sodalis* endosymbiont. To achieve this, we processed the raw genomic data from each feather louse sample that was determined to harbor *Sodalis*, assembled a set of ortholog genes, and conducted a phylogenomic analysis following the methods described in Johnson et al. (2021). More specifically, raw reads were processed using fastp v0.20.1 for adapter trimming and quality control (Chen et al. 2018). We used aTRAM 2.0 (Allen et al. 2018) to assemble 2,395 single copy orthologs from a reference set of protein-coding genes (Johnson et al. 2018a) from the human louse, *Pediculus humanus.* We translated the nucleotide sequences to amino acids using a custom Python script and then, we performed a phylogenomic analysis, first aligning amino acid sequences using MAFFT v7.471 (Katoh et al. 2002, Katoh and Standley 2013) and then back-translating them to DNA sequences. The gene alignments were trimmed using trimAL v1.4. rev22 (Capella-Gutierrez et al. 2019). The resulting gene alignments were concatenated into a supermatrix using AMAS v1.0. (Borowiec 2016). For phylogenetic analysis, we employed IQ-TREE 2 v2.1.2 (Minh et al. 2020) with parameters for partitioning and model selection to reconstruct a tree based on the concatenated gene sequences. We rooted the tree using *Proechinophthirus fluctus* (Anoplura), a species with a published *Sodalis* endosymbiont genome (Boyd et al. 2016). Since Anoplura (sucking lice) and Ischnocera (feather lice) are closely related sister groups, this allowed us to focus on lice with the *Sodalis* endosymbiont for direct comparison with the evolutionary patterns of the *Sodalis* bacteria (above). Ultrafast bootstrapping with UFBoot2 was used to assess tree support (Minh et al. 2013; Hoang et al. 2018). To account for incomplete lineage sorting, individual gene trees were generated with IQ-TREE 2 (-m MFP) and used in a coalescent analysis to construct a species tree with ASTRAL-III (Zhang et al. 2018). This software also calculated local posterior probabilities for each node in the coalescent tree.

#### Cophylogenetic analysis

We compared the partitioned concatenated louse and *Sodalis* endosymbiont trees using eMPRess v1.0 (Santichaivekin et al. 2020). For this comparison, we pruned our overall bacterial tree to contain only *Sodalis* taxa from lice. This software summarizes events across equally parsimonious cophylogenetic reconstructions into median maximum parsimony reconstructions (mMPR). We followed the cost scheme (duplication: 1, sorting: 1, and host-switching: 2) used in various published cophylogenetic studies on lice (Sweet et al. 2016; de Moya et al. 2019; Johnson et al. 2021). This cost scheme makes the combined weight of duplication and sorting equal to the weight assigned to host-switching, providing an alternative method for reconstructing conflicting nodes in parasite and bacterial trees. Notably, cospeciation is consistently assigned a zero cost in the techniques of cophylogenetic reconstruction. Because switching of endosymbionts between species of lice might be implausible, given that they are isolated on different bird host species, we also performed the eMPRess analysis to minimize host-switching events by setting the host-switching cost parameter to 15 (following Boyd et al. 2024). The other cost parameters were left at the prior values (0 for cospeciation events, 1 for duplication, and 1 for losses). To test whether the reconstructed cost was less than expected by chance we randomized the *Sodalis* tree 100 times to compare the cost for the reconstruction of the actual trees to those from the randomized distribution. This essentially tests whether the *Sodalis* tree is more similar (i.e. contains more cospeciation events) than expected by chance. Additionally, we used the louse and bacteria phylogenies to build a tanglegram, using the R package phytools (version 2.3-0) (Revell 2024).

## Results

### Identification and distribution of Sodalis endosymbionts

The Illumina sequencing of genomic libraries derived from individual lice yielded a range of 19 to 110 million total 150 bp reads (Read1 + Read2) per sample. Using MinYS assembly with MiGA annotation, we obtained robust assemblies of *Sodalis* from 228 of the 1,020 (22.35%) feather-feeding louse genomes that we examined (supplementary material, table S1). Of the 156 genera of feather lice, 57 of them harbored bacterial sequences sharing high identity with *Sodalis spp.* (supplementary material, table S2). From assemblies, we also found evidence of *Sodalis* in another 25 samples, but these did not maintain sufficient coverage to be deemed suitable for inclusion in further analyses. Some genera of feather lice tended to have a high fraction of representatives harboring *Sodalis* endosymbionts (positives), while in others no *Sodalis* was detected (negatives) (supplementary material, table S2). For example, in genera with more than 20 samples represented in the study, prevalence of *Sodalis* ranged from 6% to 88%: 88.4% (23/26) in *Brueelia*; 75% (15/20) in *Picicola;* 20.6% (6/29) in *Anaticola*; 22.2% (10/45) in *Quadraceps*; 42.5% (20/47) in *Guimaraesiella*; 48.7% (38/78) in *Columbicola*; and 6.2% (5/81) in *Rallicola*. Cases in which multiple individuals of the same louse species were sequenced generally indicated that these individuals harbor the same or near identical lineages of *Sodalis* endosymbionts. For example, two individuals of *Columbicola tasmaniensis* (from two different dove hosts) had *Sodalis* endosymbionts differing by only 0.22% uncorrected pairwise sequence divergence across all ESS genes combined. Likewise, two individuals of *Strongylocotes* sp. from *Crypturellus soui* harbored *Sodalis* that differed by only 0.03%, and two individuals of *Saemundssonia wumisuzume* harbored *Sodalis* differing by only 0.21%. Some lice have genetically differentiated populations or cryptic species (Johnson et al. 2002). In many of these cases, the *Sodalis* from related louse individuals were closely related, yet genetically distinct. For example, samples of *Columbicola extinctus* from Band-tailed Pigeons (*Patagioenas fasciata*) in the U.S. versus Peru harbored *Sodalis* endosymbionts that were sister taxa, yet genetically distinct (6.97% different).

Similarly, cryptic species of dove lice, *Columbicola passerinae* 1 and 2, had related *Sodalis* species that were genetically differentiated (6.29%). Another example occurs between two individuals of the parrot louse *Neopsittaconirmus circumfasciatus* on *Alisterus chloropterus* and *Alisterus scapularis*, which had *Sodalis* differing by 4.06%.

### Phylogenetic patterns in Sodalis

The phylogeny resulting from IQTREE analyses of bacteria within Enterobacterales, including *Sodalis*, was generally very well resolved and supported in terms of relationships between bacterial genera. However, the overall topology of the tree within the genus *Sodalis* reveals a star-like pattern. Specifically, the backbone of relationships among *Sodalis* endosymbionts of feather lice was characterized by short branches and low bootstrap support values. However, this phylogeny did reveal two main clades of *Sodalis*, each with 100% bootstrap support. Both of these clades included insect endosymbionts. Clade A included *Sodalis glossinidius*, the well-studied endosymbiont of the tsetse fly, along with *Sodalis* lineages from the louse genera *Quadraceps* (four species), *Mulcticola* (one species), and *Cirrophthirius* (one species). Clade B comprised the vast majority of feather louse associated *Sodalis*. It also included *S. pierantonius* (a nascent/recently-derived grain weevil symbiont) and the free-living *Sodalis praecaptivus*, which are closely related to the majority of *Sodalis* strains from feather-feeding lice. In our analysis, we also included *Sodalis baculum* (a seed bug symbiont, Santos-Garcia et al. 2017), which is also a member of Clade B, clustering with other louse endosymbionts on a relatively long branch. In addition, *S. melophagi* (a sheep ked symbiont, Chrudimsky et al. 2012) falls within Clade B, but on a relatively shorter branch, again clustering with other louse endosymbionts.

One notable feature of the phylogeny of *Sodalis* was the extreme variation in branch lengths. As in a prior analysis (Boyd et al. 2024), the free-living *Sodalis praecaptivus* was placed on a short terminal branch in comparison with endosymbiont lineages. However, some endosymbionts of lice were also on very short terminal branches. For example, *Sodalis* from *Philoceanus robertsi* was on a very short terminal branch, not markedly dissimilar to *S. praecaptivus*. In contrast, certain *Sodalis* lineages, such as the *Ibidoecus flavus* endosymbiont, were on very long branches, indeed the longest branch in the entire phylogeny. Other *Sodalis* taxa in feather lice with the longest branches, including that of *Ibidoecus flavus*, tended to cluster together.

However, many of the nodes uniting these long-branch taxa had low bootstrap support (<90%), perhaps indicative of the artifact of long-branch attraction (Felsenstein 1978) or possibly arising from a common base compositional bias, often A+T bias, as documented in many symbiont lineages (Moran 2003).

### Louse gene assembly and phylogenomic analysis

Assuming a genome size of 200–300 Mbp for Ischnocera (Baldwin-Brown et al. 2021; Sweet et al. 2023), coverage of louse genomes ranged from around 20X to 100X. Assemblies of 2395 single copy ortholog genes using aTRAM 2 (Allen et al. 2018) resulted in assemblies ranging from 872 to 2352 genes, depending on the sample, with an average of 2316 genes. After alignment, we retained 2376 genes for phylogenomic analysis. Following trimming, the concatenated alignment consisted of 3,857,202 aligned base positions. The analysis in IQ-TREE identified 432 optimal partitions with separate ML models, producing a fully resolved tree with 100% bootstrap support for all but five branches, which were 98-99% (Fig. S1). ASTRAL-III coalescent searches produced a nearly identical tree. The branching pattern of the main lineages of feather lice was generally identical to that of prior studies (Johnson et al. 2018b, de Moya et al. 2019), albeit the taxon sample of the current tree was limited to those lice for which we found *Sodalis* as a likely endosymbiont.

### Cophylogenetic analysis

When examining host distribution, the phylogenetic relationships of *Sodalis* across different feather louse genera, show a mix of patterns. In several cases, some lineages of *Sodalis* from the same louse genus are closely related (*e.g. Anaticola*, some *Formicaphagus*). In many other cases, *Sodalis* from lice in the same genus are spread throughout the tree (*e.g. Columbicola*, *Strongylocotes*, *Brueelia*). These patterns suggest there could be a mix of codivergence between lice and *Sodalis* and phylogenetic incongruence.

We tested these patterns more formally by employing a cophylogenetic analysis. In particular, we aimed to assess the level of congruence between the host (louse) and endosymbiont trees. Generally, cophylogenetic reconstruction methods allow for cospeciation, host-switching, duplication, and sorting events. For the cost scheme employed in many cophylogenetic studies in which cospeciation is 0, host-switching 2, duplication 1, and sorting events 1, the cophylogenetic reconstruction in eMPRess that included every sample as a terminal taxon, reconstructed 73 cospeciation events, 0 duplications, 154 host-switches, and 9 losses. The cost for this reconstruction is much less than that for random trees (*P* < 0.01), indicating more cospeciation events than expected by chance. Cophylogenetic reconstruction methods do not currently account for a scenario of repeated acquisition, so we repeated the analysis after increasing the cost of host-switching to 15, following Boyd et al. (2024), to more closely simulate a scenario of repeated acquisition, while minimizing inferred host-switching. In this case, inferred duplication events and losses provide a measure of independent acquisition (Boyd et al. 2024). In this scenario, the reconstruction returned 125 cospeciation events, 91 duplications, 11 host-switches, and 1,113 losses while congruence between the *Sodalis* and louse trees was still found to be significant (*P* < 0.01).

## Discussion

Phylogenomic analysis of *Sodalis* endosymbionts of feather-feeding lice revealed significant diversity of *Sodalis* lineages and robust evidence for their independent and repeated acquisition by this group of insects. Among the 1,020 louse genomes analyzed, 22.35% contained evidence of associated *Sodalis* genomes, distributed across 57 genera of lice. The widespread, but not universal, nature of these endosymbiotic associations suggests *Sodalis* in feather lice have undergone multiple acquisitions and losses (Baum 2005; Moran et al. 2008). There are several lines of evidence (Boyd et al. 2024) supporting this conclusion.

First, the phylogeny of *Sodalis* endosymbionts exhibits a star-like topology with many long terminal branches connected by short, weakly supported internodes. In addition, in many cases, closely related species of lice harbor *Sodalis* endosymbionts widely separated in the tree. This pattern is reflected in considerable incongruence between the feather louse and *Sodalis* trees, resulting in a tangled cophylogenetic pattern. However, there is also evidence for consistency of the same bacterial lineage between individuals of a given louse species, and for some shorter term codivergence between closely related species of lice and their *Sodalis* endosymbionts. This supports previous findings from fluorescent in situ hybridization (FISH; Fukatsu et al. 2007) studies that *Sodalis* in feather lice is maternally transmitted through the eggs, predicting a pattern of host-endosymbiont codiversification. In short, a mechanism of maternal transmission, together with the overall characteristics of the *Sodalis* phylogeny (below), provide evidence for repeated acquisition and replacement of *Sodalis* endosymbionts throughout the history of diversification of feather lice.

### Star-like phylogeny

Our phylogenetic results indicate that the process of repeated acquisition of *Sodalis* by feather-feeding lice is occurring more broadly, not just within a few genera that have been previously studied (Smith et al. 2013; Grossi et al. 2024a,b; Boyd et al. 2024). One of the key findings of our study is the star-like phylogeny of *Sodalis* endosymbionts associated with feather lice, characterized by long terminal branches and very short internal node that are weakly supported. In addition, these *Sodalis* endosymbionts are very closely related to the free-living *Sodalis praecaptivus*, which is on a very short terminal branch. Together, these patterns suggest that many independent acquisitions of *Sodalis* from a free-living ancestor have occurred throughout the radiation of feather-feeding lice (Smith et al. 2013; Boyd et al. 2024). This scenario of recurrent acquisitions is supported by previous studies demonstrating that free-living *Sodalis* have transitioned to an endosymbiotic lifestyle many times in distantly related insect hosts (McCutcheon et al. 2019; Renoz et al. 2023; Boyd et al. 2024).

In the context of feather lice, the star-like topology indicates recurrent symbiont acquisition within a relatively short evolutionary timeframe, as suggested by evolutionary simulations (Smith et al. 2013). In these simulations, a free-living ancestral bacterium transitions to endosymbiosis multiple times. In the free-living lineage, molecular evolution proceeds at a slower rate, because a large effective population size allows natural selection to purge even very slightly deleterious mutations. In contrast, once a bacterial lineage becomes an endosymbiont, selection in the novel environment, combined with a much smaller effective population size within an individual host insect, results in an accelerated rate of molecular evolution, leading to long terminal branches for endosymbiont lineages. Many *Sodalis* endosymbionts have also lost mutation repair genes (Boyd et al. 2024), which is expected to further accelerate their rate of mutation. These processes, together with the fact that symbiont genomes evolve in a strictly reductive manner, yield a scenario in which the symbiont gene inventories are subsets of their free-living progenitors as evidenced by comparison with the extant, close free-living relative, *S. praecaptivus* (Boyd et al. 2024). Likewise, the multiple endosymbiont lineages derived from this ancestor do not fundamentally have a phylogenetic structure, yielding short, weakly supported internal branches.

Star-like phylogenies in diverse systems are often ascribed to dynamic evolutionary processes, such as frequent endosymbiont gains and losses. In various insect hosts, including weevils, stinkbugs, and louse flies, *Sodalis* endosymbionts exhibit similar phylogenetic patterns characterized by long branches and weak internal node support, indicating independent acquisitions from environmental reservoirs (Toju et al. 2013; Hosokawa et al. 2015; Šochová et al. 2017). For example, *Sodalis*-allied symbionts in *Sitophilus* weevils demonstrate a dynamic evolutionary history with frequent re-associations, acquisitions, horizontal transfers, replacements, and losses (Toju et al. 2013; Oakeson et al. 2014). In louse flies, the phylogenetic clustering and occasional replacements of *Sodalis* suggest multiple independent acquisitions over evolutionary time. Similarly, stinkbugs show variable *Sodalis* infection frequencies across species combined with host-symbiont phylogenetic incongruence (Hosokawa et al. 2015).

These recurring acquisitions suggest that *Sodalis* bacteria have repeatedly transitioned between free-living and endosymbiotic lifestyles across diverse insect taxa, underscoring their adaptive versatility in establishing endosymbiotic relationships (Renoz et al. 2023).

### Phylogenetic and cophylogenetic relationships of *Sodalis* endosymbionts

Another indicator that repeated acquisition from a free-living ancestor may be occuring is that closely related louse species often harbor distantly related *Sodalis* strains. For example, some members of the louse genus *Quadraceps* harbor *Sodalis* endosymbionts from Clade A, while others harbor representatives from Clade B. This phylogenetic pattern suggests that these bacteria have been independently acquired by different louse species through multiple evolutionary events (Smith et al. 2013), leading to significant evolutionary divergence among endosymbionts even in closely related hosts. Within Clade B *Sodalis*, this pattern is also evident, with some louse genera (*e.g. Columbicola*, *Brueelia*, and *Strongylocotes*) having *Sodalis* that appear in multiple positions throughout the phylogeny of Clade B. These patterns are reflected in widespread discordance between the louse and *Sodalis* trees.

Cophylogenetic comparisons of the louse and *Sodalis* trees revealed that the phylogenies of *Sodalis* endosymbionts and feather lice are largely incongruent. Vertical transmission of feather louse endosymbionts (Fukatsu et al. 2007, Smith et al. 2013) would normally be expected to result in a pattern of widespread codivergence, where the phylogenies of the host and endosymbiont mirror each other. Such congruence has been observed in many other insect-endosymbiont systems, including psyllids and *Carsonella* (Thao et al. 2000), aphids and *Buchnera* (Clark et al. 2000), whiteflies and *Portiera* (Thao and Baumann, 2004), weevils and *Nardonella* (Conord et al. 2008), and bat flies and *Aschnera* (Hosokawa et al. 2012), among others. However, our cophylogenetic analysis of feather lice and their *Sodalis* endosymbionts revealed a relatively low number of cospeciation events (red dots and connecting lines, Figure 2). These mainly occurred between very closely related terminal louse taxa or between cryptic species of lice, as has been found in other recent studies within single genera of feather lice (Grossi et al. 2024a, Boyd et al. 2024). Frequent replacement of endosymbionts is predicted to overwrite evidence of past louse-endosymbiont cospeciation events (Boyd et al. 2024), leading to the overall patterns observed. Thus, while vertical transmission and cospeciation does occur, independent acquisition seems to be prevalent across the diversification of feather lice, especially over longer evolutionary timescales.

**Figure 1.**
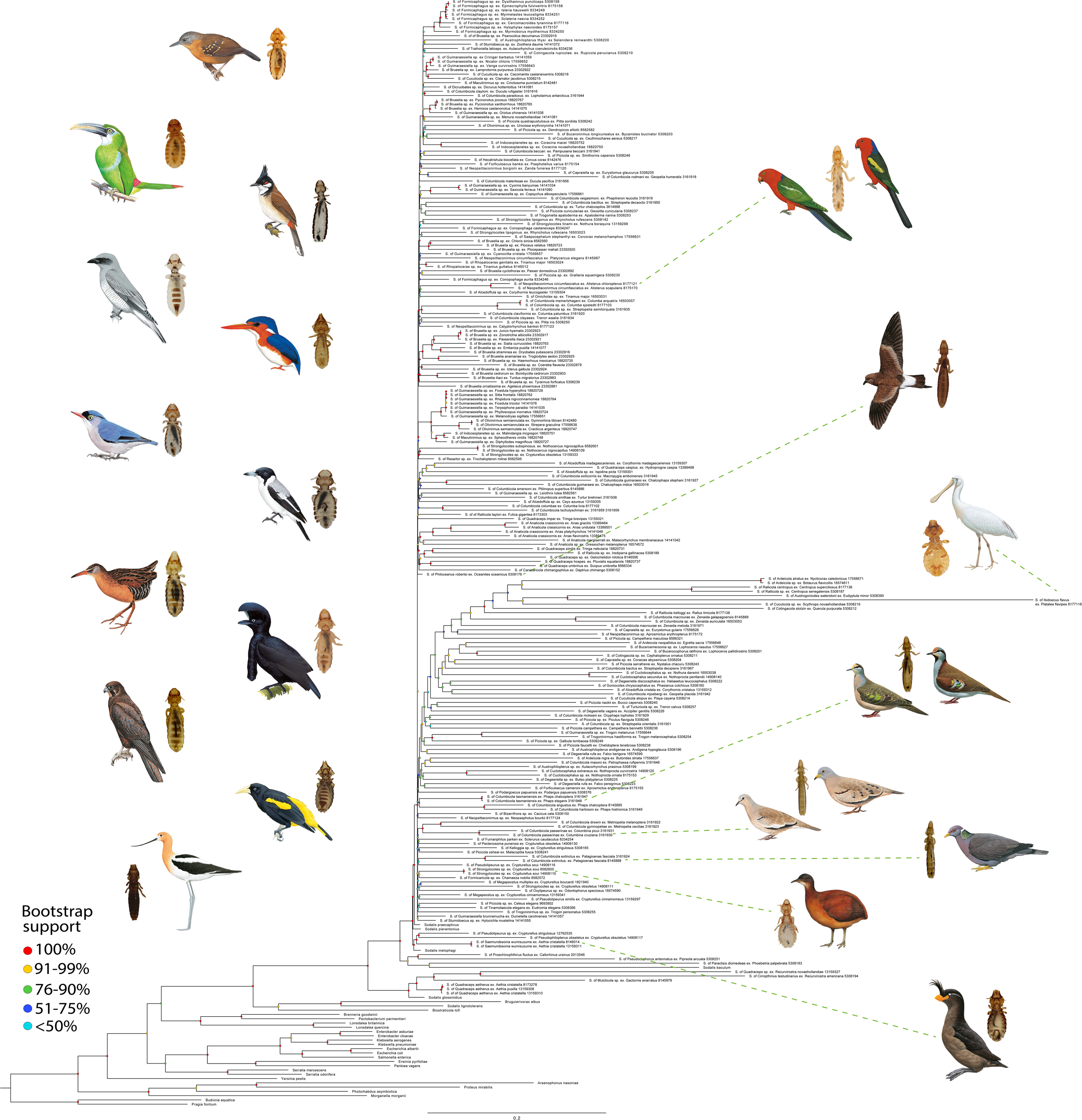
**Phylogeny of *Sodalis* spp. symbionts from feather-feeding lice**. Phylogeny of *Sodalis* spp. symbionts from feather lice (Ischnocera) and related bacteria based on a partitioned IQ-TREE ML analysis using the General Time Reversible (GTR) model and the Discrete Gamma model 4 categories for rate heterogeneity. Tree support assessed with 1000 ultra-fast bootstrap replicates. Values of bootstrap support on branches are indicated by color bullets as follows: light blue (<50%), dark blue (51-75%), green (76- 90%), yellow (91-99%), and red (100%). Branch lengths are proportional to substitution per site. Tips names indicate the *Sodalis* strain from each louse and avian host followed by respective SRA NCBI accession numbers. Bird and louse images on the left represent diverse examples of lice and their avian hosts from which *Sodalis* was identified. Bird and louse images on the right represent louse host of *Sodalis* that are mentioned in the text with lines connecting these lineages to associated images. Bird illustrations reprinted by permission from © Lynx Nature Books and © Cornell Lab of Ornithology (see supplemental material, Table S3 for licensing details).

**Figure 2.**
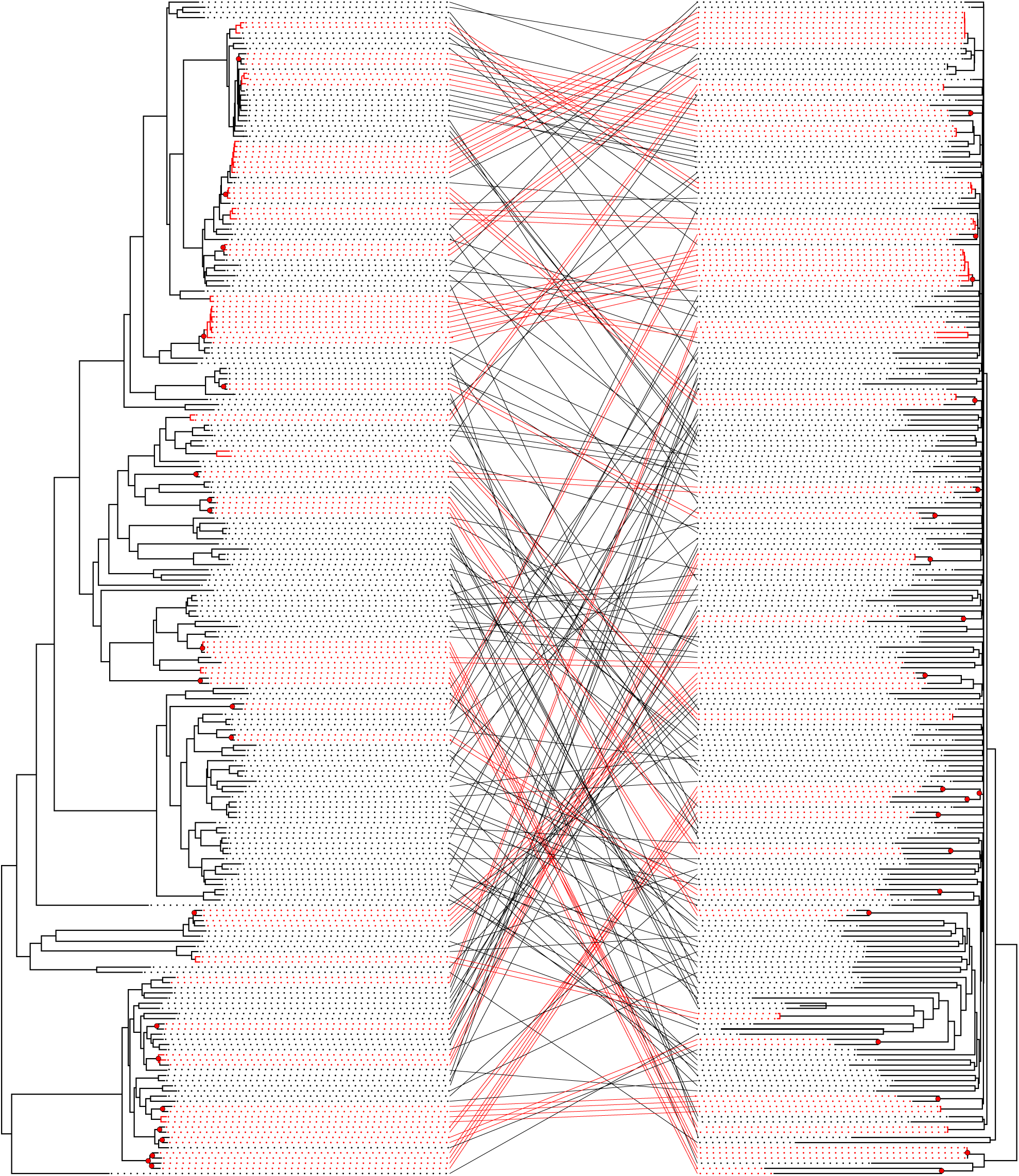
Tanglegram comparing the phylogeny of feather lice (left) with the phylogeny of their *Sodalis* endosymbionts (right). The louse tree was estimated from a partitioned IQ-TREE ML search of a concatenated matrix of 2359 single copy ortholog genes (same orientation as Figure S1). The endosymbiont tree is the same as depicted in Figure 1, but excludes the outgroups. Red branches, connecting dots and lines represents the same louse-symbiont species based on COI distances. Bulleted nodes connected with Figure S1. Phylogenetic tree of feather-feeding lice (Ischnocera) based on a partitioned IQ-TREE maximum likelihood (ML) analysis of a concatenated matrix of 2,359 single- copy ortholog genes. Bootstrap values are 100%, except where indicated. Hash marks on branch leading to *Quadraceps hospes* shorten the visually long branch resulting from missing data for this species. The tree topology is rotated to match the structure of the louse tree in the tanglegram shown in Figure 2. Tree rooted on Anoplura (*Proechinophthirus fluctus*), not shown.red lines indicate cospeciation events.

### Host-switching

There are a few cases in the overall *Sodalis* phylogeny that are not consistent with either codivergence or a pattern of repeated acquisition from a free-living bacterial ancestor.

These are cases involving *Sodalis* from somewhat distantly related louse hosts that are united on a comparatively longer, well-supported internal branch, which would be a phylogenetic pattern consistent with host-switching (horizontal transfer) of an endosymbiont from one louse lineage into another. One case occurs within the genus *Columbicola*, in which *Sodalis* from two species (*C. columbae* and *C. tsuschulysman*) are supported as sister taxa on a comparatively long, well-supported internal branch even though these lice are not closely related within the phylogeny of the louse genus *Columbicola* (Boyd et al. 2017). These two louse species occur on the same species of bird (the Rock Pigeon [*Columba livia*]), and this may provide an opportunity for an endosymbiont to switch from one louse host into another.

Another more complex case of potential horizontal transfer involves *Sodalis* in several species in multiple genera (*Guimaraesiella*, *Olivnirmus, Indoceoplanetes,* and *Maculinirmus*) of lice within the *Brueelia*-complex. The *Sodalis* lineages (at least three) in these species are united by a long, well-supported internal branch. Unlike the case in *Columbicola*, these lice do not occur on the same species of bird, but do occur in the same general biogeographic region. Given that other *Sodalis* from some of these genera occur in other places in the *Sodalis* tree, it could be possible that these lineages represent a shared *Sodalis* ancestor in a common ancestor of the *Brueelia-*complex with subsequent loss and replacement. However, the phylogenetic relationships among the *Sodalis* lineages within this clade do not directly mirror the relationships of their louse hosts. Instead, this seems most likely to be a case of ancestral contact between ancestral louse lineages resulting in the transfer of *Sodalis* from one louse lineage into others.

Further investigation with additional sampling from these genera could be revealing as to the nature of this case. Overall, however, the phylogenetic pattern of the tree suggests that host switching of *Sodalis* between louse lineages is comparatively rare, if it occurs at all.

### Variation in branch lengths

Although not necessarily directly related to the process of repeated acquisition, the significant variation in branch lengths within the *Sodalis* phylogeny reveals a dynamic evolutionary landscape for these endosymbionts. Terminal branch lengths vary by more than an order of magnitude across the tree. *Sodalis praecaptivus* occurs on a very short terminal branch, likely indicating the slow rate of molecular evolution within free-living bacteria that have very large effective population sizes. However, even within *Sodalis* endosymbionts of feather lice, there are some terminal branches that are remarkably short. For example, the *Sodalis* from *Philoceanus robertsi* is on a very short terminal branch, comparable to *Sodalis praecaptivus*, indicating either a slow rate of molecular evolution or a very recent acquisition from a free-living ancestor. Most other *Sodalis* strains in feather lice, however, occur on much longer terminal branches with the longest being *Sodalis* from *Ibidoecus flavus*. This species forms a cluster together with many of the other longest branches in the tree, albeit with relatively low bootstrap support among these branches (<90%). This clustering may reflect the artifact of long branch attraction (Felsenstein 1978) rather than any phylogenetic relationship. A further complication is that long branch *Sodalis* strains also tend to have higher AT base composition (Boyd et al. 2024), which may further cause long-branch taxa to cluster together. Overall, we posit that each of these long-branch taxa is an example of an independent acquisition event, and the application of phylogenetic analysis forces a tree structure among taxa, when in reality evolution did not proceed in a bifurcating fashion. Rather, independent origins from the same (or similar) free-living ancestor would produce the star-like phylogeny with variation in branch lengths that is observed in this study.

### Future directions

Our study of *Sodalis* endosymbionts in diverse feather-feeding lice reveals promising directions for future research in evolutionary biology and symbiosis. We found strong evidence supporting independent and repeated acquisitions of these bacteria, posing several key questions for further exploration. Feather lice offer unique opportunities as a model to study repeated instances of the process of genome evolution across multiple endosymbiont acquisition events from related free-living bacteria (Boyd et al. 2024).

The well-characterized genome of *Sodalis praecaptivus*, which can be cultured, allows for detailed investigation into the consistent retention or loss of specific genes.

Additionally, studying feather lice can shed light on evolutionary contingencies and mechanisms of genome evolution, including base composition change and the acceleration of mutation rate (Boyd et al. 2024). For example, recent findings by Boyd et al. (2024) concerning *Sodalis* endosymbionts of the louse genus *Columbicola* highlight that while genome degeneration in these symbionts is largely deterministic, stochastic processes shape the loss of genes with redundant functions, resulting in patterns of gene loss that are strongly influence by contingency.

Another interesting question is why some groups of feather lice appear to have such a high prevalence of *Sodalis* endosymbionts, while others do not. Environment or geography may play a role in which bacteria are available for acquisition. However, many of the genera of lice included in our study are geographically widespread (like their avian hosts), and there is currently no clear pattern, beyond louse phylogeny, in the pattern of distribution of *Sodalis* across feather lice.

In conclusion, our study provides robust evidence for the independent and repeated acquisition of *Sodalis* endosymbionts in feather-feeding lice. By leveraging whole genome sequencing and phylogenomic techniques, we have elucidated the distribution and evolutionary dynamics of these symbionts across diverse louse genera. Our findings contribute significant insights into the evolutionary patterns and mechanisms driving endosymbiont acquisition in insect-bacteria associations.

## Ethics

Research on animals was conducted under University of Illinois, Champaign, Illinois IACUC protocols 10119, 13121, and 15212.

## Data accessibility

Raw genomic reads for each sample have been deposited in NCBI SRA (Table S1). Code and data for running analyses, including Concatenated data matrix, gene alignments, gene trees, and all tree files are available from the FigShare digital repository https://figshare.com/s/9b914a733dbac506c7ef. Voucher lice photos are

deposited in the Price Institute of Parasite Research, University of Utah, Salt Lake City, US (Catalog numbers: PIPR050302-PIPR051322; accessible via the "Symbiota Collections of Arthropods Network" at https://scan-bugs.org and the “Ecdysis portal for live-data arthropod collections” at https://ecdysis.org), and at the FigShare digital repository https://figshare.com/s/3ebae5aea796e59cc3be. This dataset includes high-resolution photos of the specimens analyzed in the study. Each photo corresponds to a unique specimen identifier, providing visual documentation for verification and reference.

## Supplementary material

Electronic supplementary material is available online at the FigShare digital repository https://figshare.com/s/af82b1916754a9fc0ee3

## Declaration of AI use

We have not used AI-assisted technologies in creating this article.

## Author’s contributions

J.S-P.: Conceptualization, Data curation, Formal analysis, Investigation, Methodology, Visualization, Writing – original draft. K.K.W.: Data curation, Formal analysis, Methodology, Writing – original draft. J.D.: Formal analysis, Methodology, Writing – review & editing, L.D.: Formal analysis, Methodology, Writing – review & editing.

S.E.B.: Conceptualization, Funding Acquisition, Resources, Writing – review & editing, D.H.C.: Conceptualization, Funding Acquisition, Resources, Writing – review & editing, C.D.: Conceptualization, Funding Acquisition, Writing – review & editing, K.P.J.: Conceptualization, Funding Acquisition, Project administration, Resources, Supervision, Writing – original draft.

All authors gave final approval for publication and agreed to be held accountable for the work performed herein.

## Conflict of interest declaration

We declare we have no competing interests.

## Funding

This work was supported by U.S. NSF DEB-1239788, DEB-13426045, DEB-1926919, DEB-1925487, and DEB-2328118 to K.P.J. and by DEB-1926738 to C.D., S.E.B., and D.H.C.

## Acknowledgments

We thank C. Adam, R.J. Adams, A. Alexio, S. Barker, J. Bates, B. Benz, S. Bertelli, I. Beveridge, S. Cameron, H. Campbell, M. Carbajal, T. Catanach, J. Cheriton, T. Chesser, M. Cottam, L. Cueto, F. Daunt, B. Dawson, E. Diblasi, H. Eascott, R. Empson, H. Fandel, R. Faucett, R. Furness, A. Garitano-Zavala, T. Galloway, N. Gawani, T. Gnoske, S. Goodman, A. Gouvea, D. Gustafsson, J. Hagelin, C. Harbison, N. Hoffman, R. Jakob-Hoff, J. Jankowski, J. Jolly, R. Junge, J. Kaderitz, S. Kenney, J. Kirchman, J. Klicka, A. Krater, E. Kuschel, D. Lane, P. Loi, H. Lutz, B. Marks, J. Malenke, I. Mason, K. McCracken, J. Merkel, M. Meyer, M. Miller, R. Moyle, E. Neutze, B. O’Shea, E. Osnas, R. Palma, R. Palmer, V. Piacentini, A. Porzecanski, D. Roach, M. Robbins, K. Rose, J. Sailer, D. Santiago-Alarcon, J. Scherer, F. Schunck, F. Sheldon, V. Smith, S. Sontshugen, D. Steadman, A. Sweet, S. Trewick, W. Veronesi, D. Verrier, S. Volponi, J. Weckstein, N. Whiteman, D. Willard, R. Wilson, B. Winger, C. Witt, J. Wombey, B. Zonfrillo for assistance in obtaining specimens for this study. We thank A. Hernandez and C. Wright at the University of Illinois Roy J. Carver Biotechnology Center for assistance with Illumina sequencing.

---

Table S1. Louse samples analyzed, including the genus and species, their vertebrate host species, country of origin, NCBI SRA accession numbers, and the presence of *Sodalis* endosymbionts. Presence indicates whether the *Sodalis* endosymbiont was detected in the sample (Positive/Negative). Each row corresponds to a distinct sample.

Table S2. Summary of *Sodalis* detection in louse samples by genus. This table shows the counts and percentages of positive and negative samples for each louse genus analyzed. It includes the total number of samples, as well as the proportion of samples that tested positive or negative for *Sodalis*.

Table S3. Details of the bird images used in Figure 1, including the English names, scientific names, artist or author of the illustrations, and credit and copyright information.

Figure S1. Phylogenetic tree of feather-feeding lice (Ischnocera) based on a partitioned IQ-TREE maximum likelihood (ML) analysis of a concatenated matrix of 2,359 single copy ortholog genes. Bootstrap values are 100%, except where indicated. Hash marks on branch leading to Quadraceps hospes shorten t 1140 he visually long branch resulting from missing data for this species. The tree topology is rotated to match the structure of the louse tree in the tanglegram shown in Figure 2. Tree rooted on Anoplura (Proechinophthirus fluctus), not shown.

## Notes

### Competing Interest Statement

The authors have declared no competing interest.

