## Supplementary_material for "Independent and repeated acquisition of endosymbiotic bacteria across the diversification of feather lice"

**Table S1.** Louse samples analyzed, including the genus and species, their vertebrate host species, country of origin, NCBI SRA accession numbers, and the presence of *Sodalis* endosymbionts. Presence indicates whether the *Sodalis* endosymbiont was detected in the sample (Positive/Negative). Each row corresponds to a distinct sample.

| Count_number | Sodalis_detection | Library_code | Louse_Genus | Louse_Species | Vertebrate_Host | Collection_Country | NCBI_SRA |
|---|---|---|---|---|---|---|---|
| 1 | Positive | FmspDypun | *Formicaphagus* | *sp.* | *Dysithamnus puncticeps* | Panama | SRR5308158 |
| 2 | Positive | FmspMyful | *Formicaphagus* | *sp.* | *Epinecrophylla fulviventris* | Panama | SRR8175158 |
| 3 | Positive | FmspMyhau | *Formicaphagus* | *sp.* | *Isleria hauxwelli* | Brazil | SRR8334249 |
| 4 | Positive | FmspScleu | *Formicaphagus* | *sp.* | *Myrmelastes leucostigma* | Brazil | SRR8334251 |
| 5 | Positive | FmspScnae | *Formicaphagus* | *sp.* | *Sclateria naevia* | Brazil | SRR8334252 |
| 6 | Positive | FmspCetyr | *Formicaphagus* | *sp.* | *Cercomacroides tyrannina* | Panama | SRR8177116 |
| 7 | Positive | FmspHynae | *Formicaphagus* | *sp.* | *Hylophylax naevioides* | Panama | SRR8175157 |
| 8 | Positive | FmspMymyo | *Formicaphagus* | *sp.* | *Myrmoborus myotherinus* | Brazil | SRR8334250 |
| 9 | Positive | BrspPsdec | *Brueelia* | *sp.* | *Psarocolius decumanus* | Panama | SRR23302919 |
| 10 | Positive | Authy | *Austrophilopterus* | *thysi* | *Selenidera reinwardtii* | Peru | SRR5308200 |
| 11 | Positive | BrspZodam | *Sturnidoecus* | *sp.* | *Zoothera dauma* | China | SRR14141072 |
| 12 | Positive | BrlatAucoe | *Traihoriella* | *laticeps* | *Aulacorhynchus coeruleicinctis* | Bolivia | SRR8334236 |
| 13 | Positive | Corup | *Cotingacola* | *rupicolae* | *Rupicola peruvianus* | Peru | SRR5308210 |
| 14 | Positive | BrspCrbar | *Guimaraesiella* | *sp.* | *Criniger barbatus* | Ghana | SRR14141059 |
| 15 | Positive | GuspNichl | *Guimaraesiella* | *sp.* | *Nicator chloris* | Ghana | SRR17556652 |
| 16 | Positive | GuspVacur | *Guimaraesiella* | *sp.* | *Vanga curvirostris* | Madagascar | SRR17556643 |
| 17 | Positive | BrspLapur | *Brueelia* | *sp.* | *Lamprotornis purpureus* | Ghana | SRR23302922 |
| 18 | Positive | CuspCacas | *Cuculicola* | *sp.* | *Cacomantis castaneiventris* | Papua New Guinea | SRR5308216 |
| 19 | Positive | Cusp.Clcaf | *Cuculicola* | *sp.* | *Clamator jacobinus* | Malawi | SRR5308215 |
| 20 | Positive | BrspCipun | *Maculinirmus* | *sp.* | *Cinclosoma punctatum* | Australia | SRR8142481 |
| 21 | Positive | BrspDihot | *Dicruobates* | *sp.* | *Dicrurus hottentottus* | China | SRR14141081 |
| 22 | Positive | Cocln | *Columbicola* | *claytoni* | *Ducula rufigaster* | Papua New Guinea | SRR3161916 |
| 23 | Positive | Copar | *Columbicola* | *paradoxus* | *Lopholaimus antarcticus* | Australia | SRR3161944 |
| 24 | Positive | BrspPyjoc | *Brueelia* | *sp.* | *Pycnonotus jocosus* | China | SRR18820767 |
| 25 | Positive | BrspPyxan | *Brueelia* | *sp.* | *Pycnonotus xanthorrhous* | China | SRR18820765 |
| 26 | Positive | BrspHecas | *Brueelia* | *sp.* | *Hemixos castanonotus* | China | SRR14141075 |
| 27 | Positive | BrspOrchi | *Guimaraesiella* | *sp.* | *Oriolus chinensis* | China | SRR14141038 |

| Count_number | Sodalis_detection | Library_code | Louse_Genus | Louse_Species | Vertebrate_Host | Collection_Country | NCBI_SRA |
|---|---|---|---|---|---|---|---|
| 28 | Positive | BrspMenov | *Guimaraesiella* | *sp.* | *Menura novaehollandiae* | Australia | SRR14141061 |
| 29 | Positive | Piqua | *Picicola* | *quadrapustulosus* | *Pitta sordida* | Papua New Guinea | SRR5308242 |
| 30 | Positive | BrspUrery | *Olivinirmus* | *sp.* | *Urocissa erythroryncha* | China | SRR14141071 |
| 31 | Positive | PnspMeell | *Picicola* | *sp.* | *Dendropicos elliotii* | Cameroon | SRR8582582 |
| 32 | Positive | Bulon | *Buceronirmus* | *longicuneatus* | *Bycanistes bucinator* | Malawi | SRR5308203 |
| 33 | Positive | CuspCeaur | *Cuculicola* | *sp.* | *Ceuthmochares aereus* | Ghana | SRR5308217 |
| 34 | Positive | MaspComac | *Indoceoplanetes* | *sp.* | *Coracina macei* | China | SRR18820752 |
| 35 | Positive | MaspConov | *Indoceoplanetes* | *sp.* | *Coracina novaehollandiae* | Australia | SRR18820750 |
| 36 | Positive | Cobec | *Columbicola* | *beccari* | *Pampusana beccarii* | Solomon Islands | SRR3161941 |
| 37 | Positive | PispSmcap | *Picicola* | *sp.* | *Smithornis capensis* | Malawi | SRR5308248 |
| 38 | Positive | Brbio | *Hecatrishula* | *biocellata* | *Corvus corax* | Canada | SRR8142476 |
| 39 | Positive | Ffban | *Forficuloecus* | *banksi* | *Psephotellus varius* | Australia | SRR8175154 |
| 40 | Positive | Nsbor | *Neopsittaconirmus* | *borgiolii* | *Zanda funerea* | Australia | SRR8177120 |
| 41 | Positive | CbspEugla | *Capraiella* | *sp.* | *Eurystomus glaucurus* | Malawi | SRR5308205 |
| 42 | Positive | Corod | *Columbicola* | *rodmani* | *Geopelia humeralis* | Australia | SRR3161918 |
| 43 | Positive | Comal | *Columbicola* | *malenkeae* | *Ducula pacifica* | Vanuatu | SRR3161956 |
| 44 | Positive | BrspCyban | *Guimaraesiella* | *sp.* | *Cyornis banyumas* | China | SRR14141034 |
| 45 | Positive | BrspSafer | *Guimaraesiella* | *sp.* | *Saxicola ferreus* | China | SRR14141080 |
| 46 | Positive | GuspCoalb | *Guimaraesiella* | *sp.* | *Copsychus albospecularis* | Madagascar | SRR17556661 |
| 47 | Positive | Covei | *Columbicola* | *veigasimoni* | *Phapitreron leucotis* | Philippines | SRR3161919 |
| 48 | Positive | Cobcs | *Columbicola* | *bacillus 1* | *Streptopelia decaocto* | Netherlands | SRR3161950 |
| 49 | Positive | CospTucha | *Columbicola* | *sp.* | *Turtur chalcospilos* | Malawi | SRR3614998 |
| 50 | Positive | Picun | *Picicola* | *cunicularíae* | *Geositta cunicularia* | Bolivia | SRR5308237 |
| 51 | Positive | Teapa | *Trogoniella* | *apaloderma* | *Apaloderma narina* | Democratic Republic of Congo | SRR5308253 |
| 52 | Positive | Sglip | *Strongylocotes* | *lipogonus* | *Rhynchotus rufescens* | Bolivia | SRR5308142 |
| 53 | Positive | Sgtin | *Strongylocotes* | *tinami* | *Nothura boraquira* | Bolivia | SRR13159299 |
| 54 | Positive | FmspCocas | *Formicaphagus* | *sp.* | *Conopophaga castaneiceps* | Peru | SRR8334247 |
| 55 | Positive | Sglip2 | *Strongylocotes* | *lipogonus* | *Rhynchotus rufescens* | Argentina | SRR16503023 |
| 56 | Positive | Smste | *Saepocephalum* | *stephenfryi* | *Corcorax melanorhamphos* | Australia | SRR17556631 |
| 57 | Positive | BrspCasin | *Brueelia* | *sp.* | *Chloris sinica* | China | SRR8582560 |
| 58 | Positive | BrspPlvel | *Brueelia* | *sp.* | *Ploceus velatus* | South Africa | SRR18820723 |
| 59 | Positive | BrspPlmah | *Brueelia* | *sp.* | *Plocepasser mahali* | South Africa | SRR23302920 |
| 60 | Positive | GuspCycri | *Guimaraesiella* | *sp.* | *Cyanocitta cristata* | United States of America | SRR17556657 |
| 61 | Positive | NscirPlele | *Neopsittaconirmus* | *circumfasciatus* | *Platycercus elegans* | Australia | SRR8145987 |
| 62 | Positive | Rpgen | *Rhopaloceras* | *genitalis* | *Tinamus major* | Brazil | SRR16503024 |
| 63 | Positive | RpspTigut | *Rhopaloceras* | *sp.* | *Tinamus guttatus* | Brazil | SRR8146012 |
| 64 | Positive | Brcyc | *Brueelia* | *cyclothorax* | *Passer domesticus* | Canada | SRR23302892 |
| 65 | Positive | IscpsGrsqu | *Picicola* | *sp.* | *Grallaria squamigera* | Peru | SRR5308230 |

| Count_number | Sodalis_detection | Library_code | Louse_Genus | Louse_Species | Vertebrate_Host | Collection_Country | NCBI_SRA |
|---|---|---|---|---|---|---|---|
| 66 | Positive | FmspCoaur | *Formicaphagus* | *sp.* | *Conopophaga aurita* | Brazil | SRR8334246 |
| 67 | Positive | NscirAlchl | *Neopsittaconirmus* | *circumfasciatus* | *Alisterus chloropterus* | Papua New Guinea | SRR8177121 |
| 68 | Positive | NscirAlsca | *Neopsittaconirmus* | *circumfasciatus* | *Alisterus scapularis* | Australia | SRR8175170 |
| 69 | Positive | AfspColeu | *Alcedoffula* | *sp.* | *Corythornis leucogaster* | Ghana | SRR13159304 |
| 70 | Positive | OrspTimaj2 | *Ornicholax* | *sp.2* | *Tinamus major* | Brazil | SRR16503031 |
| 71 | Positive | ComeiCoarq | *Columbicola* | *meinertzhageni* | *Columba arquatrix* | Malawi | SRR16503057 |
| 72 | Positive | CospCosjo | *Columbicola* | *sp.* | *Columba sjostedti* | Cameroon | SRR8177103 |
| 73 | Positive | Comei | *Columbicola* | *sp.* | *Streptopelia semitorquata* | Ghana | SRR3161935 |
| 74 | Positive | Coclv | *Columbicola* | *claviformis* | *Columba palumbus* | United Kingdom | SRR3161920 |
| 75 | Positive | Cocla | *Columbicola* | *clayae* | *Treron waalia* | Ghana | SRR3161934 |
| 76 | Positive | PispPiiri | *Picicola* | *sp.* | *Pitta iris* | Australia | SRR5308250 |
| 77 | Positive | NsspCaban | *Neopsittaconirmus* | *sp.* | *Calyptorhynchus banksii* | Australia | SRR8177123 |
| 78 | Positive | BrspJuhye | *Brueelia* | *sp.* | *Junco hyemalis* | United States of America | SRR23302923 |
| 79 | Positive | BrspZoalb | *Brueelia* | *sp.* | *Zonotrichia albicollis* | Canada | SRR23302917 |
| 80 | Positive | BrspPaili | *Brueelia* | *sp.* | *Passerella iliaca* | Canada | SRR23302921 |
| 81 | Positive | BrspSicur | *Brueelia* | *sp.* | *Sialia currucoides* | United States of America | SRR18820763 |
| 82 | Positive | BrspEmpus | *Brueelia* | *sp.* | *Emberiza pusilla* | China | SRR14141077 |
| 83 | Positive | Brstr | *Brueelia* | *straminea* | *Dryobates pubescens* | Canada | SRR23302916 |
| 84 | Positive | Brana | *Brueelia* | *anamariae* | *Troglodytes aedon* | Canada | SRR23302925 |
| 85 | Positive | BrspCamex | *Brueelia* | *sp.* | *Haemorhous mexicanus* | United States of America | SRR18820735 |
| 86 | Positive | BrspCofla | *Brueelia* | *sp.* | *Coereba flaveola* | Panama | SRR23302879 |
| 87 | Positive | Brsplcgal | *Brueelia* | *sp.* | *Icterus galbula* | United States of America | SRR23302924 |
| 88 | Positive | Brced | *Brueelia* | *cedrorum* | *Bombycilla cedrorum* | Canada | SRR23302903 |
| 89 | Positive | Brili | *Brueelia* | *iliaci* | *Turdus migratorius* | Canada | SRR23302883 |
| 90 | Positive | Pifo | *Brueelia* | *sp.* | *Tyrannus forficatus* | United States of America | SRR5308239 |
| 91 | Positive | BrornAgpho | *Brueelia* | *ornatissima* | *Agelaius phoeniceus* | Canada | SRR23302881 |
| 92 | Positive | BrspFihyp | *Guimaraesiella* | *sp.* | *Ficedula hyperythra* | Philippines | SRR18820726 |
| 93 | Positive | BrspSIfro | *Guimaraesiella* | *sp.* | *Sitta frontalis* | Philippines | SRR18820762 |
| 94 | Positive | BrspRhnig | *Guimaraesiella* | *sp.* | *Rhipidura nigrocinnamomea* | Philippines | SRR18820764 |
| 95 | Positive | BrspFitri | *Guimaraesiella* | *sp.* | *Ficedula tricolor* | China | SRR14141076 |
| 96 | Positive | BrspTepar | *Guimaraesiella* | *sp.* | *Terpsiphone paradisi* | China | SRR14141035 |
| 97 | Positive | BrspPhino | *Guimaraesiella* | *sp.* | *Phylloscopus inornatus* | China | SRR18820724 |
| 98 | Positive | GuspPesig | *Guimaraesiella* | *sp.* | *Melanodryas sigillata* | New Guinea | SRR17556651 |
| 99 | Positive | Brsem | *Olivinirmus* | *semiannulata* | *Gymnorhina tibicen* | Australia | SRR8142480 |
| 100 | Positive | Olsem | *Olivinirmus* | *semiannulata* | *Strepera graculina* | Australia | SRR17556638 |
| 101 | Positive | OlsemCrarg | *Olivinirmus* | *semiannulata* | *Cracticus argenteus* | Australia | SRR18820747 |
| 102 | Positive | MaspComcg | *Indoceoplanetes* | *sp.* | *Malindangia mcgregori* | Philippines | SRR18820751 |
| 103 | Positive | MaspSpvir | *Maculinirmus* | *sp.* | *Sphecotheres viridis* | Australia | SRR18820748 |

| Count_number | Sodalis_detection | Library_code | Louse_Genus | Louse_Species | Vertebrate_Host | Collection_Country | NCBI_SRA |
|---|---|---|---|---|---|---|---|
| 104 | Positive | BrspCimag | *Guimaraesiella* | *sp.* | *Diphyllodes magnificus* | Papua New Guinea | SRR18820727 |
| 105 | Positive | Sgsub | *Strongylocotes* | *subspinosus* | *Nothocercus nigrocapillus* | Peru | SRR8582601 |
| 106 | Positive | StspNonig | *Strongylocotes* | *sp.* | *Nothocercus nigrocapillus* | Peru | SRR14908109 |
| 107 | Positive | SgspCrobs | *Strongylocotes* | *sp.* | *Crypturellus obsoletus* | Brazil | SRR13159333 |
| 108 | Positive | RespGamil | *Resartor* | *sp.* | *Trochalopteron milnei* | China | SRR8582595 |
| 109 | Positive | Afmad | *Alcedoffula* | *madagascariensis* | *Corythornis madagascariensis* | Madagascar | SRR13159307 |
| 110 | Positive | Qucas | *Quadraceps* | *caspius* | *Hydroprogne caspia* | Canada | SRR13389498 |
| 111 | Positive | AfspIspic | *Alcedoffula* | *sp.* | *Ispidina picta* | Democratic Republic of Congo | SRR13159301 |
| 112 | Positive | Coexi4 | *Columbicola* | *exilicornis 4* | *Macropygia amboinensis* | Papua New Guinea | SRR3161945 |
| 113 | Positive | Cogui3 | *Columbicola* | *guimaraesi 3* | *Chalcophaps stephani* | Papua New Guinea | SRR3161927 |
| 114 | Positive | Cogui4 | *Columbicola* | *guimaraesi 4* | *Chalcophaps indica* | China | SRR16503016 |
| 115 | Positive | Coeme1 | *Columbicola* | *emersoni 1* | *Ptilinopus superbus* | Australia | SRR8145886 |
| 116 | Positive | BrspLelut | *Guimaraesiella* | *sp.* | *Leiothrix lutea* | China | SRR8582561 |
| 117 | Positive | Cosmi | *Columbicola* | *smithae* | *Turtur brehmeri* | Ghana | SRR3161936 |
| 118 | Positive | AfspCeazu | *Alcedoffula* | *sp.* | *Ceyx azureus* | Australia | SRR13159305 |
| 119 | Positive | CocolColiv | *Columbicola* | *columbae* | *Columba livia* | United States of America | SRR8177102 |
| 120 | Positive | Cotsc | *Columbicola* | *tschulyschman* | *Columba livia* | United States of America | SRR3161959 |
| 121 | Positive | Ratay | *Rallicola* | *taylori* | *Fulica gigantea* | Argentina | SRR8173303 |
| 122 | Positive | Quimp | *Quadraceps* | *impar* | *Tringa brevipes* | Australia | SRR13159321 |
| 123 | Positive | AncraAngra | *Anaticola* | *crassicornis* | *Anas gracilis* | Australia | SRR13389464 |
| 124 | Positive | AncraAnund | *Anaticola* | *crassicornis* | *Anas undulata* | South Africa | SRR13389501 |
| 125 | Positive | AncraAnpla | *Anaticola* | *crassicornis* | *Anas platyrhynchos* | United States of America | SRR14141048 |
| 126 | Positive | AncraAnfla | *Anaticola* | *crassicornis* | *Anas flavirostris* | Argentina | SRR13389475 |
| 127 | Positive | AnmerMamem | *Anaticola* | *mergiserrati* | *Malacorhynchus membranaceus* | Australia | SRR14141042 |
| 128 | Positive | AnspChmel | *Anaticola* | *sp.* | *Oressochen melanopterus* | Bolivia | SRR16574572 |
| 129 | Positive | Qusim | *Quadraceps* | *similis* | *Tringa nebularia* | Italy | SRR18820731 |
| 130 | Positive | RaspIrgal | *Rallicola* | *sp.* | *Irediparra gallinacea* | Australia | SRR5308188 |
| 131 | Positive | QuspStnil | *Quadraceps* | *sp.* | *Gelochelidon nilotica* | Australia | SRR8146006 |
| 132 | Positive | Quhos | *Quadraceps* | *hospes* | *Pluvialis squatarola* | Japan | SRR18820737 |
| 133 | Positive | Quumb | *Quadraceps* | *umbrinus* | *Scopus umbretta* | Malawi | SRR8566334 |
| 134 | Positive | Cachi | *Caracaricola* | *chimangophilus* | *Daptrius chimango* | Argentina | SRR5308152 |
| 135 | Positive | Plrob | *Philoceanus* | *robertsi* | *Oceanites oceanicus* | British Antarctic Territory | SRR5308176 |
| 136 | Positive | Aratr | *Ardeicola* | *atratus* | *Nycticorax caledonicus* | Australia | SRR17556671 |
| 137 | Positive | ArspIxfla | *Ardeicola* | *sp.* | *Botaurus flavicollis* | Australia | SRR16574611 |
| 138 | Positive | Racen | *Rallicola* | *centropus* | *Centropus superciliosus* | Malawi | SRR8177136 |
| 139 | Positive | RaspCesen | *Rallicola* | *sp.* | *Centropus senegalensis* | Ghana | SRR5308187 |
| 140 | Positive | Agwat | *Austrogoniodes* | *waterstoni* | *Eudyptula minor* | Australia | SRR5308390 |
| 141 | Positive | Ibfla | *Ibidoecus* | *flavus* | *Platalea flavipes* | Australia | SRR8177118 |

| Count_number | Sodalis_detection | Library_code | Louse_Genus | Louse_Species | Vertebrate_Host | Collection_Country | NCBI_SRA |
|---|---|---|---|---|---|---|---|
| 142 | Positive | CuspScnov | *Cuculicola* | *sp.* | *Scythrops novaehollandiae* | Australia | SRR5308219 |
| 143 | Positive | Costo | *Cotingacola* | *stotzi* | *Querula purpurata* | Brazil | SRR5308212 |
| 144 | Positive | Rakel | *Rallicola* | *kelloggi* | *Rallus limicola* | United States of America | SRR8177138 |
| 145 | Positive | Comac4 | *Columbicola* | *macrourae 4* | *Zenaida galapagoensis* | Ecuador | SRR8145889 |
| 146 | Positive | CospZeaur | *Columbicola* | *sp.* | *Zenaida auriculata* | Ecuador | SRR16503053 |
| 147 | Positive | CospZemel | *Columbicola* | *macrourae 6* | *Zenaida meloda* | Peru | SRR3161971 |
| 148 | Positive | CbspEugul | *Capraiella* | *sp.* | *Eurystomus gularis* | Ghana | SRR17556626 |
| 149 | Positive | NsspApery | *Neopsittaconirmus* | *sp.* | *Aprosmictus erythropterus* | Australia | SRR8175172 |
| 150 | Positive | PnspCacai | *Picicola* | *sp.* | *Campethera maculosa* | Malawi | SRR8566321 |
| 151 | Positive | Arneo | *Ardeicola* | *neopallidus* | *Egretta sacra* | Australia | SRR17556648 |
| 152 | Positive | BmspTonas | *Buceroemersonia* | *sp.* | *Lophoceros nasutus* | Ghana | SRR17556627 |
| 153 | Positive | Bclat | *Bucerocophorus* | *latifrons* | *Lophoceros pallidirostris* | Malawi | SRR5308201 |
| 154 | Positive | CospCeorn | *Cotingacola* | *sp.* | *Cephalopterus ornatus* | Peru | SRR5308211 |
| 155 | Positive | CbspCoaby | *Capraiella* | *sp.* | *Coracias abyssinicus* | Ghana | SRR5308204 |
| 156 | Positive | Piser | *Picicola* | *serrafreirei* | *Nystalus chacuru* | Bolivia | SRR5308243 |
| 157 | Positive | Cobcs2 | *Columbicola* | *bacilus 2* | *Streptopelia decipiens* | Uganda | SRR3161967 |
| 158 | Positive | CkspNodar2 | *Cuclotocephalus* | *sp.* | *Nothura darwinii* | Peru | SRR16503038 |
| 159 | Positive | CkspNopen | *Cuclotocephalus* | *secundus* | *Nothoprocta pentlandii* | Argentina | SRR14908140 |
| 160 | Positive | Dgdis | *Degeeriella* | *discocephalus* | *Haliaeetus leucocephalus* | Canada | SRR5308222 |
| 161 | Positive | Gochr | *Goniocotes* | *chrysocephalus* | *Phasianus colchicus* | United States of America | SRR5308160 |
| 162 | Positive | Afcri | *Alcedoffula* | *cristata* | *Corythornis cristatus* | Madagascar | SRR13159312 |
| 163 | Positive | Comjo3 | *Columbicola* | *mjoebergi 3* | *Geopelia placida* | Australia | SRR3161942 |
| 164 | Positive | Cuato | *Cuculicola* | *atopus* | *Piaya cayana* | Peru | SRR5308214 |
| 165 | Positive | Pinao | *Picicola* | *naokii* | *Bucco capensis* | Peru | SRR5308240 |
| 166 | Positive | TuspTrcal | *Turturicola* | *sp.* | *Treron calvus* | Malawi | SRR5308257 |
| 167 | Positive | Dgvag | *Degeeriella* | *vagans* | *Accipiter gentilis* | Canada | SRR5308226 |
| 168 | Positive | Comck | *Columbicola* | *mckeani* | *Ocyphaps lophotes* | Australia | SRR3161929 |
| 169 | Positive | PispPifla | *Picicola* | *sp.* | *Piculus flavigula* | Brazil | SRR5308246 |
| 170 | Positive | CospStori | *Columbicola* | *sp.* | *Streptopelia orientalis* | China | SRR3161951 |
| 171 | Positive | Picam | *Picicola* | *campethera* | *Campethera bennettii* | Malawi | SRR5308236 |
| 172 | Positive | GuspTrmel | *Guimaraesiella* | *sp.* | *Trogon melanurus* | Panama | SRR17556644 |
| 173 | Positive | Trhas | *Trogoninirmus* | *hastiformis* | *Trogon melanocephalus* | Mexico | SRR5308254 |
| 174 | Positive | PispGatom | *Picicola* | *sp.* | *Galbula tombacea* | Peru | SRR5308249 |
| 175 | Positive | Pifau | *Picicola* | *faucetti* | *Chelidoptera tenebrosa* | Brazil | SRR5308238 |
| 176 | Positive | Auand | *Austrophilopterus* | *andigenae* | *Andigena hypoglauca* | Peru | SRR5308196 |
| 177 | Positive | Dgruf1 | *Degeeriella* | *rufa* | *Falco berigora* | Australia | SRR16574599 |
| 178 | Positive | Arnig | *Ardeicola* | *nigra* | *Butorides striata* | Australia | SRR17556637 |
| 179 | Positive | ComasB | *Columbicola* | *masoni 2* | *Petrophassa rufipennis* | Australia | SRR3161946 |

| Count_number | Sodalis_detection | Library_code | Louse_Genus | Louse_Species | Vertebrate_Host | Collection_Country | NCBI_SRA |
|---|---|---|---|---|---|---|---|
| 180 | Positive | AuspAupra | *Austrophilopterus* | *sp.* | *Aulacorhynchus prasinus* | Peru | SRR5308199 |
| 181 | Positive | Ckext | *Cuclotocephalus* | *extraneus* | *Nothoprocta curvirostris* | Peru | SRR14908120 |
| 182 | Positive | CkspNoorn | *Cuclotocephalus* | *sp.* | *Nothoprocta ornata* | Bolivia | SRR8175153 |
| 183 | Positive | DgspBupla | *Degeeriella* | *sp.* | *Buteo platypterus* | Canada | SRR5308225 |
| 184 | Positive | DgrufFaper | *Degeeriella* | *rufa* | *Falco peregrinus* | Canada | SRR5308223 |
| 185 | Positive | Ffcam | *Forficuloecus* | *cameroni* | *Aprosmictus erythropterus* | Australia | SRR8175155 |
| 186 | Positive | Popap | *Podargoecus* | *papuensis* | *Podargus papuensis* | Australia | SRR5308376 |
| 187 | Positive | Coang | *Columbicola* | *tasmaniensis* | *Phaps chalcoptera* | Australia | SRR3161947 |
| 188 | Positive | Cotas | *Columbicola* | *tasmaniensis* | *Phaps elegans* | Australia | SRR3161948 |
| 189 | Positive | Coang07 | *Columbicola* | *angustus* | *Phaps chalcoptera* | Australia | SRR8145885 |
| 190 | Positive | Cohar | *Columbicola* | *harbisoni* | *Phaps histrionica* | Australia | SRR3161949 |
| 191 | Positive | BispCacel | *Bizarrifrons* | *sp.* | *Cacicus cela* | Panama | SRR5308150 |
| 192 | Positive | NsspNebou | *Neopsittaconirmus* | *sp.* | *Neopsephotus bourkii* | Australia | SRR8177124 |
| 193 | Positive | Codro | *Columbicola* | *drowni* | *Metriopelia melanoptera* | Argentina | SRR3161922 |
| 194 | Positive | Cogym | *Columbicola* | *gymnopeliae* | *Metriopelia ceciliae* | Peru | SRR3161923 |
| 195 | Positive | Copas1 | *Columbicola* | *passerinae 1* | *Columbina picui* | Argentina | SRR3161931 |
| 196 | Positive | Copas2 | *Columbicola* | *passerinae 2* | *Columbina cruziana* | Peru | SRR3161930 |
| 197 | Positive | Fnpar | *Furnariphilus* | *parkeri* | *Sclerurus caudacutus* | Brazil | SRR8334254 |
| 198 | Positive | Pepun | *Pectenosoma* | *punensis* | *Crypturellus obsoletus* | Peru | SRR14908130 |
| 199 | Positive | KespCrstr | *Kelloggia* | *sp.* | *Crypturellus strigulosus* | Brazil | SRR5308165 |
| 200 | Positive | Piosh | *Picicola* | *osheai* | *Malacoptila fusca* | Surinam | SRR5308241 |
| 201 | Positive | Coext | *Columbicola* | *extinctus* | *Patagioenas fasciata* | United States of America | SRR3161924 |
| 202 | Positive | Coext2 | *Columbicola* | *extinctus 2* | *Patagioenas fasciata* | Peru | SRR8145888 |
| 203 | Positive | QsspCrsou | *Pseudolipeurus* | *sp.* | *Crypturellus soui* | Brazil | SRR14908116 |
| 204 | Positive | SgspCrsou | *Strongylocotes* | *sp.* | *Crypturellus soui* | Brazil | SRR8582600 |
| 205 | Positive | SgspCrsou2 | *Strongylocotes* | *sp.* | *Crypturellus soui* | Brazil | SRR14908110 |
| 206 | Positive | FospChnob373 | *Formicaricola* | *sp.* | *Chamaeza nobilis* | Brazil | SRR8582572 |
| 207 | Positive | Memul | *Megapeostus* | *multiplex* | *Crypturellus boucardi* | Mexico | SRR1821940 |
| 208 | Positive | SgspCrobsPE | *Strongylocotes* | *sp.* | *Crypturellus obsoletus* | Peru | SRR14908111 |
| 209 | Positive | OxspOdspe | *Oxylipeurus* | *sp.* | *Odontophorus speciosus* | Peru | SRR16574590 |
| 210 | Positive | MpspCrcin2 | *Megapeostus* | *sp. 2* | *Crypturellus cinnamomeus* | Mexico | SRR13159341 |
| 211 | Positive | Pssim | *Pseudolipeurus* | *similis* | *Crypturellus cinnamomeus* | Mexico | SRR13159297 |
| 212 | Positive | PnspCeele | *Picicola* | *sp.* | *Celeus elegans* | Peru | SRR9693802 |
| 213 | Positive | Tiele | *Tinamotaecola* | *elegans* | *Eudromia elegans* | Argentina | SRR5308366 |
| 214 | Positive | TrspTrper | *Trogoninirmus* | *sp.* | *Trogon personatus* | Guyana | SRR5308255 |
| 215 | Positive | Brbru | *Guimaraesiella* | *brunneinucha* | *Dumetella carolinensis* | Canada | SRR14141057 |
| 216 | Positive | SnspHymus | *Sturnidoecus* | *sp.* | *Hylocichla mustelina* | Panama | SRR14141055 |
| 217 | Positive | PsspCrstr | *Pseudolipeurus* | *sp.* | *Crypturellus strigulosus* | Brazil | SRR12762535 |

| Count_number | Sodalis_detection | Library_code | Louse_Genus | Louse_Species | Vertebrate_Host | Collection_Country | NCBI_SRA |
|---|---|---|---|---|---|---|---|
| 218 | Positive | Qsobs2 | *Pseudophilopterus* | *obseletus* | *Crypturellus obsoletus* | Peru | SRR14908117 |
| 219 | Positive | Sawum | *Saemundssonia* | *wumisuzume* | *Aethia cristatella* | United States of America | SRR8146014 |
| 220 | Positive | Sawum2 | *Saemundssonia* | *wumisuzume* | *Aethia cristatella* | United States of America | SRR13159311 |
| 221 | Positive | Qeant | *Pseudocophorus* | *antennatus* | *Pipreola arcuata* | Peru | SRR5308251 |
| 222 | Positive | Pzdio | *Paraclisis* | *diomedeae* | *Phoebetria palpebrata* | United Kingdom | SRR5308183 |
| 223 | Positive | QuspRenov2 | *Quadraceps* | *sp. 2* | *Recurvirostra novaehollandiae* | Australia | SRR13159327 |
| 224 | Positive | Zites | *Cirropthirius* | *testudinarius* | *Recurvirostra americana* | United States of America | SRR5308194 |
| 225 | Positive | MuspCaena | *Mulcticola* | *sp.* | *Gactornis enarratus* | Madagascar | SRR8145979 |
| 226 | Positive | Quaet | *Quadraceps* | *aetherus* | *Aethia cristatella* | United States of America | SRR8173278 |
| 227 | Positive | Quaet2 | *Quadraceps* | *aetherus* | *Aethia pusilla* | United States of America | SRR13159308 |
| 228 | Positive | Quaet1 | *Quadraceps* | *aetherus* | *Aethia cristatella* | United States of America | SRR13159310 |
| 229 | Negative | Adful | *Acidoproctus* | *fuligulae* | *Netta peposaca* | Argentina | SRR16574618 |
| 230 | Negative | Adhil | *Acidoproctus* | *hilli* | *Anseranas semipalmata* | Australia | SRR5809354 |
| 231 | Negative | Adros | *Acidoproctus* | *rostratus* | *Dendrocygna viduata* | Argentina | SRR5308389 |
| 232 | Negative | Brlon | *Acronirmus* | *longus* | *Tachycineta bicolor* | Canada | SRR8142477 |
| 233 | Negative | Acmex | *Acutifrons* | *mexicanus* | *Caracara plancus* | United States of America | SRR5308195 |
| 234 | Negative | Aldel | *Alcedoecus* | *delphax* | *Dacelo novaeguineae* | Australia | SRR8172493 |
| 235 | Negative | AlspHabad | *Alcedoecus* | *sp.* | *Halcyon badia* | Ghana | SRR5308110 |
| 236 | Negative | Afalc | *Alcedoffula* | *alcyonae* | *Megaceryle alcyon* | Canada | SRR5308368 |
| 237 | Negative | Afcey | *Alcedoffula* | *ceycis* | *Ceyx erithaca* | Malaysia | SRR8172492 |
| 238 | Negative | Afcho | *Alcedoffula* | *chocoana* | *Chloroceryle inda* | Peru | SRR13159306 |
| 239 | Negative | AfspChama | *Alcedoffula* | *sp.* | *Chloroceryle amazona* | Panama | SRR13159303 |
| 240 | Negative | AfspChame | *Alcedoffula* | *sp.* | *Chloroceryle americana* | Panama | SRR13159302 |
| 241 | Negative | AnansChcae | *Anaticola* | *anseris* | *Anser caerulescens* | Canada | SRR16574606 |
| 242 | Negative | AnansBrcan | *Anaticola* | *anseris* | *Branta canadensis* | Canada | SRR16574617 |
| 243 | Negative | AnansBrhut | *Anaticola* | *anseris* | *Branta hutchinsii* | Canada | SRR14141039 |
| 244 | Negative | Anasy | *Anaticola* | *asymmetricus* | *Alopochen aegyptiaca* | South Africa | SRR13389486 |
| 245 | Negative | Anaus | *Anaticola* | *australis* | *Cereopsis novaehollandiae* | Australia | SRR16574595 |
| 246 | Negative | AncraOxvit | *Anaticola* | *crassicornis* | *Oxyura vittata* | Argentina | SRR14141046 |
| 247 | Negative | Anjam | *Anaticola* | *jamesi* | *Phoenicoparrus jamesi* | Argentina | SRR13389492 |
| 248 | Negative | Anmag | *Anaticola* | *magnificus* | *Tadorna tadornoides* | Australia | SRR14141040 |
| 249 | Negative | Anmar | *Anaticola* | *marginella* | *Chloephaga picta* | Falkland Islands | SRR16574584 |
| 250 | Negative | AnmerAispo | *Anaticola* | *mergiserrati* | *Aix sponsa* | Canada | SRR16574575 |
| 251 | Negative | AnmerAyaus | *Anaticola* | *mergiserrati* | *Aythya australis* | Australia | SRR14141044 |
| 252 | Negative | AnmerAymar | *Anaticola* | *mergiserrati* | *Aythya marila* | United States of America | SRR14141045 |
| 253 | Negative | AnmerHihis | *Anaticola* | *mergiserrati* | *Histrionicus histrionicus* | United States of America | SRR16574574 |
| 254 | Negative | Anpho | *Anaticola* | *phoenicopteri* | *Phoenicopterus chilensis* | Argentina | SRR5308382 |
| 255 | Negative | AnphoPhrub | *Anaticola* | *phoenicopteri* | *Phoenicopterus ruber* | Italy | SRR13389491 |

| Count_number | Sodalis_detection | Library_code | Louse_Genus | Louse_Species | Vertebrate_Host | Collection_Country | NCBI_SRA |
|---|---|---|---|---|---|---|---|
| 256 | Negative | Anrhe | *Anaticola* | *rheinwaldi* | *Branta bernicla* | Sweden | SRR16574573 |
| 257 | Negative | AnspBilob | *Anaticola* | *sp.* | *Biziura lobata* | Australia | SRR13389490 |
| 258 | Negative | AnspBualb | *Anaticola* | *sp.* | *Bucephala albeola* | Canada | SRR16503061 |
| 259 | Negative | AnspCaleu | *Anaticola* | *sp.* | *Callonetta leucophrys* | Argentina | SRR14141041 |
| 260 | Negative | AnspChjub | *Anaticola* | *sp.* | *Chenonetta jubata* | Australia | SRR14141052 |
| 261 | Negative | AnspPhand | *Anaticola* | *sp.* | *Phoenicoparrus andinus* | Argentina | SRR13389489 |
| 262 | Negative | AnspMearm | *Anaticola* | *sp.* | *Merganetta armata* | Peru | SRR14141043 |
| 263 | Negative | Antho | *Anaticola* | *thoracicus* | *Radjah radjah* | Australia | SRR16574571 |
| 264 | Negative | AtdenBrcan | *Anatoecus* | *dentatus* | *Branta canadensis* | United States of America | SRR16574610 |
| 265 | Negative | AtictAyaff | *Anatoecus* | *icterodes* | *Aythya affinis* | United States of America | SRR16574608 |
| 266 | Negative | Atict | *Anatoecus* | *icterodes* | *Spatula cyanoptera* | United States of America | SRR5308111 |
| 267 | Negative | Atict1 | *Anatoecus* | *icterodes* | *Spatula cyanoptera* | United States of America | SRR16574609 |
| 268 | Negative | AtkeyPhjam | *Anatoecus* | *keymeri* | *Phoenicoparrus jamesi* | Argentina | SRR13389484 |
| 269 | Negative | Atkey | *Anatoecus* | *keymeri* | *Phoenicopterus chilensis* | Argentina | SRR5308381 |
| 270 | Negative | Atpen | *Anatoecus* | *penicillatus* | *Cygnus olor* | Sweden | SRR16503060 |
| 271 | Negative | AtspChpic | *Anatoecus* | *sp.* | *Chloephaga picta* | Argentina | SRR17556628 |
| 272 | Negative | AtspCyatr | *Anatoecus* | *sp.* | *Cygnus atratus* | Australia | SRR13389483 |
| 273 | Negative | AtspPhand | *Anatoecus* | *sp.* | *Phoenicoparrus andinus* | Argentina | SRR13389482 |
| 274 | Negative | Aqocc | *Aquanirmus* | *occidentalis* | *Aechmophorus occidentalis* | Canada | SRR5308392 |
| 275 | Negative | AqpodPocri | *Aquanirmus* | *podicepis* | *Podiceps cristatus* | Australia | SRR16574616 |
| 276 | Negative | Aqpod | *Aquanirmus* | *podilymbus* | *Podilymbus podiceps* | Canada | SRR8177097 |
| 277 | Negative | Aqrol | *Aquanirmus* | *rollandii* | *Rollandia rolland* | Argentina | SRR13389488 |
| 278 | Negative | AqspPopol | *Aquanirmus* | *sp.* | *Poliocephalus poliocephalus* | Australia | SRR16574615 |
| 279 | Negative | AqspTanov | *Aquanirmus* | *sp.* | *Tachybaptus novaehollandiae* | Australia | SRR16574614 |
| 280 | Negative | Araus | *Ardeicola* | *australis* | *Threskiornis spinicollis* | Australia | SRR17556670 |
| 281 | Negative | Arexp | *Ardeicola* | *expallidus* | *Bubulcus ibis* | United States of America | SRR5308391 |
| 282 | Negative | ArexpEggar | *Ardeicola* | *expallidus* | *Egretta garzetta* | Australia | SRR16574613 |
| 283 | Negative | Arger | *Ardeicola* | *geronticorum* | *Geronticus calvus* | Morocco | SRR17556659 |
| 284 | Negative | Arhar | *Ardeicola* | *harrisoni* | *Platalea flavipes* | Australia | SRR8177098 |
| 285 | Negative | Aribi | *Ardeicola* | *ibis* | *Threskiornis molucca* | Australia | SRR13389485 |
| 286 | Negative | Arpil | *Ardeicola* | *pilgrimi* | *Egretta novaehollandiae* | Australia | SRR16574612 |
| 287 | Negative | Arrha2 | *Ardeicola* | *rhaphidius* | *Plegadis chihi* | United States of America | SRR5308147 |
| 288 | Negative | ArrhaPlfal | *Ardeicola* | *rhaphidius* | *Plegadis falcinellus* | Australia | SRR17556630 |
| 289 | Negative | ArspAribi | *Ardeicola* | *sp.* | *Bubulcus ibis* | Australia | SRR17556629 |
| 290 | Negative | Aacoc | *Ardeiphagus* | *cochlearius* | *Cochlearius cochlearius* | Brazil | SRR5308384 |
| 291 | Negative | Aasim | *Ardeiphagus* | *similis* | *Tigrisoma lineatum* | Brazil | SRR13389487 |
| 292 | Negative | Auaff | *Auricotes* | *affinis* | *Ducula rufigaster* | Papua New Guinea | SRR8172496 |
| 293 | Negative | AuspPtriv | *Auricotes* | *bellus* | *Ptilinopus rivoli* | Papua New Guinea | SRR8172498 |

| Count_number | Sodalis_detection | Library_code | Louse_Genus | Louse_Species | Vertebrate_Host | Collection_Country | NCBI_SRA |
|---|---|---|---|---|---|---|---|
| 294 | Negative | Aurot | *Auricotes* | *rotundus* | *Ptilinopus occipitalis* | Philippines | SRR18820769 |
| 295 | Negative | AuspDubak | *Auricotes* | *sp.* | *Ducula bakeri* | Vanuatu | SRR18820768 |
| 296 | Negative | AuspDubic | *Auricotes* | *sp.* | *Ducula bicolor* | Australia | SRR5308148 |
| 297 | Negative | AuspDupac | *Auricotes* | *sp.* | *Ducula pacifica* | Vanuatu | SRR8172497 |
| 298 | Negative | Auint | *Austrokelloggia* | *intermedia* | *Nothocercus nigrocapillus* | Peru | SRR14908121 |
| 299 | Negative | Apcan | *Austrophilopterus* | *cancellosus* | *Ramphastos sulfuratus* | Panama | SRR5308369 |
| 300 | Negative | Aucan | *Austrophilopterus* | *cancellosus* | *Ramphastos tucanus* | Brazil | SRR5308197 |
| 301 | Negative | Aufla | *Austrophilopterus* | *flavirostris* | *Pteroglossus aracari aracari* | Brazil | SRR5308198 |
| 302 | Negative | Beuni | *Bedfordiella* | *unica* | *Aphrodroma brevirostris* | Kerguelan Island | SRR5308149 |
| 303 | Negative | Btmac1 | *Bothriometopus* | *macrocnemis* | *Chauna torquata* | Argentina | SRR16574603 |
| 304 | Negative | Brbr | *Brueelia* | *brachythorax* | *Bombycilla garrulus* | Ukraine | SRR16503049 |
| 305 | Negative | Brdef | *Brueelia* | *deficiens* | *Aphelocoma californica* | United States of America | SRR18820757 |
| 306 | Negative | BrspPamel | *Brueelia* | *sp.* | *Passer melanurus* | South Africa | SRR8177100 |
| 307 | Negative | Bmcla | *Buceroemersonia* | *clarkei* | *Lophoceros pallidirostris* | Malawi | SRR5308202 |
| 308 | Negative | BlspBrlep | *Buerelius* | *sp.* | *Brachypteracias leptosomus* | Madagascar | SRR5308151 |
| 309 | Negative | Cabid | *Campanulotes* | *bidentatus* | *Columba palumbus* | United Kingdom | SRR8172500 |
| 310 | Negative | Cacam5337 | *Campanulotes* | *campanulatus* | *Nesoenas picturatus* | Madagascar | SRR8172501 |
| 311 | Negative | Cacom | *Campanulotes* | *compar* | *Columba livia* | United States of America | SRR1821983 |
| 312 | Negative | Cacom1 | *Campanulotes* | *compar* | *Columba livia* | United States of America | SRR12762520 |
| 313 | Negative | Cadur | *Campanulotes* | *durdeni* | *Ocyphaps lophotes* | Australia | SRR8172502 |
| 314 | Negative | Caele | *Campanulotes* | *elegans* | *Phaps elegans* | Australia | SRR8145855 |
| 315 | Negative | Cafla4 | *Campanulotes* | *flavus* | *Leucosarcia melanoleuca* | Australia | SRR8172504 |
| 316 | Negative | Cafla3 | *Campanulotes* | *flavus* | *Phaps chalcoptera* | Australia | SRR8172503 |
| 317 | Negative | Cafre | *Campanulotes* | *frenatus* | *Zentrygon frenata* | Peru | SRR8172505 |
| 318 | Negative | CaspGehum | *Campanulotes* | *sp.* | *Geopelia humeralis* | Australia | SRR8172507 |
| 319 | Negative | CaspGeplu | *Campanulotes* | *sp.* | *Geophaps plumifera* | Australia | SRR8172508 |
| 320 | Negative | CaspGesmi | *Campanulotes* | *sp.* | *Geophaps smithii* | Australia | SRR8145857 |
| 321 | Negative | Cdlap | *Carduiceps* | *laponicus* | *Limosa lapponica* | Japan | SRR18820759 |
| 322 | Negative | Cdzon | *Carduiceps* | *zonarius* | *Calidris fuscicollis* | Brazil | SRR5308206 |
| 323 | Negative | PyspOdgut | *Chelopistes* | *sp.* | *Odontophorus guttatus* | Panama | SRR5308181 |
| 324 | Negative | Chtex | *Chelopistes* | *texanus* | *Ortalis vetula* | United States of America | SRR5308114 |
| 325 | Negative | Chtex1 | *Chelopistes* | *texanus* | *Ortalis vetula* | United States of America | SRR16574602 |
| 326 | Negative | CispCileu | *Cincloecus* | *neotropicalis* | *Cinclus leucocephalus* | Peru | SRR14887915 |
| 327 | Negative | ClspMomo | *Clayiella* | *sp.* | *Momotus momota* | Brazil | SRR14887914 |
| 328 | Negative | Cpcol | *Colilipeurus* | *colius* | *Urocolius indicus* | South Africa | SRR5308386 |
| 329 | Negative | Cpobs | *Colilipeurus* | *obscurior* | *Colius colius* | South Africa | SRR5308370 |
| 330 | Negative | Cpobs2 | *Colilipeurus* | *obscurior* | *Colius colius* | South Africa | SRR16574601 |
| 331 | Negative | Cxdoc | *Colinicola* | *docophoroides* | *Callipepla californica* | United States of America | SRR5308220 |

| Count_number | Sodalis_detection | Library_code | Louse_Genus | Louse_Species | Vertebrate_Host | Collection_Country | NCBI_SRA |
|---|---|---|---|---|---|---|---|
| 332 | Negative | Cxmea | *Colinicola* | *mearnsi* | *Cyrtonyx montezumae* | United States of America | SRR5308221 |
| 333 | Negative | Cccas | *Coloceras* | *castroi* | *Turtur tympanistria* | Democratic Republic of Congo | SRR8172510 |
| 334 | Negative | CcspStcap | *Coloceras* | *chinense* | *Streptopelia capicola* | Kenya | SRR8145873 |
| 335 | Negative | Ccchi | *Coloceras* | *chinense* | *Cobcs* | United States of America | SRR8145858 |
| 336 | Negative | Cccla | *Coloceras* | *clayae* | *Aplopelia larvata* | Malawi | SRR8172499 |
| 337 | Negative | Cccly | *Coloceras* | *clypeatus* | *Phapitreron amethystinus* | Philippines | SRR18820761 |
| 338 | Negative | Ccdam | *Coloceras* | *damicorne* | *Columba palumbus* | United Kingdom | SRR8145859 |
| 339 | Negative | CcdorMaamb | *Coloceras* | *doryanus* | *Macropygia amboinensis* | Australia | SRR8145860 |
| 340 | Negative | CcdorManig | *Coloceras* | *doryanus* | *Macropygia nigrirostris* | Papua New Guinea | SRR8145861 |
| 341 | Negative | Ccfur | *Coloceras* | *furcatum* | *Lopholaimus antarcticus* | Australia | SRR8145862 |
| 342 | Negative | Ccgra | *Coloceras* | *grande* | *Phaps chalcoptera* | Australia | SRR8145863 |
| 343 | Negative | Cchil | *Coloceras* | *hilli* | *Streptopelia decaocto* | United States of America | SRR8172511 |
| 344 | Negative | CcspStpic | *Coloceras* | *hoogstrali* | *Nesoenas picturatus* | Madagascar | SRR8145875 |
| 345 | Negative | Cclat | *Coloceras* | *laticlypeatus* | *Turtur brehmeri* | Ghana | SRR8145864 |
| 346 | Negative | Ccmus | *Coloceras* | *museihalense* | *Reinwardtoena reinwardti* | Papua New Guinea | SRR8145865 |
| 347 | Negative | Ccset | *Coloceras* | *setosum* | *Treron waalia* | Ghana | SRR8172513 |
| 348 | Negative | CcspAplar | *Coloceras* | *sp.* | *Aplopelia larvata* | Malawi | SRR8145866 |
| 349 | Negative | CcspChind | *Coloceras* | *sp.* | *Chalcophaps indica* | Vanuatu | SRR8172514 |
| 350 | Negative | Cahet | *Coloceras* | *sp.* | *Columba leucomela* | Australia | SRR8145856 |
| 351 | Negative | CaspColcm | *Coloceras* | *sp.* | *Columba leucomela* | Australia | SRR8172506 |
| 352 | Negative | CcspCosjo | *Coloceras* | *sp.* | *Columba sjostedti* | Cameroon | SRR8177101 |
| 353 | Negative | CcspGecun | *Coloceras* | *sp.* | *Geopelia cuneata* | Australia | SRR8172515 |
| 354 | Negative | CcspGehum | *Coloceras* | *sp.* | *Geopelia humeralis* | Australia | SRR8145867 |
| 355 | Negative | CcspGepla | *Coloceras* | *sp.* | *Geopelia placida* | Australia | SRR8145868 |
| 356 | Negative | CcspGestr | *Coloceras* | *sp.* | *Geopelia striata* | United States of America | SRR9693826 |
| 357 | Negative | CcspHenov | *Coloceras* | *sp.* | *Hemiphaga novaeseelandiae* | New Zealand | SRR18820760 |
| 358 | Negative | CcspLemel | *Coloceras* | *sp.* | *Leucosarcia melanoleuca* | Australia | SRR8145869 |
| 359 | Negative | CcspMamac | *Coloceras* | *sp.* | *Macropygia mackinlayi* | Vanuatu | SRR8172516 |
| 360 | Negative | CcpsOclop | *Coloceras* | *sp.* | *Ocyphaps lophotes* | Australia | SRR8172512 |
| 361 | Negative | CcspStlug | *Coloceras* | *sp.* | *Streptopelia lugens* | Malawi | SRR8145874 |
| 362 | Negative | CcspStsem | *Coloceras* | *sp.* | *Streptopelia semitorquata* | Ghana | SRR8145876 |
| 363 | Negative | CcspTucha | *Coloceras* | *sp.* | *Turtur chalcospilos* | Malawi | SRR8145877 |
| 364 | Negative | Ccste | *Coloceras* | *stephani* | *Chalcophaps stephani* | Papua New Guinea | SRR8145879 |
| 365 | Negative | CcspTutym | *Coloceras* | *theresae* | *Turtur tympanistria* | Democratic Republic of Congo | SRR8145878 |
| 366 | Negative | Cctav | *Coloceras* | *tavornikae* | *Columba livia* | Canada | SRR5308153 |
| 367 | Negative | CospPaoen | *Columbicola* | *adamsi* | *Patagioenas oenops* | Peru | SRR3161963 |
| 368 | Negative | Coada | *Columbicola* | *adamsi* | *Patagioenas speciosa* | Mexico | SRR3161912 |
| 369 | Negative | Cobac | *Columbicola* | *baculoides* | *Zenaida macroura* | United States of America | SRR13159343 |

| Count_number | Sodalis_detection | Library_code | Louse_Genus | Louse_Species | Vertebrate_Host | Collection_Country | NCBI_SRA |
|---|---|---|---|---|---|---|---|
| 370 | Negative | Cocol2 | *Columbicola* | *columbae 2* | *Columba guinea* | South Africa | SRR3161917 |
| 371 | Negative | Coelb209 | *Columbicola* | *elbeli* | *Treron vernans* | Malaysia | SRR16503027 |
| 372 | Negative | Coelb | *Columbicola* | *elbeli* | *Treron vernans* | Malaysia | SRR3161966 |
| 373 | Negative | CoexiMaruf | *Columbicola* | *exilicornis 3* | *Macropygia ruficeps* | Malaysia | SRR16503018 |
| 374 | Negative | Coexi3 | *Columbicola* | *exilicornis 3* | *Macropygia ruficeps* | Malaysia | SRR3161962 |
| 375 | Negative | Cofor | *Columbicola* | *fortis* | *Otidiphaps nobilis* | Papua New Guinea | SRR3161925 |
| 376 | Negative | CospAplar | *Columbicola* | *fradei* | *Aplopelia larvata* | Malawi | SRR3161954 |
| 377 | Negative | Cogra | *Columbicola* | *gracilicapitis* | *Leptotila jamaicensis* | Mexico | SRR3161913 |
| 378 | Negative | Cogui2 | *Columbicola* | *guimaraesi 2* | *Chalcophaps indica* | Australia | SRR16503017 |
| 379 | Negative | Cohoo | *Columbicola* | *hoogstraali* | *Nesoenas picturatus* | Madagascar | SRR3161968 |
| 380 | Negative | ComacZeart | *Columbicola* | *macrourae* | *Zenaida aurita* | Jamaica | SRR16503014 |
| 381 | Negative | ComacLever | *Columbicola* | *macrourae 1* | *Leptotila verreauxi* | Brazil | SRR16503015 |
| 382 | Negative | ComacZeasi | *Columbicola* | *macrourae 2* | *Zenaida asiatica* | United States of America | SRR16503059 |
| 383 | Negative | Comac2 | *Columbicola* | *macrourae 2* | *Zenaida asiatica* | United States of America | SRR3161952 |
| 384 | Negative | ComacZemac | *Columbicola* | *macrourae 3* | *Zenaida macroura* | United States of America | SRR16503058 |
| 385 | Negative | Comac3 | *Columbicola* | *macrourae 3* | *Zenaida macroura* | United States of America | SRR3161953 |
| 386 | Negative | ComjoGestr | *Columbicola* | *mjoebergi* | *Geopelia striata* | United States of America | SRR9693822 |
| 387 | Negative | ComjoGecun | *Columbicola* | *mjoebergi 1* | *Geopelia cuneata* | Australia | SRR16503056 |
| 388 | Negative | Comjo1 | *Columbicola* | *mjoebergi 1* | *Geophaps smithii* | Australia | SRR3161957 |
| 389 | Negative | Coplm33389 | *Columbicola* | *palmai* | *Leucosarcia melanoleuca* | Australia | SRR16503055 |
| 390 | Negative | Coplm | *Columbicola* | *palmai* | *Leucosarcia melanoleuca* | Australia | SRR3161932 |
| 391 | Negative | PnpicBlrub | *Columbicola* | *sp.* | *Blythipicus rubiginosus* | Borneo | SRR9693808 |
| 392 | Negative | CospGevrg | *Columbicola* | *sp.* | *Leptotrygon veraguensis* | Panama | SRR16503054 |
| 393 | Negative | CospOecap | *Columbicola* | *sp.* | *Oena capensis* | Madagascar | SRR8177104 |
| 394 | Negative | Tusp.Tutym | *Columbicola* | *sp.* | *Turtur tympanistria* | Malawi | SRR5308256 |
| 395 | Negative | Cotch266 | *Columbicola* | *taschenbergi* | *Reinwardtoena reinwardti* | Papua New Guinea | SRR16503052 |
| 396 | Negative | Cotch | *Columbicola* | *taschenbergi* | *Reinwardtoena reinwardti* | Papua New Guinea | SRR3161928 |
| 397 | Negative | Cotim2170 | *Columbicola* | *timmermanni* | *Leptotila rufaxilla* | Guyana | SRR16503051 |
| 398 | Negative | Cotim | *Columbicola* | *timmermanni* | *Leptotila rufaxilla* | Guyana | SRR3161965 |
| 399 | Negative | Cowag | *Columbicola* | *waggermanni* | *Patagioenas leucocephala* | Jamaica | SRR16503050 |
| 400 | Negative | Cowai | *Columbicola* | *waiteae* | *Columba leucomela* | Australia | SRR3161940 |
| 401 | Negative | Cowal | *Columbicola* | *waltheri* | *Zentrygon frenata* | Peru | SRR3161933 |
| 402 | Negative | Cowec272 | *Columbicola* | *wecksteini* | *Ptilinopus rivoli* | Papua New Guinea | SRR16503048 |
| 403 | Negative | Cowec | *Columbicola* | *wecksteini* | *Ptilinopus rivoli* | Papua New Guinea | SRR3161926 |
| 404 | Negative | Coeme4 | *Columbicola* | *emersoni 4* | *Ptilinopus regina* | Australia | SRR8145887 |
| 405 | Negative | Cowol23 | *Columbicola* | *wolffhuegeli* | *Ducula bicolor* | Australia | SRR8145890 |
| 406 | Negative | Cowol76 | *Columbicola* | *wolffhuegeli* | *Ducula bicolor* | Australia | SRR8145891 |
| 407 | Negative | Brqua | *Corvonirmus* | *quadrangularis* | *Corvus albus* | Malawi | SRR8334237 |

| Count_number | Sodalis_detection | Library_code | Louse_Genus | Louse_Species | Vertebrate_Host | Collection_Country | NCBI_SRA |
|---|---|---|---|---|---|---|---|
| 408 | Negative | Brrot | *Corvonirmus* | *rotundata* | *Corvus brachyrhynchos* | Canada | SRR14141056 |
| 409 | Negative | Coeng | *Cotingacola* | *engeli* | *Phoenicircus nigricollis* | Brazil | SRR5308209 |
| 410 | Negative | Coter | *Cotingacola* | *tergalis* | *Pipreola riefferii chachapoyas* | Peru | SRR5308213 |
| 411 | Negative | Cfpac | *Craspedorrhynchus* | *pachypus* | *Haliastur sphenurus* | Australia | SRR17556669 |
| 412 | Negative | CfspFaber | *Craspedorrhynchus* | *sp.* | *Falco berigora* | Australia | SRR8145880 |
| 413 | Negative | CfspBualb | *Craspedorrhynchus* | *sp.* | *Geranoaetus albicaudatus* | United States of America | SRR17556668 |
| 414 | Negative | Cfsub | *Craspedorrhynchus* | *subhaematopus* | *Accipiter cooperii* | Canada | SRR5308371 |
| 415 | Negative | CkspNodar | *Cuclotocephalus* | *sp.* | *Nothura darwinii* | Bolivia | SRR13159337 |
| 416 | Negative | Cgcin | *Cuclotogaster* | *cinereus* | *Coturnix coturnix africana* | Malawi | SRR5308207 |
| 417 | Negative | Liesp | *Cuclotogaster* | *sp.* | *Pternistis capensis* | South Africa | SRR5308232 |
| 418 | Negative | Cgsp.Frhil | *Cuclotogaster* | *sp.* | *Pternistis hildebrandti* | Malawi | SRR5308208 |
| 419 | Negative | Cuspl | *Cuculicola* | *splendidus* | *Geococcyx californianus* | United States of America | SRR5308218 |
| 420 | Negative | Csafr | *Cuculoecus* | *africanus* | *Chrysococcyx cupreus* | Ghana | SRR5308372 |
| 421 | Negative | Cspia | *Cuculoecus* | *piayae* | *Piaya cayana* | Panama | SRR17556667 |
| 422 | Negative | Daasy1 | *Dahlemhornia* | *asymmetrica* | *Dromaius novaehollandiae* | Australia | SRR5308358 |
| 423 | Negative | Daasy2 | *Dahlemhornia* | *asymmetrica* | *Dromaius novaehollandiae* | Australia | SRR16574600 |
| 424 | Negative | Daasy5 | *Dahlemhornia* | *asymmetrica* | *Dromaius novaehollandiae* | Australia | SRR5308359 |
| 425 | Negative | DgspBujam | *Degeeriella* | *sp.* | *Buteo jamaicensis harlani* | Canada | SRR5308224 |
| 426 | Negative | Dimex | *Discocorpus* | *mexicanus* | *Crypturellus cinnamomeus* | Mexico | SRR5308387 |
| 427 | Negative | DispCrobs | *Discocorpus* | *sp.* | *Crypturellus obsoletus* | Brazil | SRR8172518 |
| 428 | Negative | DispCrstr | *Discocorpus* | *sp.* | *Crypturellus strigulosus* | Brazil | SRR12762519 |
| 429 | Negative | DispCrtra | *Discocorpus* | *sp.* | *Crypturellus transfasciatus* | Peru | SRR14908129 |
| 430 | Negative | Dosex | *Docophorocotes* | *sexsetosus* | *Rhynchotus rufescens* | Argentina | SRR16503047 |
| 431 | Negative | Dobre | *Docophoroides* | *brevis* | *Diomedea exulans* | United Kingdom | SRR5308117 |
| 432 | Negative | Etang | *Echinophilopterus* | *angustoclypeatus* | *Polytelis anthopeplus* | Australia | SRR8177112 |
| 433 | Negative | EtspPocry | *Echinophilopterus* | *sp.* | *Poicephalus cryptoxanthus* | Malawi | SRR8334244 |
| 434 | Negative | Embra | *Emersoniella* | *bracteata* | *Dacelo novaeguineae* | Australia | SRR5308227 |
| 435 | Negative | EmspTasyl | *Emersoniella* | *sp.* | *Tanysiptera sylvia* | Australia | SRR5308228 |
| 436 | Negative | EpspCarub | *Epipicus* | *sp.* | *Campephilus rubricollis* | Brazil | SRR5308155 |
| 437 | Negative | Epped | *Episbates* | *pederiformis* | *Phoebastria irrorata* | Ecuador | SRR5308154 |
| 438 | Negative | Esbre | *Esthiopterum* | *brevicephalum* | *Antigone canadensis* | Canada | SRR5308385 |
| 439 | Negative | Famar | *Falcolipeurus* | *marginalis* | *Cathartes aura* | United States of America | SRR5308118 |
| 440 | Negative | Fasec | *Falcolipeurus* | *secretarius* | *Sagittarius serpentarius* | Kenya | SRR16574598 |
| 441 | Negative | FaspAqwah | *Falcolipeurus* | *sp.* | *Hieraaetus wahlbergi* | Malawi | SRR8334245 |
| 442 | Negative | FfforPlads | *Forficuloecus* | *forficula* | *Platycercus adscitus* | Australia | SRR8177113 |
| 443 | Negative | FfforPlele | *Forficuloecus* | *forficula* | *Platycercus elegans* | Australia | SRR8145951 |
| 444 | Negative | Ffjos | *Forficuloecus* | *josephi* | *Neophema bourkii* | Australia | SRR8175156 |
| 445 | Negative | Ffpal | *Forficuloecus* | *palmai* | *Barnardius zonarius* | Australia | SRR5308157 |

| Count_number | Sodalis_detection | Library_code | Louse_Genus | Louse_Species | Vertebrate_Host | Collection_Country | NCBI_SRA |
|---|---|---|---|---|---|---|---|
| 446 | Negative | FfspCynov | *Forficuloecus* | *sp.* | *Cyanoramphus novaezelandiae* | Australia | SRR8582567 |
| 447 | Negative | FfspNemer | *Forficuloecus* | *sp.* | *Nestor meridionalis septentrionalis* | Australia | SRR8582568 |
| 448 | Negative | FfpsPsdis | *Forficuloecus* | *sp.* | *Psephotus dissimilis* | Australia | SRR8177114 |
| 449 | Negative | FfspPshae | *Forficuloecus* | *sp.* | *Psephotus haematonotus* | Australia | SRR8145952 |
| 450 | Negative | Ffwil | *Forficuloecus* | *wilsoni* | *Platycercus venustus* | Australia | SRR8177115 |
| 451 | Negative | FmspHypoe67 | *Formicaphagus* | *sp.* | *Willisornis poecilinotus* | Brazil | SRR8582570 |
| 452 | Negative | FoanaFocol430 | *Formicaricola* | *analoides* | *Formicarius colma* | Brazil | SRR8582571 |
| 453 | Negative | Fulon | *Fulicoffula* | *longipila* | *Fulica americana* | United States of America | SRR5308119 |
| 454 | Negative | FulurFuatr | *Fulicoffula* | *lurida* | *Fulica atra* | Australia | SRR16574597 |
| 455 | Negative | FulurFucri | *Fulicoffula* | *lurida* | *Fulica cristata* | South Africa | SRR16574596 |
| 456 | Negative | FulurFuleu | *Fulicoffula* | *lurida* | *Fulica leucoptera* | Argentina | SRR13389481 |
| 457 | Negative | FuspFuala | *Fulicoffula* | *sp.* | *Fulica alai* | United States of America | SRR17556666 |
| 458 | Negative | FuspFuard | *Fulicoffula* | *sp.* | *Fulica ardesiaca* | Argentina | SRR13389480 |
| 459 | Negative | FuspFuarm | *Fulicoffula* | *sp.* | *Fulica armillata* | Argentina | SRR13389479 |
| 460 | Negative | FuspFugig | *Fulicoffula* | *sp.* | *Fulica gigantea* | Argentina | SRR13389478 |
| 461 | Negative | FuspFuruf | *Fulicoffula* | *sp.* | *Fulica rufifrons* | Argentina | SRR17556665 |
| 462 | Negative | FuspPocar | *Fulicoffula* | *sp.* | *Porzana carolina* | United States of America | SRR8334257 |
| 463 | Negative | FnspAupar | *Furnariphilus* | *sp.* | *Automolus paraensis* | Brazil | SRR8334255 |
| 464 | Negative | FispScgua | *Furnariphilus* | *sp.* | *Sclerurus guatemalensis* | Panama | SRR8145953 |
| 465 | Negative | GospAllat | *Goniocotes* | *sp.* | *Alectura lathami* | Australia | SRR13389466 |
| 466 | Negative | GospMerei | *Goniocotes* | *sp.* | *Megapodius reinwardt* | Australia | SRR8145957 |
| 467 | Negative | GospPtpet | *Goniocotes* | *sp.* | *Ptilopachus petrosus* | Ghana | SRR13389465 |
| 468 | Negative | GospTycup | *Goniocotes* | *sp.* | *Tympanuchus cupido attwateri* | United States of America | SRR13389463 |
| 469 | Negative | Gotal | *Goniocotes* | *talegallae* | *Talegalla fuscirostris* | Papua New Guinea | SRR13389462 |
| 470 | Negative | Gdast | *Goniodes* | *astrocephalus* | *Coturnix coturnix* | Russia | SRR13389477 |
| 471 | Negative | Gdbio | *Goniodes* | *biordinatus* | *Megapodius reinwardt* | Australia | SRR8145955 |
| 472 | Negative | Gdcen | *Goniodes* | *centrocerci* | *Centrocercus urophasianus* | United States of America | SRR8582573 |
| 473 | Negative | Gdcor | *Goniodes* | *coronatus* | *Rollulus rouloul* | Malaysia | SRR13389476 |
| 474 | Negative | Gdmer | *Goniodes* | *merriamanus* | *Dendragapus obscurus* | United States of America | SRR8582574 |
| 475 | Negative | Gdort | *Goniodes* | *ortygis* | *Colinus virginianus* | United States of America | SRR5308120 |
| 476 | Negative | GdspAllat | *Goniodes* | *sp.* | *Alectura lathami* | Australia | SRR13389474 |
| 477 | Negative | GdspCacal | *Goniodes* | *sp.* | *Callipepla californica* | United States of America | SRR13389473 |
| 478 | Negative | GdspCrmon | *Goniodes* | *sp.* | *Cyrtonyx montezumae* | United States of America | SRR13389472 |
| 479 | Negative | GdspNumel | *Goniodes* | *sp.* | *Numida meleagris* | South Africa | SRR13389469 |
| 480 | Negative | GdspPhcol | *Goniodes* | *sp.* | *Phasianus colchicus* | United States of America | SRR13389468 |
| 481 | Negative | GdspFraha | *Goniodes* | *sp.* | *Pternistis ahantensis* | Ghana | SRR13389471 |
| 482 | Negative | GdspFrbic | *Goniodes* | *sp.* | *Pternistis bicalcaratus* | Ghana | SRR13389470 |
| 483 | Negative | Gdass | *Goniodes* | *sp.* | *Ptilopachus petrosus* | Ghana | SRR8175159 |

| Count_number | Sodalis_detection | Library_code | Louse_Genus | Louse_Species | Vertebrate_Host | Collection_Country | NCBI_SRA |
|---|---|---|---|---|---|---|---|
| 484 | Negative | GdspPtswa | *Gonoides* | *sp.* | *Pternistis swainsonii* | South Africa | SRR13389467 |
| 485 | Negative | GuspSeper | *Guimaraesiella* | *sp.* | *Aethomyias perspicillatus* | New Guinea | SRR17556645 |
| 486 | Negative | BrspAicra | *Guimaraesiella* | *sp.* | *Ailuroedus crassirostris* | Australia | SRR16574607 |
| 487 | Negative | BrspAlmor | *Guimaraesiella* | *sp.* | *Alcippe morrisonia* | China | SRR14141079 |
| 488 | Negative | GuspAraur | *Guimaraesiella* | *sp.* | *Arremon aurantiirostris* | Panama | SRR17556664 |
| 489 | Negative | BrspArcya | *Guimaraesiella* | *sp.* | *Artamus cyanopterus* | Australia | SRR14141053 |
| 490 | Negative | GuspBlcan | *Guimaraesiella* | *sp.* | *Bleda canicapillus* | Ghana | SRR17556663 |
| 491 | Negative | GuspBlexi | *Guimaraesiella* | *sp.* | *Bleda eximius* | Ghana | SRR17556662 |
| 492 | Negative | GuspBlsyn | *Guimaraesiella* | *sp.* | *Bleda syndactylus* | Ghana | SRR18820758 |
| 493 | Negative | GuspChpap | *Guimaraesiella* | *sp.* | *Chaetorhynchus papuensis* | Papua New Guinea | SRR18820755 |
| 494 | Negative | GuspChnuc | *Guimaraesiella* | *sp.* | *Chlamydera nuchalis* | Australia | SRR18820756 |
| 495 | Negative | BrspClpic | *Guimaraesiella* | *sp.* | *Climacteris picumnus* | Australia | SRR16574605 |
| 496 | Negative | GuspCohar | *Guimaraesiella* | *sp.* | *Colluricincla harmonica* | Australia | SRR17556660 |
| 497 | Negative | GuspCrbar | *Guimaraesiella* | *sp.* | *Criniger barbatus* | Ghana | SRR17556658 |
| 498 | Negative | GuspFafro | *Guimaraesiella* | *sp.* | *Falcunculus frontatus* | Australia | SRR17556656 |
| 499 | Negative | BrspHyleu | *Guimaraesiella* | *sp.* | *Hypsipetes leucocephalus* | China | SRR14141074 |
| 500 | Negative | GuspManit | *Guimaraesiella* | *sp.* | *Malimbus nitens* | Ghana | SRR17556655 |
| 501 | Negative | GuspMycol | *Guimaraesiella* | *sp.* | *Myadestes coloratus* | Panama | SRR17556654 |
| 502 | Negative | BrspMycol | *Guimaraesiella* | *sp.* | *Myadestes coloratus* | Panama | SRR14141050 |
| 503 | Negative | GuspNepoe | *Guimaraesiella* | *sp.* | *Neocossyphus poensis* | Ghana | SRR17556653 |
| 504 | Negative | BrspPaele | *Guimaraesiella* | *sp.* | *Periparus elegans* | Philippines | SRR18820725 |
| 505 | Negative | GuspPhalb | *Guimaraesiella* | *sp.* | *Phyllastrephus albigularis* | Ghana | SRR17556650 |
| 506 | Negative | GuspPhict | *Guimaraesiella* | *sp.* | *Phyllastrephus icterinus* | Ghana | SRR17556649 |
| 507 | Negative | GuspPyplu | *Guimaraesiella* | *sp.* | *Pycnonotus plumosus* | Malaysia | SRR17556647 |
| 508 | Negative | GuspSenoh | *Guimaraesiella* | *sp.* | *Sericornis nouhuysi* | New Guinea | SRR17556646 |
| 509 | Negative | BrspTrvir | *Guimaraesiella* | *sp.* | *Trogon viridis* | Brazil | SRR8334239 |
| 510 | Negative | GuspZohei | *Guimaraesiella* | *sp.* | *Zoothera heinei* | Australia | SRR17556642 |
| 511 | Negative | GuspZolun | *Guimaraesiella* | *sp.* | *Zoothera lunulata* | Australia | SRR17556641 |
| 512 | Negative | Hfgra | *Haffneria* | *grandis* | *Stercorarius skua* | United Kingdom | SRR5308162 |
| 513 | Negative | Hlabn | *Halipeurus* | *abnormis* | *Calonectris diomedea* | United States of America | SRR8334260 |
| 514 | Negative | Hldiv | *Halipeurus* | *diversus* | *Ardenna tenuirostris* | Australia | SRR5308124 |
| 515 | Negative | Hrden | *Harrisoniella* | *densa* | *Phoebastria immutabilis* | United States of America | SRR5308163 |
| 516 | Negative | Hebol | *Heptapsogaster* | *bolivianus* | *Nothura darwinii* | Peru | SRR16503046 |
| 517 | Negative | Hefri | *Heptapsogaster* | *frielingi* | *Cariama cristata* | Brazil | SRR8175160 |
| 518 | Negative | HespCrsou2 | *Heptapsogaster* | *inexpectatus* | *Crypturellus soui* | Brazil | SRR14908119 |
| 519 | Negative | HespCrsou | *Heptapsogaster* | *mandibularis* | *Crypturellus soui* | Brazil | SRR14908101 |
| 520 | Negative | Heman | *Heptapsogaster* | *mandibularis* | *Crypturellus tataupa* | Brazil | SRR12762508 |
| 521 | Negative | Hemin | *Heptapsogaster* | *minor* | *Nothura maculosa* | Argentina | SRR8175161 |

| Count_number | Sodalis_detection | Library_code | Louse_Genus | Louse_Species | Vertebrate_Host | Collection_Country | NCBI_SRA |
|---|---|---|---|---|---|---|---|
| 522 | Negative | Henot | *Heptapsogaster* | *nothocercae* | *Nothocercus nigrocapillus* | Peru | SRR14908106 |
| 523 | Negative | HespChbur | *Heptapsogaster* | *sp.* | *Chunga burmeisteri* | Bolivia | SRR8145958 |
| 524 | Negative | HespCrbou | *Heptapsogaster* | *sp.* | *Crypturellus boucardi* | Mexico | SRR14908105 |
| 525 | Negative | HespCrobs | *Heptapsogaster* | *sp.* | *Crypturellus obsoletus* | Brazil | SRR13159334 |
| 526 | Negative | HespCrobsPE | *Heptapsogaster* | *sp.* | *Crypturellus obsoletus* | Peru | SRR14908103 |
| 527 | Negative | HespCrpar | *Heptapsogaster* | *sp.* | *Crypturellus parvirostris* | Bolivia | SRR13159309 |
| 528 | Negative | HespCrstr | *Heptapsogaster* | *sp.* | *Crypturellus strigulosus* | Brazil | SRR5308161 |
| 529 | Negative | HespCrtra | *Heptapsogaster* | *sp.* | *Crypturellus transfasciatus* | Peru | SRR14908102 |
| 530 | Negative | HespCrund | *Heptapsogaster* | *sp.* | *Crypturellus undulatus* | Bolivia | SRR12762497 |
| 531 | Negative | HespEuele | *Heptapsogaster* | *sp.* | *Eudromia elegans* | Argentina | SRR8175162 |
| 532 | Negative | HespNocin | *Heptapsogaster* | *sp.* | *Nothoprocta cinerascens* | Argentina | SRR14908148 |
| 533 | Negative | HespNopen | *Heptapsogaster* | *sp.* | *Nothoprocta pentlandii* | Argentina | SRR14908147 |
| 534 | Negative | HespNobor | *Heptapsogaster* | *sp.* | *Nothura boraquira* | Bolivia | SRR13159298 |
| 535 | Negative | HespNodar2 | *Heptapsogaster* | *sp.* | *Nothura darwinii* | Peru | SRR16503045 |
| 536 | Negative | HespNodar | *Heptapsogaster* | *sp.* | *Nothura darwinii* | Bolivia | SRR13159338 |
| 537 | Negative | HespTigut | *Heptapsogaster* | *sp.* | *Tinamus guttatus* | Brazil | SRR8175163 |
| 538 | Negative | HespCrstr2 | *Heptapsogaster* | *sp. 2* | *Crypturellus strigulosus* | Brazil | SRR13159336 |
| 539 | Negative | HespCrbou2 | *Heptapsogaster* | *sp.2* | *Crypturellus boucardi* | Mexico | SRR14908104 |
| 540 | Negative | Hesub | *Heptapsogaster* | *subteres* | *Nothoprocta pentlandii* | Argentina | SRR16503040 |
| 541 | Negative | Heteres | *Heptapsogaster* | *teres* | *Nothoprocta perdicaria* | Chile | SRR16503039 |
| 542 | Negative | Hetes | *Heptapsogaster* | *tesselatus* | *Nothoprocta curvirostris* | Peru | SRR14908145 |
| 543 | Negative | Hetest | *Heptapsogaster* | *testudo* | *Nothoprocta cinerascens* | Argentina | SRR14908144 |
| 544 | Negative | HespNoper | *Heptapsogaster* | *testudo* | *Nothoprocta perdicaria* | Chile | SRR16503044 |
| 545 | Negative | Hetru | *Heptapsogaster* | *truncatus* | *Nothoprocta ornata* | Bolivia | SRR13159340 |
| 546 | Negative | Heine | *Heptapsus* | *inexpectatus* | *Nothocercus nigrocapillus* | Peru | SRR14908115 |
| 547 | Negative | Heter | *Heptapsus* | *tergalis* | *Nothocercus nigrocapillus* | Peru | SRR14908146 |
| 548 | Negative | HespTimaj | *Heptarthrogaster* | *sp.1* | *Tinamus major* | Brazil | SRR16503042 |
| 549 | Negative | HespTimaj2 | *Heptarthrogaster* | *sp.2* | *Tinamus major* | Brazil | SRR16503041 |
| 550 | Negative | HespTigut2 | *Heptathrogaster* | *sp.* | *Tinamus guttatus* | Brazil | SRR16503043 |
| 551 | Negative | Hkhop | *Hopkinsiella* | *hopkinsi* | *Phoeniculus bollei jacksoni* | Democratic Republic of Congo | SRR5308229 |
| 552 | Negative | HyspCrobs | *Hypocrypturellus* | *sp.* | *Crypturellus obsoletus* | Peru | SRR14908143 |
| 553 | Negative | HyspCrsou | *Hypocrypturellus* | *sp.* | *Crypturellus soui* | Brazil | SRR14908142 |
| 554 | Negative | HyspTimaj | *Hypocrypturellus* | *sp.* | *Tinamus major* | Brazil | SRR16503036 |
| 555 | Negative | HyspTigut | *Hyprocrypturellus* | *sp.* | *Tinamus guttatus* | Brazil | SRR16503037 |
| 556 | Negative | HyspTitao | *Hyprocrypturellus* | *sp.* | *Tinamus tao* | Peru | SRR16503035 |
| 557 | Negative | Ibaus | *Ibidoecus* | *australis* | *Threskiornis spinicollis* | Australia | SRR17556640 |
| 558 | Negative | Ibbis | *Ibidoecus* | *bisignatus* | *Plegadis chihi* | United States of America | SRR5308126 |
| 559 | Negative | IbbisPlfal | *Ibidoecus* | *bisignatus* | *Plegadis falcinellus* | Australia | SRR17556639 |

| Count_number | Sodalis_detection | Library_code | Louse_Genus | Louse_Species | Vertebrate_Host | Collection_Country | NCBI_SRA |
|---|---|---|---|---|---|---|---|
| 560 | Negative | Ibdia | *Ibidoecus* | *dianae* | *Threskiornis molucca* | Australia | SRR13389461 |
| 561 | Negative | InspRaele | *Incidifrons* | *sp.* | *Rallus elegans* | United States of America | SRR8566360 |
| 562 | Negative | Intra | *Incidifrons* | *transpositus* | *Fulica americana* | Canada | SRR5308164 |
| 563 | Negative | BrspErtri | *Indoceoplanetes* | *sp.* | *Erythrura trichroa* | Australia | SRR14141054 |
| 564 | Negative | FuspSaruf | Ischnocera | *sp.* | *Sarothrura rufa* | Malawi | SRR8334258 |
| 565 | Negative | Kemen | *Kelloggia* | *mendax* | *Crypturellus parvirostris* | Bolivia | SRR13159331 |
| 566 | Negative | KespCrtat | *Kelloggia* | *sp.* | *Crypturellus tataupa* | Brazil | SRR12762534 |
| 567 | Negative | KespTigut | *Kelloggia* | *sp.* | *Tinamus guttatus* | Brazil | SRR8145965 |
| 568 | Negative | Keund | *Kelloggia* | *undulatus* | *Crypturellus undulatus* | Bolivia | SRR12762525 |
| 569 | Negative | Kobra | *Kodocephalon* | *bradicephalum* | *Goura scheepmakeri* | Papua New Guinea | SRR18820754 |
| 570 | Negative | Kosub | *Kodocephalon* | *suborbiculatum* | *Goura victoria* | Papua New Guinea | SRR5308166 |
| 571 | Negative | LbspPesup | *Labicotes* | *sp.* | *Penelope superciliaris* | Brazil | SRR5308167 |
| 572 | Negative | Lggib | *Lagopoecus* | *gibsoni* | *Centrocercus urophasianus* | United States of America | SRR5308168 |
| 573 | Negative | Lgobs | *Lagopoecus* | *obscurus* | *Dendragapus obscurus* | United States of America | SRR8582575 |
| 574 | Negative | Lphir | *Lamprocorpus* | *hirsutus* | *Nothoprocta ornata* | Bolivia | SRR8145973 |
| 575 | Negative | LpspNodar | *Lamprocorpus* | *sp.* | *Nothura darwinii* | Bolivia | SRR13159339 |
| 576 | Negative | Lpspi | *Lamprocorpus* | *spinosus* | *Nothoprocta curvirostris* | Peru | SRR14908141 |
| 577 | Negative | Licap | *Lipeurus* | *caponis* | *Gallus gallus* | Ecuador | SRR5308373 |
| 578 | Negative | LicraAllat | *Lipeurus* | *crassus* | *Alectura lathami* | Australia | SRR5308231 |
| 579 | Negative | Oxina | *Lipeurus* | *inaequalis* | *Megapodius reinwardt* | Australia | SRR16574591 |
| 580 | Negative | Lisp.Numel | *Lipeurus* | *sp.* | *Numida meleagris* | Malawi | SRR5308233 |
| 581 | Negative | LispPtswa | *Lipeurus* | *sp.* | *Pternistis swainsonii* | South Africa | SRR13389508 |
| 582 | Negative | Lndro | *Lunaceps* | *drosti* | *Calidris tenuirostris* | Japan | SRR18820753 |
| 583 | Negative | Lnnum | *Lunaceps* | *numenii* | *Numenius phaeopus* | Australia | SRR8175164 |
| 584 | Negative | LnspTrsub | *Lunaceps* | *sp.* | *Calidris subruficollis* | Brazil | SRR5809352 |
| 585 | Negative | BrspOrfla | *Maculinirmus* | *sp.* | *Oriolus flavocinctus* | Australia | SRR14141060 |
| 586 | Negative | Mgema | *Megaginus* | *emarginatus* | *Crypturellus obsoletus* | Peru | SRR14908136 |
| 587 | Negative | MgspCrsou | *Megaginus* | *sp.* | *Crypturellus soui* | Brazil | SRR14908135 |
| 588 | Negative | MgspCrund | *Megaginus* | *sp.* | *Crypturellus undulatus* | Brazil | SRR12762524 |
| 589 | Negative | Mgtat | *Megaginus* | *tataupensis* | *Crypturellus tataupa* | Bolivia | SRR5308131 |
| 590 | Negative | Mpasy | *Megapeostus* | *asymmetricus* | *Crypturellus undulatus* | Bolivia | SRR12762523 |
| 591 | Negative | Mphep | *Megapeostus* | *heptarthrogastriformis* | *Crypturellus obsoletus* | Brazil | SRR8175166 |
| 592 | Negative | Mppla | *Megapeostus* | *petersi* | *Crypturellus soui* | Brazil | SRR14908134 |
| 593 | Negative | Mppet | *Megapeostus* | *petersi* | *Crypturellus tataupa* | Brazil | SRR12762522 |
| 594 | Negative | MespCrobs | *Megapeostus* | *sp.* | *Crypturellus obsoletus* | Peru | SRR14908137 |
| 595 | Negative | MpspCrstr | *Megapeostus* | *sp.* | *Crypturellus strigulosus* | Brazil | SRR5308169 |
| 596 | Negative | MpspCrcin1 | *Megapeostus* | *sp. 1* | *Crypturellus cinnamomeus* | Mexico | SRR13159296 |
| 597 | Negative | MrspMeorn | *Meropoecus* | *sp.* | *Merops ornatus* | Australia | SRR5308363 |

| Count_number | Sodalis_detection | Library_code | Louse_Genus | Louse_Species | Vertebrate_Host | Collection_Country | NCBI_SRA |
|---|---|---|---|---|---|---|---|
| 598 | Negative | BrspMegul | *Meropsiella* | *sp.* | *Merops gularis* | Ghana | SRR14141051 |
| 599 | Negative | BrspMeorn | *Meropsiella* | *sp.* | *Merops ornatus* | Australia | SRR5308362 |
| 600 | Negative | BrspLofus | *Mirandofures* | *sp.* | *Lonchura fuscans* | Malaysia | SRR8142483 |
| 601 | Negative | BrspLostr | *Mirandofures* | *sp.* | *Lonchura striata* | China | SRR8582562 |
| 602 | Negative | Brmar | *Motmotnirmus* | *marginellus* | *Momotus momota* | Peru | SRR8142478 |
| 603 | Negative | MuspEuarg | *Mulcticola* | *sp.* | *Eurostopodus argus* | Australia | SRR8145980 |
| 604 | Negative | MuspNyalb | *Mulcticola* | *sp.* | *Nyctidromus albicollis* | Panama | SRR5308374 |
| 605 | Negative | Nahar | *Naubates* | *harrisoni* | *Puffinus boydi* | Cape Verde | SRR5308170 |
| 606 | Negative | Npinc | *Neophilopterus* | *incompletus* | *Ciconia ciconia* | Sweden | SRR9693827 |
| 607 | Negative | Nsafr | *Neopsittaconirmus* | *africanus* | *Poicephalus gulielmi* | Ghana | SRR8177119 |
| 608 | Negative | Nsalb | *Neopsittaconirmus* | *albus* | *Cacatua galerita* | Australia | SRR8145986 |
| 609 | Negative | NseosCasan | *Neopsittaconirmus* | *eos* | *Cacatua sanguinea* | Australia | SRR8177122 |
| 610 | Negative | NseosEoros | *Neopsittaconirmus* | *eos* | *Eolophus roseicapilla* | Australia | SRR8175171 |
| 611 | Negative | NsspCalea | *Neopsittaconirmus* | *sp.* | *Cacatua leadbeateri* | Australia | SRR8145988 |
| 612 | Negative | NsspNyhol | *Neopsittaconirmus* | *sp.* | *Nymphicus hollandicus* | Australia | SRR8145989 |
| 613 | Negative | NsspPosen | *Neopsittaconirmus* | *sp.* | *Poicephalus senegalus* | Ghana | SRR8175173 |
| 614 | Negative | NsspPshae | *Neopsittaconirmus* | *sp.* | *Psephotus haematonotus* | Australia | SRR8175174 |
| 615 | Negative | NsspPsvar | *Neopsittaconirmus* | *sp.* | *Psephotus varius* | Australia | SRR8145990 |
| 616 | Negative | Nndem1 | *Nesiotinus* | *demersus* | *Aptenodytes patagonicus* | Crozet Island | SRR9693830 |
| 617 | Negative | Nndem2 | *Nesiotinus* | *demersus* | *Aptenodytes patagonicus* | Crozet Island | SRR9693831 |
| 618 | Negative | Nndem3 | *Nesiotinus* | *demersus* | *Aptenodytes patagonicus* | Crozet Island | SRR9693828 |
| 619 | Negative | Nndem4 | *Nesiotinus* | *demersus* | *Aptenodytes patagonicus* | Crozet Island | SRR9693829 |
| 620 | Negative | NospNonig | *Nothocolus* | *sp.* | *Nothocercus nigrocapillus* | Peru | SRR14908133 |
| 621 | Negative | Nosub | *Nothocotus* | *subsimilis* | *Nothocercus nigrocapillus* | Peru | SRR8334267 |
| 622 | Negative | NylonNygri | *Nyctibicola* | *longirostris* | *Nyctibius griseus* | Brazil | SRR8177125 |
| 623 | Negative | Nylon | *Nyctibicola* | *longirostris* | *Nyctibius jamaicensis* | Mexico | SRR5308388 |
| 624 | Negative | Brmor | *Olivinirmus* | *moriona* | *Psilorhinus morio* | Mexico | SRR18820746 |
| 625 | Negative | OnspTigut | *Ornicholax* | *sp.* | *Tinamus guttatus* | Brazil | SRR16503034 |
| 626 | Negative | OnspTimaj | *Ornicholax* | *sp.* | *Tinamus major* | Brazil | SRR8566313 |
| 627 | Negative | OrspTitao | *Ornicholax* | *sp.* | *Tinamus tao* | Peru | SRR16503030 |
| 628 | Negative | OrspTimaj1 | *Ornicholax* | *sp.1* | *Tinamus major* | Brazil | SRR16503032 |
| 629 | Negative | OnspTigut2 | *Ornicholax* | *sp.2* | *Tinamus guttatus* | Brazil | SRR16503033 |
| 630 | Negative | Orgon | *Ornithobius* | *goniopleurus* | *Branta canadensis* | United States of America | SRR5308171 |
| 631 | Negative | Oscur1 | *Osculotes* | *curta* | *Opisthocomus hoazin* | Brazil | SRR16574594 |
| 632 | Negative | Oscur | *Osculotes* | *curta* | *Opisthocomus hoazin* | Peru | SRR5308133 |
| 633 | Negative | Osmac | *Osculotes* | *macropoda* | *Opisthocomus hoazin* | Brazil | SRR16574593 |
| 634 | Negative | Othou | *Otidoecus* | *houbarae* | *Chlamydotis undulata* | Morocco | SRR5308235 |
| 635 | Negative | Oxchi | *Oxylipeurus* | *chiniri* | *Ortalis vetula* | United States of America | SRR5308134 |

| Count_number | Sodalis_detection | Library_code | Louse_Genus | Louse_Species | Vertebrate_Host | Collection_Country | NCBI_SRA |
|---|---|---|---|---|---|---|---|
| 636 | Negative | Oxchi1 | *Oxylipeurus* | *chiniri* | *Ortalis vetula* | United States of America | SRR16574592 |
| 637 | Negative | Cxnum | *Oxylipeurus* | *numidianus* | *Colinus virginianus* | United States of America | SRR5308393 |
| 638 | Negative | Lisp.Rorou | *Oxylipeurus* | *sp.* | *Rollulus rouloul* | Malaysia | SRR5308234 |
| 639 | Negative | PaspPsvir | *Palmaellus* | *sp.* | *Psophia viridis* | Brazil | SRR5308173 |
| 640 | Negative | Pagra | *Paragoniocotes* | *grandis* | *Amazona amazonica* | Brazil | SRR5809339 |
| 641 | Negative | Pagua | *Paragoniocotes* | *guajirensis* | *Brotogeris jugularis* | Panama | SRR8175177 |
| 642 | Negative | Palon | *Paragoniocotes* | *longulufrons* | *Pionus sordidus* | Peru | SRR8334268 |
| 643 | Negative | PaspAmkaw | *Paragoniocotes* | *sp.* | *Amazona kawalli* | Brazil | SRR8334269 |
| 644 | Negative | PaspArsev | *Paragoniocotes* | *sp.* | *Ara severus* | Brazil | SRR8173262 |
| 645 | Negative | PaspArjan | *Paragoniocotes* | *sp.* | *Aratinga jandaya* | Brazil | SRR8334270 |
| 646 | Negative | PaspArast | *Paragoniocotes* | *sp.* | *Eupsittula nana* | Mexico | SRR8177126 |
| 647 | Negative | PaspPisen | *Paragoniocotes* | *sp.* | *Pionus senilis* | Panama | SRR5308172 |
| 648 | Negative | PaspArleu | *Paragoniocotes* | *sp.* | *Psittacara leucophthalmus* | Peru | SRR8334271 |
| 649 | Negative | PaspPylep248 | *Paragoniocotes* | *sp.* | *Pyrrhura lepida* | Brazil | SRR8582578 |
| 650 | Negative | PaspPyane | *Paragoniocotes* | *sp.* | *Pyrrhura lepida anerythra* | Brazil | SRR8173263 |
| 651 | Negative | PaspPymel | *Paragoniocotes* | *sp.* | *Pyrrhura melanura* | Brazil | SRR8334272 |
| 652 | Negative | PaspPyper | *Paragoniocotes* | *sp.* | *Pyrrhura perlata* | Brazil | SRR8173264 |
| 653 | Negative | Paven | *Paragoniocotes* | *venezolanus* | *Eupsittula pertinax* | Panama | SRR8173265 |
| 654 | Negative | PyspOdguj | *Passonomedea* | *sp.* | *Odontophorus gujanensis* | Brazil | SRR13389500 |
| 655 | Negative | PyspOdspe | *Passonomedea* | *sp.* | *Odontophorus speciosus* | Peru | SRR13389499 |
| 656 | Negative | PyspOdste | *Passonomedea* | *sp.* | *Odontophorus stellatus* | Brazil | SRR5308182 |
| 657 | Negative | Peang | *Pectenosoma* | *angusta* | *Crypturellus boucardi* | Mexico | SRR14908132 |
| 658 | Negative | Pemes | *Pectenosoma* | *meserythra* | *Crypturellus soui* | Brazil | SRR14908131 |
| 659 | Negative | Pepar | *Pectenosoma* | *parva* | *Crypturellus tataupa* | Brazil | SRR12762521 |
| 660 | Negative | PespCrstr | *Pectenosoma* | *sp.* | *Crypturellus strigulosus* | Brazil | SRR5308175 |
| 661 | Negative | PespCrtra | *Pectenosoma* | *sp.* | *Crypturellus transfasciatus* | Peru | SRR14908128 |
| 662 | Negative | Pesbp | *Pectenosoma* | *subparva* | *Crypturellus parvirostris* | Bolivia | SRR13159320 |
| 663 | Negative | Peyap | *Pectenosoma* | *yapurae* | *Crypturellus undulatus* | Bolivia | SRR12762518 |
| 664 | Negative | Pgacu | *Pectinopygus* | *acutofasciatus* | *Anhinga melanogaster* | Australia | SRR16574589 |
| 665 | Negative | Pgafe | *Pectinopygus* | *afer* | *Microcarbo africanus* | South Africa | SRR16574588 |
| 666 | Negative | Pgann | *Pectinopygus* | *annulatus* | *Sula dactylatra* | unknown | SRR16574587 |
| 667 | Negative | Pgbas | *Pectinopygus* | *bassani* | *Morus bassanus* | United States of America | SRR8177130 |
| 668 | Negative | PgbasMoser | *Pectinopygus* | *bassani* | *Morus serrator* | Australia | SRR13389507 |
| 669 | Negative | Pgdis | *Pectinopygus* | *dispar* | *Microcarbo melanoleucos* | Australia | SRR13389506 |
| 670 | Negative | Pgexc | *Pectinopygus* | *excornis* | *Microcarbo pygmaeus* | Italy | SRR13389505 |
| 671 | Negative | Pgfre | *Pectinopygus* | *fregatiphagus* | *Fregata magnificens* | Mexico | SRR16574586 |
| 672 | Negative | Pggra | *Pectinopygus* | *gracilicornis* | *Fregata minor* | unknown | SRR16574585 |
| 673 | Negative | Pggyr | *Pectinopygus* | *gyricornis* | *Phalacrocorax carbo* | Italy | SRR13389504 |

| Count_number | Sodalis_detection | Library_code | Louse_Genus | Louse_Species | Vertebrate_Host | Collection_Country | NCBI_SRA |
|---|---|---|---|---|---|---|---|
| 674 | Negative | Pgpun | *Pectinopygus* | *punctatus* | *Phalacrocorax punctatus* | New Zealand | SRR13389503 |
| 675 | Negative | Pgset | *Pectinopygus* | *setosus* | *Phalacrocorax sulcirostris* | Australia | SRR13389502 |
| 676 | Negative | PgspPhhar | *Pectinopygus* | *sp.* | *Nannopterum harrisi* | Ecuador | SRR17556636 |
| 677 | Negative | PgspSusul | *Pectinopygus* | *sp.* | *Sula sula* | United States of America | SRR17556635 |
| 678 | Negative | Pgtor | *Pectinopygus* | *tordoffi* | *Pelecanus erythrorhynchos* | United States of America | SRR8177131 |
| 679 | Negative | Pgtur | *Pectinopygus* | *turbinatus* | *Leucocarbo atriceps* | Argentina | SRR17556634 |
| 680 | Negative | Qdend | *Pelmatocerandra* | *enderleini* | *Pelecanoides georgicus* | Kerguelan Island | SRR5308184 |
| 681 | Negative | PnarcPitri | *Penenirmus* | *arcticus* | *Picoides tridactylus* | Russia | SRR9693840 |
| 682 | Negative | PnaurCopun | *Penenirmus* | *auritus* | *Colaptes punctigula* | Peru | SRR8566314 |
| 683 | Negative | PnaurDemaj | *Penenirmus* | *auritus* | *Dendrocopos major* | Russia | SRR9693839 |
| 684 | Negative | PnaurDegoe | *Penenirmus* | *auritus* | *Dendropicos goertae* | Ghana | SRR9693805 |
| 685 | Negative | PnaurPipub | *Penenirmus* | *auritus* | *Dryobates pubescens* | United States of America | SRR9693833 |
| 686 | Negative | PnaurDrpil | *Penenirmus* | *auritus* | *Dryocopus pileatus* | United States of America | SRR8566315 |
| 687 | Negative | PnaurMeaur | *Penenirmus* | *auritus* | *Melanerpes aurifrons* | United States of America | SRR9693837 |
| 688 | Negative | PnaurMecan | *Penenirmus* | *auritus* | *Melanerpes candidus* | Bolivia | SRR9693807 |
| 689 | Negative | PnaurMecru | *Penenirmus* | *auritus* | *Melanerpes cruentatus* | Peru | SRR9693836 |
| 690 | Negative | PnaurMeery | *Penenirmus* | *auritus* | *Melanerpes erythrocephalus* | United States of America | SRR8582579 |
| 691 | Negative | PnaurPifla | *Penenirmus* | *auritus* | *Piculus flavigula* | Brazil | SRR9693813 |
| 692 | Negative | PnaurPiaur | *Penenirmus* | *auritus* | *Picumnus aurifrons* | Brazil | SRR9693812 |
| 693 | Negative | PnaurSpnuc | *Penenirmus* | *auritus* | *Sphyrapicus nuchalis* | United States of America | SRR9693834 |
| 694 | Negative | PnaurSpvar1 | *Penenirmus* | *auritus* | *Sphyrapicus varius* | United States of America | SRR16574583 |
| 695 | Negative | PnguiLydub | *Penenirmus* | *guineensis* | *Lybius dubius* | Ghana | SRR9693804 |
| 696 | Negative | Pnjun | *Penenirmus* | *jungens* | *Colaptes auratus* | United States of America | SRR8566316 |
| 697 | Negative | Pnmar | *Penenirmus* | *marginatus* | *Indicator indicator* | Malawi | SRR8566317 |
| 698 | Negative | PnpicPican | *Penenirmus* | *pici* | *Picus canus* | Russia | SRR9693838 |
| 699 | Negative | PnspAnlin | *Penenirmus* | *sp.* | *Anthus lineiventris* | Malawi | SRR8566318 |
| 700 | Negative | PnspBrbab | *Penenirmus* | *sp.* | *Bradypterus baboecala* | Malawi | SRR8566319 |
| 701 | Negative | PnspCatur | *Penenirmus* | *sp.* | *Campylorhynchus turdinus* | Brazil | SRR8566322 |
| 702 | Negative | PnspCaaur | *Penenirmus* | *sp.* | *Capito auratus* | Peru | SRR9693811 |
| 703 | Negative | PnspCaaur2 | *Penenirmus* | *sp.* | *Capito aurovirens* | Peru | SRR9693835 |
| 704 | Negative | PnspCabru | *Penenirmus* | *sp.* | *Capito brunneipectus* | Brazil | SRR9693803 |
| 705 | Negative | PnspCeame | *Penenirmus* | *sp.* | *Certhia americana* | Mexico | SRR8566323 |
| 706 | Negative | PnspCiruf | *Penenirmus* | *sp.* | *Cisticola rufilatus* | Malawi | SRR8566324 |
| 707 | Negative | PnspDegri | *Penenirmus* | *sp.* | *Dendropicos griseocephalus* | Malawi | SRR8566325 |
| 708 | Negative | PnspVenig | *Penenirmus* | *sp.* | *Dryobates nigriceps* | Peru | SRR8582584 |
| 709 | Negative | PnspEuric | *Penenirmus* | *sp.* | *Eubucco richardsoni* | Brazil | SRR8566326 |
| 710 | Negative | PnspEuver | *Penenirmus* | *sp.* | *Eubucco versicolor* | Peru | SRR9693810 |
| 711 | Negative | PnspGycal | *Penenirmus* | *sp.* | *Gymnobucco calvus* | Ghana | SRR8145998 |

| Count_number | Sodalis_detection | Library_code | Louse_Genus | Louse_Species | Vertebrate_Host | Collection_Country | NCBI_SRA |
|---|---|---|---|---|---|---|---|
| 712 | Negative | PnspGypel | *Penenirmus* | *sp.* | *Gymnobucco peli* | Ghana | SRR9693841 |
| 713 | Negative | PnspInmin | *Penenirmus* | *sp.* | *Indicator minor* | Malawi | SRR8566327 |
| 714 | Negative | PnspInvar | *Penenirmus* | *sp.* | *Indicator variegatus* | Malawi | SRR8566328 |
| 715 | Negative | PnspInwil | *Penenirmus* | *sp.* | *Indicator willcocksi* | Ghana | SRR8173272 |
| 716 | Negative | PnspMerub | *Penenirmus* | *sp.* | *Melanerpes rubricapillus* | Panama | SRR9693819 |
| 717 | Negative | PnspPobil | *Penenirmus* | *sp.* | *Pogoniulus bilineatus* | Democratic Republic of Congo | SRR8582583 |
| 718 | Negative | PnspPsmin | *Penenirmus* | *sp.* | *Psaltriparus minimus* | Mexico | SRR8566330 |
| 719 | Negative | PnspMechr | *Penenirmus* | *sp.* | *Psilopogon chrysopogon* | Malaysia | SRR8582581 |
| 720 | Negative | BrspMefra | *Penenirmus* | *sp.* | *Psilopogon franklinii* | China | SRR14141047 |
| 721 | Negative | PnspLyleu | *Penenirmus* | *sp.* | *Tricholaema leucomelas* | South Africa | SRR8582580 |
| 722 | Negative | Pnzum | *Penenirmus* | *zumpti* | *Lybius torquatus* | South Africa | SRR9693832 |
| 723 | Negative | Prcir | *Perineus* | *circumfasciatus* | *Thalassarche melanophris* | United Kingdom | SRR5308177 |
| 724 | Negative | Wiabs1 | *Pessoiaella* | *absita* | *Opisthocomus hoazin* | Brazil | SRR16574576 |
| 725 | Negative | Ppage | *Philopterus* | *agelaii* | *Agelaius phoeniceus* | United States of America | SRR14887883 |
| 726 | Negative | Ppbio | *Philopterus* | *biocellatus* | *Struthidea cinerea* | Australia | SRR12762517 |
| 727 | Negative | Ppcor | *Philopterus* | *corvi* | *Corvus corax* | Canada | SRR12762516 |
| 728 | Negative | Ppino | *Philopterus* | *inopinatus* | *Corcorax melanorhamphos* | Australia | SRR12762515 |
| 729 | Negative | Ppkek | *Philopterus* | *kekilovae* | *Eremophila alpestris* | United States of America | SRR12762514 |
| 730 | Negative | Pporn | *Philopterus* | *ornatus* | *Oriolus larvatus* | Mozambique | SRR14887882 |
| 731 | Negative | Ppsol | *Philopterus* | *solus* | *Rhinopomastus cyanomelas* | Malawi | SRR14887880 |
| 732 | Negative | PpspAmmac | *Philopterus* | *sp.* | *Amblyornis macgregoriae* | New Guinea | SRR12762513 |
| 733 | Negative | PpspAshel | *Philopterus* | *sp.* | *Asthenes helleri* | Peru | SRR14141066 |
| 734 | Negative | PpspBamar | *Philopterus* | *sp.* | *Baryphthengus martii* | Panama | SRR8145999 |
| 735 | Negative | PhspBamo | *Philopterus* | *sp.* | *Batis molitor* | Malawi | SRR14887885 |
| 736 | Negative | PpspCagut | *Philopterus* | *sp.* | *Catharus guttatus* | United States of America | SRR14908127 |
| 737 | Negative | PpspPimen | *Philopterus* | *sp.* | *Ceratopipra mentalis* | Panama | SRR12762498 |
| 738 | Negative | PpspChjef | *Philopterus* | *sp.* | *Chlamydochaera jefferyi* | Malaysia | SRR14887879 |
| 739 | Negative | PpspCipun | *Philopterus* | *sp.* | *Cinclosoma punctatum* | Australia | SRR12762512 |
| 740 | Negative | PnspNear | *Philopterus* | *sp.* | *Cinnyris afer* | Malawi | SRR8566329 |
| 741 | Negative | PpspClpic | *Philopterus* | *sp.* | *Climacteris picumnus* | Australia | SRR12762511 |
| 742 | Negative | PpspColon | *Philopterus* | *sp.* | *Coracina longicauda* | New Guinea | SRR12762509 |
| 743 | Negative | Brste | *Philopterus* | *sp.* | *Corcorax melanorhamphos* | Australia | SRR8142486 |
| 744 | Negative | PpspColeu | *Philopterus* | *sp.* | *Cormobates leucophaea* | Australia | SRR12762510 |
| 745 | Negative | PpspCoorr | *Philopterus* | *sp.* | *Corvus orru* | Australia | SRR12762507 |
| 746 | Negative | PpspCrquo | *Philopterus* | *sp.* | *Cracticus quoyi* | Australia | SRR9693817 |
| 747 | Negative | PpspDiads | *Philopterus* | *sp.* | *Dicrurus adsimilis* | Ghana | SRR12762506 |
| 748 | Negative | PpspDileu | *Philopterus* | *sp.* | *Dicrurus leucophaeus* | China | SRR14141036 |
| 749 | Negative | PpspEurub | *Philopterus* | *sp.* | *Eugerygone rubra* | New Guinea | SRR12762505 |

| Count_number | Sodalis_detection | Library_code | Louse_Genus | Louse_Species | Vertebrate_Host | Collection_Country | NCBI_SRA |
|---|---|---|---|---|---|---|---|
| 750 | Negative | PpspEuann | *Philopterus* | *sp.* | *Euphonia anneae* | Panama | SRR9693818 |
| 751 | Negative | MaspGaal | *Philopterus* | *sp.* | *Galbula albirostris* | Brazil | SRR14887888 |
| 752 | Negative | PpspGrcya | *Philopterus* | *sp.* | *Grallina cyanoleuca* | Australia | SRR12762504 |
| 753 | Negative | PpspHakas | *Philopterus* | *sp.* | *Harpactes kasumba* | Malaysia | SRR8177132 |
| 754 | Negative | PpspHymad | *Philopterus* | *sp.* | *Hypsipetes madagascariensis* | Madagascar | SRR12762503 |
| 755 | Negative | MaspJaau | *Philopterus* | *sp.* | *Jacamerops aureus* | Brazil | SRR14887887 |
| 756 | Negative | PpspLaaet | *Philopterus* | *sp.* | *Laniarius aethiopicus* | Democratic Republic of Congo | SRR14908126 |
| 757 | Negative | PpspLecoe | *Philopterus* | *sp.* | *Lepidothrix coeruleocapilla* | Peru | SRR14141065 |
| 758 | Negative | PpspLopil | *Philopterus* | *sp.* | *Lophotriccus pileatus* | Panama | SRR12762502 |
| 759 | Negative | MaspMafu | *Philopterus* | *sp.* | *Malacoptila fusca* | Peru | SRR14908139 |
| 760 | Negative | PpspPanig | *Philopterus* | *sp.* | *Melaniparus niger* | Malawi | SRR14887877 |
| 761 | Negative | PpspMioli | *Philopterus* | *sp.* | *Mionectes olivaceus* | Panama | SRR12762501 |
| 762 | Negative | MaspMomo | *Philopterus* | *sp.* | *Monasa morphoeus* | Brazil | SRR14887886 |
| 763 | Negative | PpspMocla | *Philopterus* | *sp.* | *Motacilla clara* | Kenya | SRR14908125 |
| 764 | Negative | PpspMycol | *Philopterus* | *sp.* | *Myadestes coloratus* | Panama | SRR12762500 |
| 765 | Negative | PpspNobru | *Philopterus* | *sp.* | *Nonnula brunnea* | Peru | SRR14141064 |
| 766 | Negative | PpspNoruf | *Philopterus* | *sp.* | *Nonnula ruficapilla* | Brazil | SRR9693825 |
| 767 | Negative | TyspPamin | *Philopterus* | *sp.* | *Pachyramphus minor* | Brazil | SRR14908107 |
| 768 | Negative | PpspPagri | *Philopterus* | *sp.* | *Passer griseus* | Malawi | SRR14887878 |
| 769 | Negative | PpspPhict | *Philopterus* | *sp.* | *Phyllastrephus icterinus* | Ghana | SRR12762499 |
| 770 | Negative | PpspPiint | *Philopterus* | *sp.* | *Pipreola intermedia* | Peru | SRR14141063 |
| 771 | Negative | PpspPiang | *Philopterus* | *sp.* | *Pitta angolensis* | Malawi | SRR14887876 |
| 772 | Negative | PpspPiiri | *Philopterus* | *sp.* | *Pitta iris* | Australia | SRR9693824 |
| 773 | Negative | PpspPrnew | *Philopterus* | *sp.* | *Prionodura newtoniana* | Australia | SRR12762544 |
| 774 | Negative | PpspPrplu | *Philopterus* | *sp.* | *Prionops plumatus* | Malawi | SRR14887875 |
| 775 | Negative | PpspPtgui | *Philopterus* | *sp.* | *Ptiloprora guisei* | New Guinea | SRR12762543 |
| 776 | Negative | PpspPycin | *Philopterus* | *sp.* | *Pyrrhomyias cinnamomeus* | Peru | SRR14141070 |
| 777 | Negative | PhspQuqu | *Philopterus* | *sp.* | *Quelea quelea* | Malawi | SRR14887884 |
| 778 | Negative | PpspSaspi | *Philopterus* | *sp.* | *Salpornis spilonota* | Malawi | SRR14887874 |
| 779 | Negative | PpspSctur | *Philopterus* | *sp.* | *Schiffornis turdina* | Panama | SRR12762542 |
| 780 | Negative | PpspSyspo | *Philopterus* | *sp.* | *Silvicultrix frontalis* | Peru | SRR14141067 |
| 781 | Negative | PpspOcpul | *Philopterus* | *sp.* | *Silvicultrix pulchella* | Peru | SRR14141062 |
| 782 | Negative | PpspSppin | *Philopterus* | *sp.* | *Spinus pinus* | Canada | SRR12762541 |
| 783 | Negative | PpspSppus | *Philopterus* | *sp.* | *Spizella pusilla* | United States of America | SRR12762540 |
| 784 | Negative | PpspTenig | *Philopterus* | *sp.* | *Telophorus nigrifrons* | Malawi | SRR14887873 |
| 785 | Negative | PpspTeruf | *Philopterus* | *sp.* | *Terpsiphone rufiventer* | Ghana | SRR12762539 |
| 786 | Negative | PpspThpun | *Philopterus* | *sp.* | *Thamnophilus punctatus* | Brazil | SRR8177133 |
| 787 | Negative | PpspThruf | *Philopterus* | *sp.* | *Thlypopsis ruficeps* | Peru | SRR14141068 |

| Count_number | Sodalis_detection | Library_code | Louse_Genus | Louse_Species | Vertebrate_Host | Collection_Country | NCBI_SRA |
|---|---|---|---|---|---|---|---|
| 788 | Negative | PpspTosul | *Philopterus* | *sp.* | *Tolmomyias sulphurescens* | Panama | SRR12762538 |
| 789 | Negative | PpspTymel | *Philopterus* | *sp.* | *Tyrannus melancholicus* | Panama | SRR5308375 |
| 790 | Negative | PpspVojac | *Philopterus* | *sp.* | *Volatinia jacarina* | Panama | SRR12762537 |
| 791 | Negative | PpspHidau | *Philopterus* | *sp.* | *Cecropis daurica* | China | SRR8582587 |
| 792 | Negative | PpspDihot | *Philopterus* | *sp.* | *Dicrurus hottentottus* | China | SRR8582585 |
| 793 | Negative | PpspHepic | *Philopterus* | *sp.* | *Hemipus picatus* | China | SRR8582586 |
| 794 | Negative | Pweme | *Physconella* | *emersoni* | *Crypturellus undulatus* | Brazil | SRR13159342 |
| 795 | Negative | Pwnot | *Physconella* | *nothocercae* | *Nothocercus nigrocapillus* | Peru | SRR14908122 |
| 796 | Negative | PwspCrobs | *Physconella* | *sp.* | *Crypturellus obsoletus* | Brazil | SRR5308180 |
| 797 | Negative | PwspCrsou | *Physconella* | *sp.* | *Crypturellus soui* | Brazil | SRR14908118 |
| 798 | Negative | Phano2 | *Physconelloides* | *anolaimae 2* | *Patagioenas plumbea* | Brazil | SRR18820745 |
| 799 | Negative | CcspPealb | *Physconelloides* | *australiensis* | *Petrophassa albipennis* | Australia | SRR8145870 |
| 800 | Negative | Phcer1 | *Physconelloides* | *ceratoceps 1* | *Leptotila jamaicensis* | Mexico | SRR8173267 |
| 801 | Negative | Phcer4 | *Physconelloides* | *ceratoceps 4* | *Leptotila verreauxi* | Mexico | SRR18820744 |
| 802 | Negative | Phcer5 | *Physconelloides* | *ceratoceps 5* | *Leptotila rufaxilla* | Guyana | SRR8145993 |
| 803 | Negative | Phcub | *Physconelloides* | *cubanus* | *Geotrygon montana* | Mexico | SRR8173268 |
| 804 | Negative | Phper | *Physconelloides* | *perijae* | *Zentrygon frenata* | Peru | SRR8173269 |
| 805 | Negative | PhspGevrg | *Physconelloides* | *sp.* | *Leptotrygon veraguensis* | Panama | SRR17556633 |
| 806 | Negative | CcspPeruf | *Physconelloides* | *sp.* | *Petrophassa rufipennis* | Australia | SRR8145871 |
| 807 | Negative | PhspPhcha | *Physconelloides* | *sp.* | *Phaps chalcoptera* | Australia | SRR8173271 |
| 808 | Negative | CcspPhele | *Physconelloides* | *sp.* | *Phaps elegans* | Australia | SRR8145872 |
| 809 | Negative | CaspPhhis | *Physconelloides* | *sp.* | *Phaps histrionica* | Australia | SRR8172509 |
| 810 | Negative | Phspe1 | *Physconelloides* | *spenceri 1* | *Coada* | Brazil | SRR8173270 |
| 811 | Negative | Phwis | *Physconelloides* | *wisemani* | *Zenaida asiatica* | United States of America | SRR8145994 |
| 812 | Negative | Phzen2 | *Physconelloides* | *zenaidurae* | *Zenaida auriculata* | Argentina | SRR17556632 |
| 813 | Negative | Phzen | *Physconelloides* | *zenaidurae* | *Zenaida macroura* | United States of America | SRR8145995 |
| 814 | Negative | Pisno | *Picicola* | *snodgrassi* | *Melanerpes carolinus* | United States of America | SRR5308244 |
| 815 | Negative | PnspCaben | *Picicola* | *sp.* | *Campethera bennettii* | Malawi | SRR8566320 |
| 816 | Negative | Pisp.Cegra | *Picicola* | *sp.* | *Celeus grammicus* | Brazil | SRR5308245 |
| 817 | Negative | BrspPynig | *Picicola* | *sp.* | *Pycnonotus nigricans* | South Africa | SRR18820766 |
| 818 | Negative | Pisp.Pyalb | *Picicola* | *sp.* | *Pygarrhichas albogularis* | Argentina | SRR5308247 |
| 819 | Negative | Postr | *Podargoecus* | *strigoides* | *Podargus strigoides* | Australia | SRR8173273 |
| 820 | Negative | BrspGamae | *Priceiella* | *sp.* | *Garrulax maesi* | China | SRR14141069 |
| 821 | Negative | BrspNabre | *Priceiella* | *sp.* | *Gypsophila brevicaudata* | China | SRR14141037 |
| 822 | Negative | PrspTubre | *Priceiella* | *sp.* | *Gypsophila brevicaudata* | China | SRR18820741 |
| 823 | Negative | PrspLacin | *Priceiella* | *sp.* | *Ianthocincla cineracea* | China | SRR18820743 |
| 824 | Negative | BrspPoruf | *Priceiella* | *sp.* | *Pomatorhinus ruficollis* | China | SRR14141073 |
| 825 | Negative | PrspPoruf | *Priceiella* | *sp.* | *Pomatorhinus ruficollis* | China | SRR18820742 |

| Count_number | Sodalis_detection | Library_code | Louse_Genus | Louse_Species | Vertebrate_Host | Collection_Country | NCBI_SRA |
|---|---|---|---|---|---|---|---|
| 826 | Negative | BrspPagul | *Priceiella* | *sp.* | *Psittiparus gularis* | China | SRR14141078 |
| 827 | Negative | BrspStstr | *Priceiella* | *sp.* | *Stachyris strialata* | China | SRR14141058 |
| 828 | Negative | PrspGamil | *Priceiella* | *sp.* | *Trochalopteron milnei* | China | SRR8582588 |
| 829 | Negative | Psgra | *Pseudolipeurus* | *grandis* | *Nothocercus nigrocapillus* | Peru | SRR14908124 |
| 830 | Negative | Psmac | *Pseudolipeurus* | *macrogenitalis* | *Crypturellus undulatus* | Bolivia | SRR12762536 |
| 831 | Negative | Psplu | *Pseudolipeurus* | *plumbeus* | *Crypturellus tataupa* | Bolivia | SRR5308356 |
| 832 | Negative | Psplu1 | *Pseudolipeurus* | *plumbeus* | *Crypturellus tataupa* | Bolivia | SRR16574582 |
| 833 | Negative | PsspCrobs | *Pseudolipeurus* | *sp.* | *Crypturellus obsoletus* | Peru | SRR14908123 |
| 834 | Negative | PsspTigut | *Pseudolipeurus* | *sp.* | *Tinamus guttatus* | Brazil | SRR8173274 |
| 835 | Negative | PsspTimaj | *Pseudolipeurus* | *sp.* | *Tinamus major* | Brazil | SRR16503029 |
| 836 | Negative | Pulug | *Pseudonirmus* | *lugubris* | *Thalassoica antarctica* | New Zealand | SRR5308179 |
| 837 | Negative | Qsobs | *Pseudophilopterus* | *obseletus* | *Crypturellus obsoletus* | Brazil | SRR8173276 |
| 838 | Negative | QsspCrstr | *Pseudophilopterus* | *sp.* | *Crypturellus strigulosus* | Brazil | SRR5308185 |
| 839 | Negative | PcspOrarf | *Psittaconirmus* | *sp.* | *Oreopsittacus arfaki* | Papua New Guinea | SRR8173266 |
| 840 | Negative | PcspPsdes | *Psittaconirmus* | *sp.* | *Psittaculirostris desmarestii)* | Papua New Guinea | SRR8177128 |
| 841 | Negative | PcspTrrub | *Psittaconirmus* | *sp.* | *Trichoglossus rubritorquis* | Australia | SRR8177129 |
| 842 | Negative | QkeosCasan | *Psittoecus* | *eos* | *Cacatua sanguinea* | Australia | SRR8177135 |
| 843 | Negative | QkspCagal | *Psittoecus* | *sp.* | *Cacatua galerita* | Australia | SRR5308377 |
| 844 | Negative | QkspCalea | *Psittoecus* | *sp.* | *Cacatua leadbeateri* | Australia | SRR8173275 |
| 845 | Negative | Psfoe553 | *Psophiicola* | *foedus* | *Psophia crepitans* | Brazil | SRR8334273 |
| 846 | Negative | PsfoePsleu | *Psophiicola* | *foedus* | *Psophia leucoptera* | Brazil | SRR8177134 |
| 847 | Negative | PsspPscre | *Psophiicola* | *sp.* | *Psophia crepitans* | Brazil | SRR8566332 |
| 848 | Negative | PsspPsvir | *Psophiicola* | *sp.* | *Psophia viridis* | Brazil | SRR5308178 |
| 849 | Negative | Ptab | *Pterocotes* | *aberrans* | *Tinamus major* | Brazil | SRR16503028 |
| 850 | Negative | QrspCrund | *Pterocotes* | *sp.* | *Crypturellus undulatus* | Brazil | SRR8146004 |
| 851 | Negative | PtspTimaj | *Pterocotes* | *sp.* | *Tinamus major* | Brazil | SRR8566333 |
| 852 | Negative | Quaur | *Quadraceps* | *auratus* | *Haematopus ostralegus* | Australia | SRR13159318 |
| 853 | Negative | Qubic | *Quadraceps* | *bicuspus* | *Charadrius dubius* | Romania | SRR18820740 |
| 854 | Negative | Quboo | *Quadraceps* | *boophilus* | *Charadrius vociferus* | Canada | SRR8173279 |
| 855 | Negative | Qucha | *Quadraceps* | *charadrii* | *Pluvialis apricaria* | Sweden | SRR18820739 |
| 856 | Negative | Qudre | *Quadraceps* | *dressleri* | *Elseyornis melanops* | Australia | SRR13159323 |
| 857 | Negative | Lnhae | *Quadraceps* | *haematopi* | *Haematopus ostralegus* | Australia | SRR13159317 |
| 858 | Negative | Quhia | *Quadraceps* | *hiaticulae* | *Charadrius hiaticula* | Sweden | SRR18820738 |
| 859 | Negative | Qujun | *Quadraceps* | *junceus* | *Vanellus vanellus* | Italy | SRR18820736 |
| 860 | Negative | Qukos | *Quadraceps* | *kosswigi* | *Cladorhynchus leucocephalus* | Australia | SRR13159330 |
| 861 | Negative | Qunyc | *Quadraceps* | *nycthemerus* | *Sternula nereis* | Australia | SRR13159315 |
| 862 | Negative | Quobl | *Quadraceps* | *obliquus* | *Uria aalge* | Sweden | SRR18820734 |
| 863 | Negative | Quobs | *Quadraceps* | *obscurus* | *Tringa glareola* | Romania | SRR18820733 |

| Count_number | Sodalis_detection | Library_code | Louse_Genus | Louse_Species | Vertebrate_Host | Collection_Country | NCBI_SRA |
|---|---|---|---|---|---|---|---|
| 864 | Negative | Quorn | *Quadraceps* | *ornatus* | *Larus argentatus* | Canada | SRR13389497 |
| 865 | Negative | QupunLanov | *Quadraceps* | *punctatus* | *Chroicocephalus novaehollandiae* | Australia | SRR13159316 |
| 866 | Negative | Qupun1 | *Quadraceps* | *punctatus* | *Larus argentatus* | Canada | SRR16574581 |
| 867 | Negative | Qupun | *Quadraceps* | *punctatus* | *Larus argentatus* | Canada | SRR5308139 |
| 868 | Negative | QupunLepip | *Quadraceps* | *punctatus* | *Leucophaeus pipixcan* | Canada | SRR13389496 |
| 869 | Negative | Qurav | *Quadraceps* | *ravus* | *Actitis hypoleucos* | Japan | SRR18820732 |
| 870 | Negative | Zirec | *Quadraceps* | *recurvirostrae* | *Recurvirostra novaehollandiae* | Australia | SRR13159329 |
| 871 | Negative | QuspBugra | *Quadraceps* | *sp.* | *Burhinus grallarius* | Japan | SRR13159322 |
| 872 | Negative | QuspThrub | *Quadraceps* | *sp.* | *Charadrius cucullatus* | Australia | SRR8173284 |
| 873 | Negative | QuspEsmag | *Quadraceps* | *sp.* | *Esacus magnirostris* | Australia | SRR8173281 |
| 874 | Negative | QuspHaful | *Quadraceps* | *sp.* | *Haematopus fuliginosus* | Australia | SRR8146005 |
| 875 | Negative | QuspHimex | *Quadraceps* | *sp.* | *Himantopus mexicanus* | United States of America | SRR8173282 |
| 876 | Negative | QuspLapac | *Quadraceps* | *sp.* | *Larus pacificus* | Australia | SRR13159326 |
| 877 | Negative | QuspPlful | *Quadraceps* | *sp.* | *Pluvialis fulva* | Australia | SRR18820730 |
| 878 | Negative | QuspReame | *Quadraceps* | *sp.* | *Recurvirostra americana* | United States of America | SRR8173283 |
| 879 | Negative | QuspCucha | *Quadraceps* | *sp.* | *Rhinoptilus chalcopterus* | Ghana | SRR8173280 |
| 880 | Negative | QuspStisa | *Quadraceps* | *sp.* | *Stiltia isabella* | Australia | SRR5809353 |
| 881 | Negative | QuspStisa2 | *Quadraceps* | *sp.* | *Stiltia isabella* | Australia | SRR13389495 |
| 882 | Negative | QuspStber | *Quadraceps* | *sp.* | *Thalasseus bergii)* | Australia | SRR13159325 |
| 883 | Negative | QuspThorb | *Quadraceps* | *sp.* | *Thinocorus orbignyianus* | Bolivia | SRR13159313 |
| 884 | Negative | QuspVamil | *Quadraceps* | *sp.* | *Vanellus miles* | Australia | SRR8173285 |
| 885 | Negative | QuspRenov1 | *Quadraceps* | *sp. 1* | *Recurvirostra novaehollandiae* | Australia | SRR13159328 |
| 886 | Negative | Qustr | *Quadraceps* | *strepsilaris* | *Arenaria interpres* | Australia | SRR8173286 |
| 887 | Negative | Raadv | *Rallicola* | *advenus* | *Fulica americana* | United States of America | SRR5308186 |
| 888 | Negative | Racal | *Rallicola* | *californicus* | *Rallus longirostris* | United States of America | SRR8566335 |
| 889 | Negative | Racep127 | *Rallicola* | *cephalosa* | *Glyphorynchus spirurus* | Panama | SRR8173287 |
| 890 | Negative | Racer | *Rallicola* | *certhia* | *Dendrocolaptes platyrostris* | Panama | SRR8177137 |
| 891 | Negative | Racor | *Rallicola* | *cornutae* | *Fulica cornuta* | Argentina | SRR8173288 |
| 892 | Negative | RacunXigut | *Rallicola* | *cunchotambo* | *Xiphorhynchus guttatus* | Brazil | SRR8566336 |
| 893 | Negative | RacunXioce | *Rallicola* | *cunchotambo* | *Xiphorhynchus ocellatus* | Brazil | SRR8566337 |
| 894 | Negative | RacunXitri | *Rallicola* | *cunchotambo* | *Xiphorhynchus triangularis* | Peru | SRR8334275 |
| 895 | Negative | Radec | *Rallicola* | *deckeri* | *Dendrocincla homochroa* | Panama | SRR8173289 |
| 896 | Negative | Raell | *Rallicola* | *elliotti* | *Porphyrio martinica* | United States of America | SRR8566338 |
| 897 | Negative | RafulFuatr | *Rallicola* | *fulicae* | *Fulica atra* | Australia | SRR8173290 |
| 898 | Negative | RafulFucri | *Rallicola* | *fulicae* | *Fulica cristata* | South Africa | SRR8146007 |
| 899 | Negative | Ragra | *Rallicola* | *gracilentus* | *Apteryx haastii* | New Zealand | SRR9693796 |
| 900 | Negative | Ragui | *Rallicola* | *guimaraesi* | *Fulica rufifrons* | Argentina | SRR8173291 |
| 901 | Negative | Rahar42 | *Rallicola* | *harrisoni* | *Gallirallus australis* | New Zealand | SRR9693799 |

| Count_number | Sodalis_detection | Library_code | Louse_Genus | Louse_Species | Vertebrate_Host | Collection_Country | NCBI_SRA |
|---|---|---|---|---|---|---|---|
| 902 | Negative | Rahar10 | *Rallicola* | *harrisoni* | *Gallirallus australis hectori* | New Zealand | SRR8582590 |
| 903 | Negative | RaharDesti | *Rallicola* | *harveyi* | *Certhiasomus stictolaemus* | Brazil | SRR8566339 |
| 904 | Negative | Rains | *Rallicola* | *insularis* | *Corvus kubaryi* | Marianas Islands | SRR9693801 |
| 905 | Negative | Rakey | *Rallicola* | *keymerae* | *Dendrexetastes rufigula* | Brazil | SRR8566340 |
| 906 | Negative | RalacXifla | *Rallicola* | *lachrymosa* | *Xiphorhynchus flavigaster* | Nicaragua | SRR8177139 |
| 907 | Negative | Ralac | *Rallicola* | *lachrymosa* | *Xiphorhynchus lachrymosus* | Panama | SRR8146008 |
| 908 | Negative | RalatSyrut | *Rallicola* | *laticephala* | *Synallaxis rutilans* | Brazil | SRR8566341 |
| 909 | Negative | Raleu | *Rallicola* | *leucopterae* | *Fulica leucoptera* | Argentina | SRR8173292 |
| 910 | Negative | Rapic | *Rallicola* | *picirostris* | *Dendrocolaptes picumnus* | Brazil | SRR8566342 |
| 911 | Negative | Raruf | *Rallicola* | *rufigularis* | *Sclerurus rufigularis* | Brazil | SRR8566343 |
| 912 | Negative | Rasca | *Rallicola* | *scapanoides* | *Campephilus melanoleucos* | Panama | SRR9693820 |
| 913 | Negative | RaspAnstr | *Rallicola* | *sp.* | *Anabacerthia striaticollis* | Peru | SRR8334276 |
| 914 | Negative | FospAnvar | *Rallicola* | *sp.* | *Anabacerthia variegaticeps* | Panama | SRR5308159 |
| 915 | Negative | RaspAnvar | *Rallicola* | *sp.* | *Anabacerthia variegaticeps* | Panama | SRR8173293 |
| 916 | Negative | RaspApaus | *Rallicola* | *sp.* | *Apteryx australis* | New Zealand | SRR9693794 |
| 917 | Negative | RaspApowe | *Rallicola* | *sp.* | *Apteryx owenii* | New Zealand | SRR9693795 |
| 918 | Negative | RaspApsp | *Rallicola* | *sp.* | *Apteryx sp.* | New Zealand | SRR5308364 |
| 919 | Negative | RaspAuinf | *Rallicola* | *sp.* | *Automolus infuscatus* | Brazil | SRR8566344 |
| 920 | Negative | RaspAuoch | *Rallicola* | *sp.* | *Automolus ochrolaemus* | Brazil | SRR8566345 |
| 921 | Negative | RaspCapus | *Rallicola* | *sp.* | *Campylorhamphus pusillus* | Peru | SRR8566346 |
| 922 | Negative | FmspCorob | *Rallicola* | *sp.* | *Conopophaga roberti* | Brazil | SRR8334248 |
| 923 | Negative | FmspCylin | *Rallicola* | *sp.* | *Cymbilaimus lineatus* | Brazil | SRR8582569 |
| 924 | Negative | RaspDeful | *Rallicola* | *sp.* | *Dendrocincla fuliginosa* | Panama | SRR8173295 |
| 925 | Negative | RaspDemer | *Rallicola* | *sp.* | *Dendrocincla merula* | Brazil | SRR8566347 |
| 926 | Negative | RaspDecer | *Rallicola* | *sp.* | *Dendrocolaptes certhia* | Panama | SRR8173294 |
| 927 | Negative | RaspFuala | *Rallicola* | *sp.* | *Fulica alai* | United States of America | SRR8173297 |
| 928 | Negative | RaspFuard | *Rallicola* | *sp.* | *Fulica ardesiaca* | Argentina | SRR8173298 |
| 929 | Negative | RaspGagal | *Rallicola* | *sp.* | *Gallinula galeata sandvicensis* | United States of America | SRR8173299 |
| 930 | Negative | RaspEucas | *Rallicola* | *sp.* | *Gallirallus castaneoventris* | Australia | SRR8173296 |
| 931 | Negative | RaspGaphi | *Rallicola* | *sp.* | *Gallirallus philippensis* | New Zealand | SRR9693797 |
| 932 | Negative | RaspHybri | *Rallicola* | *sp.* | *Hylexetastes uniformis* | Brazil | SRR8177140 |
| 933 | Negative | RaspLamel | *Rallicola* | *sp.* | *Laterallus melanophaius* | Brazil | SRR8566348 |
| 934 | Negative | RaspLeaff | *Rallicola* | *sp.* | *Lepidocolaptes affinis* | Peru | SRR8566349 |
| 935 | Negative | RaspLealb | *Rallicola* | *sp.* | *Lepidocolaptes albolineatus* | Brazil | SRR8566350 |
| 936 | Negative | RaspMasqu | *Rallicola* | *sp.* | *Margarornis squamiger* | Peru | SRR8566351 |
| 937 | Negative | RaspNalon | *Rallicola* | *sp.* | *Nasica longirostris* | Brazil | SRR8177141 |
| 938 | Negative | MaspOrnig | *Rallicola* | *sp.* | *Oriolus nigripennis* | Ghana | SRR18820749 |
| 939 | Negative | PnspThgen | *Rallicola* | *sp.* | *Pheugopedius genibarbis* | Brazil | SRR8566331 |

| Count_number | Sodalis_detection | Library_code | Louse_Genus | Louse_Species | Vertebrate_Host | Collection_Country | NCBI_SRA |
|---|---|---|---|---|---|---|---|
| 940 | Negative | RaspPhery | *Rallicola* | *sp.* | *Philydor erythrocercum* | Brazil | SRR8566352 |
| 941 | Negative | RaspPhpyr | *Rallicola* | *sp.* | *Philydor pyrrhodes* | Brazil | SRR8566353 |
| 942 | Negative | RaspPopor1 | *Rallicola* | *sp.* | *Porphyrio melanotus* | New Zealand | SRR8582591 |
| 943 | Negative | RaspPopor10 | *Rallicola* | *sp.* | *Porphyrio melanotus* | New Zealand | SRR8582592 |
| 944 | Negative | RaspPopor33 | *Rallicola* | *sp.* | *Porphyrio melanotus* | New Zealand | SRR9693798 |
| 945 | Negative | RaspPopor | *Rallicola* | *sp.* | *Porphyrio porphyrio* | Australia | SRR8173300 |
| 946 | Negative | RaspPocar | *Rallicola* | *sp.* | *Porzana carolina* | United States of America | SRR8566354 |
| 947 | Negative | RaspPrbru | *Rallicola* | *sp.* | *Premnoplex brunnescens* | Peru | SRR8173301 |
| 948 | Negative | RaspPrbru | *Rallicola* | *sp.* | *Premnoplex brunnescens* | Panama | SRR8173301 |
| 949 | Negative | RaspRaele | *Rallicola* | *sp.* | *Rallus elegans* | United States of America | SRR8566355 |
| 950 | Negative | RaspSaruf | *Rallicola* | *sp.* | *Sarothrura rufa rufa* | Malawi | SRR8566356 |
| 951 | Negative | RaspScruf | *Rallicola* | *sp.* | *Sclerurus rufigularis* | Brazil | SRR8566357 |
| 952 | Negative | RaspScsp | *Rallicola* | *sp.* | *Scytalopus sp.* | Peru | SRR8566358 |
| 953 | Negative | RaspSyruf | *Rallicola* | *sp.* | *Syndactyla rufosuperciliata* | Peru | SRR8177143 |
| 954 | Negative | RaspSysub | *Rallicola* | *sp.* | *Syndactyla subalaris* | Panama | SRR8173302 |
| 955 | Negative | RaspXerut | *Rallicola* | *sp.* | *Xenops rutilans* | Peru | SRR8334277 |
| 956 | Negative | RaspXiery | *Rallicola* | *sp.* | *Xiphorhynchus erythropygius* | Panama | SRR5308189 |
| 957 | Negative | RaspXiobs | *Rallicola* | *sp.* | *Xiphorhynchus obsoletus* | Brazil | SRR8177144 |
| 958 | Negative | Ratak102 | *Rallicola* | *takahe* | *Porphyrio hochstetteri* | New Zealand | SRR8582593 |
| 959 | Negative | Ratak34 | *Rallicola* | *takahe* | *Porphyrio hochstetteri* | New Zealand | SRR9693800 |
| 960 | Negative | Ratak93 | *Rallicola* | *takahe* | *Porphyrio hochstetteri* | New Zealand | SRR8582594 |
| 961 | Negative | Ratom | *Rallicola* | *tomkinsi* | *Sclerurus caudacutus* | Brazil | SRR8566359 |
| 962 | Negative | Rawer | *Rallicola* | *wernecki* | *Fulica armillata* | Argentina | SRR8173304 |
| 963 | Negative | Rpabb | *Rhopaloceras* | *abbreviatus* | *Nothocercus nigrocapillus* | Peru | SRR14908114 |
| 964 | Negative | Rpbre | *Rhopaloceras* | *brevitemporalis* | *Crypturellus obsoletus* | Brazil | SRR13159335 |
| 965 | Negative | RpbrePE | *Rhopaloceras* | *brevitemporalis* | *Crypturellus obsoletus* | Peru | SRR14908113 |
| 966 | Negative | Rppen | *Rhopaloceras* | *pennaticeps* | *Crypturellus parvirostris* | Bolivia | SRR13159300 |
| 967 | Negative | Rprud | *Rhopaloceras* | *rudimentarius* | *Crypturellus soui* | Brazil | SRR14908112 |
| 968 | Negative | RpspCrstr | *Rhopaloceras* | *sp.* | *Crypturellus strigulosus* | Brazil | SRR5308190 |
| 969 | Negative | Rpund | *Rhopaloceras* | *undulatus* | *Crypturellus undulatus* | Bolivia | SRR12762533 |
| 970 | Negative | Rhand | *Rhynchotura* | *andina* | *Tinamotis pentlandii* | Peru | SRR16503026 |
| 971 | Negative | Rhsex | *Rhynchotura* | *sexpunctata* | *Rhynchotus rufescens* | Argentina | SRR16503025 |
| 972 | Negative | Rhpar | *Rhynonirmus* | *parsonsae* | *Scolopax minor* | United States of America | SRR5308252 |
| 973 | Negative | Sahae | *Saemundssonia* | *haematopi* | *Haematopus fuliginosus* | Australia | SRR8146013 |
| 974 | Negative | Sahae2 | *Saemundssonia* | *haematopi* | *Haematopus ostralegus* | Australia | SRR13159319 |
| 975 | Negative | Salar | *Saemundssonia* | *lari* | *Chroicocephalus novaehollandiae* | Australia | SRR5308141 |
| 976 | Negative | Salar1 | *Saemundssonia* | *lari* | *Chroicocephalus novaehollandiae* | Australia | SRR16574580 |
| 977 | Negative | Salat | *Saemundssonia* | *laticandata* | *Hydroprogne caspia* | Canada | SRR13389494 |

| Count_number | Sodalis_detection | Library_code | Louse_Genus | Louse_Species | Vertebrate_Host | Collection_Country | NCBI_SRA |
|---|---|---|---|---|---|---|---|
| 978 | Negative | Samel | *Saemundssonia* | *melanocephalus* | *Sternula nereis)* | Australia | SRR13159314 |
| 979 | Negative | Saseg | *Saemundssonia* | *segulata* | *Antigone canadensis* | Canada | SRR12762532 |
| 980 | Negative | CjspNumin | *Saemundssonia* | *sp.* | *Numenius minutus* | Australia | SRR8145882 |
| 981 | Negative | CjspNupha | *Saemundssonia* | *sp.* | *Numenius phaeopus* | Australia | SRR8145883 |
| 982 | Negative | SaspStber | *Saemundssonia* | *sp.* | *Thalasseus bergii* | Australia | SRR13159324 |
| 983 | Negative | SpspTapor | *Splendoroffula* | *sp.* | *Gallirex porphyreolophus* | South Africa | SRR5308378 |
| 984 | Negative | SpspTusch | *Splendoroffula* | *sp.* | *Tauraco schalowi* | Malawi | SRR13389493 |
| 985 | Negative | Stcur | *Strigiphilus* | *cursor* | *Asio flammeus* | Canada | SRR5308191 |
| 986 | Negative | StspNicon | *Strigiphilus* | *sp.* | *Ninox connivens* | Australia | SRR8146016 |
| 987 | Negative | StspTyalb | *Strigiphilus* | *sp.* | *Tyto alba* | United States of America | SRR16574579 |
| 988 | Negative | Sgcor | *Strongylocotes* | *cordiceps* | *Tinamus major* | Brazil | SRR8582596 |
| 989 | Negative | Sggla | *Strongylocotes* | *glabrous* | *Crypturellus tataupa* | Bolivia | SRR8582597 |
| 990 | Negative | Sglim | *Strongylocotes* | *limai* | *Crypturellus undulatus* | Brazil | SRR8582598 |
| 991 | Negative | SgorbCrpar | *Strongylocotes* | *orbicularis* | *Crypturellus parvirostris* | Bolivia | SRR8582599 |
| 992 | Negative | SgspCrstr | *Strongylocotes* | *sp.* | *Crypturellus strigulosus* | Brazil | SRR8146015 |
| 993 | Negative | StspTigut | *Strongylocotes* | *sp.* | *Tinamus guttatus* | Brazil | SRR16503022 |
| 994 | Negative | SgspTigut | *Strongylocotes* | *sp.* | *Tinamus guttatus* | Brazil | SRR8580043 |
| 995 | Negative | StspTimaj | *Strongylocotes* | *sp.* | *Tinamus major* | Brazil | SRR16503021 |
| 996 | Negative | Slstr | *Struthilipeurus* | *struthionis* | *Struthio camelus* | United Kingdom | SRR5308365 |
| 997 | Negative | SlspRhame | *Struthiolipeurus* | *sp.* | *Rhea americana* | Brazil | SRR5308383 |
| 998 | Negative | Snsp | *Sturnidoecus* | *sp.* | *Lamprotornis purpureus* | Ghana | SRR5308357 |
| 999 | Negative | Sncal | *Sturnidoecus* | *caligineus* | *Turdus grayi* | Panama | SRR18820729 |
| 1000 | Negative | Snsim | *Sturnidoecus* | *simplex* | *Turdus migratorius* | Canada | SRR18820728 |
| 1001 | Negative | Tirot | *Tinamicola* | *rotundatus* | *Rhynchotus rufescens* | Argentina | SRR16503020 |
| 1002 | Negative | Tiand | *Tinamotaecola* | *andinae* | *Tinamotis pentlandii* | Peru | SRR8334278 |
| 1003 | Negative | TispEufor | *Tinamotaecola* | *sp.* | *Eudromia formosa* | Argentina | SRR13159332 |
| 1004 | Negative | TispTipen | *Tinamotaecola* | *sp.* | *Tinamotis pentlandii* | Peru | SRR16503019 |
| 1005 | Negative | Tbhex | *Trabeculus* | *hexakon* | *Pterodroma hypoleuca* | United States of America | SRR5308192 |
| 1006 | Negative | BrspMevir | *Traihoriella* | *sp.* | *Megalaima virens* | China | SRR8582563 |
| 1007 | Negative | PnspMemon | *Traihoriella* | *sp.* | *Psilopogon monticola* | Borneo | SRR9693806 |
| 1008 | Negative | PnspMemys | *Traihoriella* | *sp.* | *Psilopogon mystacophanos* | Borneo | SRR9693809 |
| 1009 | Negative | BrspMemys | *Traihoriella* | *sp.* | *Psilopogon mystacophanos* | Borneo | SRR8142485 |
| 1010 | Negative | PnspMevir | *Traihoriella* | *sp.* | *Psilopogon virens* | China | SRR9693821 |
| 1011 | Negative | Trspi | *Trichodopeostus* | *spinosus* | *Nothocercus nigrocapillus* | Peru | SRR8334279 |
| 1012 | Negative | TrspiPE | *Trichodopeostus* | *spinosus* | *Nothocercus nigrocapillus* | Peru | SRR14908108 |
| 1013 | Negative | Tfbab2 | *Trichophilopterus* | *babakotophilus* | *Propithecus verreauxi* | Madagascar | SRR16574577 |
| 1014 | Negative | TfbabPrver | *Trichophilopterus* | *babakotophilus* | *Propithecus verreauxi* | Madagascar | SRR5308144 |
| 1015 | Negative | Tkang | *Turnicola* | *angustissimus* | *Turnix nigricollis* | Madagascar | SRR8146018 |

| Count_number | Sodalis_detection | Library_code | Louse_Genus | Louse_Species | Vertebrate_Host | Collection_Country | NCBI_SRA |
|---|---|---|---|---|---|---|---|
| 1016 | Negative | TkspTupyr | *Turnicola* | *sp.* | *Turnix pyrrhothorax* | Australia | SRR5308379 |
| 1017 | Negative | TkspTuvar | *Turnicola* | *sp.* | *Turnix varius* | Australia | SRR8146019 |
| 1018 | Negative | Upupu | *Upupicola* | *upupae* | *Upupa africana* | Malawi | SRR5308258 |
| 1019 | Negative | Veber | *Vernoniella* | *bergi* | *Guira guira* | Brazil | SRR14141049 |
| 1020 | Negative | Vegui | *Vernoniella* | *guimaraesi* | *Crotophaga ani* | Panama | SRR5308380 |
| outgroup | Positive | Pfluc4 | *Proechinophthirus* | *fluctus* | *Callorhinus ursinus* | United States of America | SRR2013546 |

**Table S2.** Summary of *Sodalis* detection in louse samples by genus. This table shows the counts and percentages of positive and negative samples for each louse genus analyzed. It includes the total number of samples, as well as the proportion of samples that were detected positive or negative for *Sodalis*.

| Louse_Genus | Total_n | Sodalis_detection | | Percentage_Negative | Percentage_Positive |
|---|---|---|---|---|---|
| | | Negative | Positive | | |
| *Acidoproctus* | 3 | 3 | 0 | 100 | 0 |
| *Acronirmus* | 1 | 1 | 0 | 100 | 0 |
| *Acutifrons* | 1 | 1 | 0 | 100 | 0 |
| *Alcedoecus* | 2 | 2 | 0 | 100 | 0 |
| *Alcedoffula* | 10 | 5 | 5 | 50 | 50 |
| *Anaticola* | 29 | 23 | 6 | 79,310345 | 20,6896552 |
| *Anatoecus* | 10 | 10 | 0 | 100 | 0 |
| *Aquanirmus* | 6 | 6 | 0 | 100 | 0 |
| *Ardeicola* | 14 | 10 | 4 | 71,428571 | 28,5714286 |
| *Ardeiphagus* | 2 | 2 | 0 | 100 | 0 |
| *Auricotes* | 6 | 6 | 0 | 100 | 0 |
| *Austrogoniodes* | 1 | 0 | 1 | 0 | 100 |
| *Austrokelloggia* | 1 | 1 | 0 | 100 | 0 |
| *Austrophilopterus* | 6 | 3 | 3 | 50 | 50 |
| *Bedfordiella* | 1 | 1 | 0 | 100 | 0 |
| *Bizarrifrons* | 1 | 0 | 1 | 0 | 100 |
| *Bothriometopus* | 1 | 1 | 0 | 100 | 0 |
| *Brueelia* | 26 | 3 | 23 | 11,538462 | 88,4615385 |
| *Bucerocophorus* | 1 | 0 | 1 | 0 | 100 |
| *Buceroemersonia* | 2 | 1 | 1 | 50 | 50 |
| *Buceronirmus* | 1 | 0 | 1 | 0 | 100 |
| *Buerelius* | 1 | 1 | 0 | 100 | 0 |
| *Campanulotes* | 12 | 12 | 0 | 100 | 0 |
| *Capraiella* | 3 | 0 | 3 | 0 | 100 |
| *Caracaricola* | 1 | 0 | 1 | 0 | 100 |

| Louse_Genus | Total_n | Sodalis_detection | | Percentage_Negative | Percentage_Positive |
|---|---|---|---|---|---|
| | | Negative | Positive | | |
| *Carduiceps* | 2 | 2 | 0 | 100 | 0 |
| *Chelopistes* | 3 | 3 | 0 | 100 | 0 |
| *Cincloecus* | 1 | 1 | 0 | 100 | 0 |
| *Cirropthirius* | 1 | 0 | 1 | 0 | 100 |
| *Clayiella* | 1 | 1 | 0 | 100 | 0 |
| *Colilipeurus* | 3 | 3 | 0 | 100 | 0 |
| *Colinicola* | 2 | 2 | 0 | 100 | 0 |
| *Coloceras* | 34 | 34 | 0 | 100 | 0 |
| *Columbicola* | 78 | 40 | 38 | 51,282051 | 48,7179487 |
| *Corvonirmus* | 2 | 2 | 0 | 100 | 0 |
| *Cotingacola* | 5 | 2 | 3 | 40 | 60 |
| *Craspedorrhynchus* | 4 | 4 | 0 | 100 | 0 |
| *Cuclotocephalus* | 5 | 1 | 4 | 20 | 80 |
| *Cuclotogaster* | 3 | 3 | 0 | 100 | 0 |
| *Cuculicola* | 6 | 1 | 5 | 16,666667 | 83,3333333 |
| *Cuculoecus* | 2 | 2 | 0 | 100 | 0 |
| *Dahlemhornia* | 3 | 3 | 0 | 100 | 0 |
| *Degeeriella* | 6 | 1 | 5 | 16,666667 | 83,3333333 |
| *Dicruobates* | 1 | 0 | 1 | 0 | 100 |
| *Discocorpus* | 4 | 4 | 0 | 100 | 0 |
| *Docophorocotes* | 1 | 1 | 0 | 100 | 0 |
| *Docophoroides* | 1 | 1 | 0 | 100 | 0 |
| *Echinophilopterus* | 2 | 2 | 0 | 100 | 0 |
| *Emersoniella* | 2 | 2 | 0 | 100 | 0 |
| *Epipicus* | 1 | 1 | 0 | 100 | 0 |
| *Episbates* | 1 | 1 | 0 | 100 | 0 |
| *Esthiopterum* | 1 | 1 | 0 | 100 | 0 |
| *Falcolipeurus* | 3 | 3 | 0 | 100 | 0 |
| *Forficuloecus* | 11 | 9 | 2 | 81,818182 | 18,1818182 |
| *Formicaphagus* | 11 | 1 | 10 | 9,0909091 | 90,9090909 |
| *Formicaricola* | 2 | 1 | 1 | 50 | 50 |
| *Fulicoffula* | 10 | 10 | 0 | 100 | 0 |
| *Furnariphilus* | 3 | 2 | 1 | 66,666667 | 33,3333333 |
| *Goniocotes* | 6 | 5 | 1 | 83,333333 | 16,6666667 |
| *Goniodes* | 14 | 14 | 0 | 100 | 0 |
| *Gonoides* | 1 | 1 | 0 | 100 | 0 |
| *Guimaraesiella* | 47 | 27 | 20 | 57,446809 | 42,5531915 |

| Louse_Genus | Total_n | Sodalis_detection | | Percentage_Negative | Percentage_Positive |
|---|---|---|---|---|---|
| | | Negative | Positive | | |
| *Haffneria* | 1 | 1 | 0 | 100 | 0 |
| *Halipeurus* | 2 | 2 | 0 | 100 | 0 |
| *Harrisoniella* | 1 | 1 | 0 | 100 | 0 |
| *Hecatrishula* | 1 | 0 | 1 | 0 | 100 |
| *Heptapsogaster* | 30 | 30 | 0 | 100 | 0 |
| *Heptapsus* | 2 | 2 | 0 | 100 | 0 |
| *Heptarthrogaster* | 2 | 2 | 0 | 100 | 0 |
| *Heptathrogaster* | 1 | 1 | 0 | 100 | 0 |
| *Hopkinsiella* | 1 | 1 | 0 | 100 | 0 |
| *Hypocrypturellus* | 3 | 3 | 0 | 100 | 0 |
| *Hyprocrypturellus* | 2 | 2 | 0 | 100 | 0 |
| *Ibidoecus* | 5 | 4 | 1 | 80 | 20 |
| *Incidifrons* | 2 | 2 | 0 | 100 | 0 |
| *Indoceoplanetes* | 4 | 1 | 3 | 25 | 75 |
| *Ischnocera* | 1 | 1 | 0 | 100 | 0 |
| *Kelloggia* | 5 | 4 | 1 | 80 | 20 |
| *Kodocephalon* | 2 | 2 | 0 | 100 | 0 |
| *Labicotes* | 1 | 1 | 0 | 100 | 0 |
| *Lagopoecus* | 2 | 2 | 0 | 100 | 0 |
| *Lamprocorpus* | 3 | 3 | 0 | 100 | 0 |
| *Lipeurus* | 5 | 5 | 0 | 100 | 0 |
| *Lunaceps* | 3 | 3 | 0 | 100 | 0 |
| *Maculinirmus* | 3 | 1 | 2 | 33,333333 | 66,6666667 |
| *Megaginus* | 4 | 4 | 0 | 100 | 0 |
| *Megapeostus* | 9 | 7 | 2 | 77,777778 | 22,2222222 |
| *Meropoecus* | 1 | 1 | 0 | 100 | 0 |
| *Meropsiella* | 2 | 2 | 0 | 100 | 0 |
| *Mirandofures* | 2 | 2 | 0 | 100 | 0 |
| *Motmotnirmus* | 1 | 1 | 0 | 100 | 0 |
| *Mulcticola* | 3 | 2 | 1 | 66,666667 | 33,3333333 |
| *Naubates* | 1 | 1 | 0 | 100 | 0 |
| *Neophilopterus* | 1 | 1 | 0 | 100 | 0 |
| *Neopsittaconirmus* | 16 | 9 | 7 | 56,25 | 43,75 |
| *Nesiotinus* | 4 | 4 | 0 | 100 | 0 |
| *Nothocolus* | 1 | 1 | 0 | 100 | 0 |
| *Nothocotus* | 1 | 1 | 0 | 100 | 0 |
| *Nyctibicola* | 2 | 2 | 0 | 100 | 0 |

| Louse_Genus | Total_n | Sodalis_detection | | Percentage_Negative | Percentage_Positive |
|---|---|---|---|---|---|
| | | Negative | Positive | | |
| *Olivinirmus* | 5 | 1 | 4 | 20 | 80 |
| *Ornicholax* | 6 | 5 | 1 | 83,333333 | 16,6666667 |
| *Ornithobius* | 1 | 1 | 0 | 100 | 0 |
| *Osculotes* | 3 | 3 | 0 | 100 | 0 |
| *Otidoecus* | 1 | 1 | 0 | 100 | 0 |
| *Oxylipeurus* | 5 | 4 | 1 | 80 | 20 |
| *Palmaellus* | 1 | 1 | 0 | 100 | 0 |
| *Paraclisis* | 1 | 0 | 1 | 0 | 100 |
| *Paragoniocotes* | 14 | 14 | 0 | 100 | 0 |
| *Passonomedea* | 3 | 3 | 0 | 100 | 0 |
| *Pectenosoma* | 8 | 7 | 1 | 87,5 | 12,5 |
| *Pectinopygus* | 16 | 16 | 0 | 100 | 0 |
| *Pelmatocerandra* | 1 | 1 | 0 | 100 | 0 |
| *Penenirmus* | 42 | 42 | 0 | 100 | 0 |
| *Perineus* | 1 | 1 | 0 | 100 | 0 |
| *Pessoiaella* | 1 | 1 | 0 | 100 | 0 |
| *Philoceanus* | 1 | 0 | 1 | 0 | 100 |
| *Philopterus* | 69 | 69 | 0 | 100 | 0 |
| *Physconella* | 4 | 4 | 0 | 100 | 0 |
| *Physconelloides* | 16 | 16 | 0 | 100 | 0 |
| *Picicola* | 20 | 5 | 15 | 25 | 75 |
| *Podargoecus* | 2 | 1 | 1 | 50 | 50 |
| *Priceiella* | 9 | 9 | 0 | 100 | 0 |
| *Pseudocophorus* | 1 | 0 | 1 | 0 | 100 |
| *Pseudolipeurus* | 10 | 7 | 3 | 70 | 30 |
| *Pseudonirmus* | 1 | 1 | 0 | 100 | 0 |
| *Pseudophilopterus* | 3 | 2 | 1 | 66,666667 | 33,3333333 |
| *Psittaconirmus* | 3 | 3 | 0 | 100 | 0 |
| *Psittoecus* | 3 | 3 | 0 | 100 | 0 |
| *Psophiicola* | 4 | 4 | 0 | 100 | 0 |
| *Pterocotes* | 3 | 3 | 0 | 100 | 0 |
| *Quadraceps* | 45 | 35 | 10 | 77,777778 | 22,2222222 |
| *Rallicola* | 81 | 76 | 5 | 93,82716 | 6,17283951 |
| *Resartor* | 1 | 0 | 1 | 0 | 100 |
| *Rhopaloceras* | 9 | 7 | 2 | 77,777778 | 22,2222222 |
| *Rhynchotura* | 2 | 2 | 0 | 100 | 0 |
| *Rhynonirmus* | 1 | 1 | 0 | 100 | 0 |

| Louse_Genus | Total_n | Sodalis_detection | | Percentage_Negative | Percentage_Positive |
|---|---|---|---|---|---|
| | | Negative | Positive | | |
| *Saemundssonia* | 12 | 10 | 2 | 83,333333 | 16,6666667 |
| *Saepocephalum* | 1 | 0 | 1 | 0 | 100 |
| *Splendoroffula* | 2 | 2 | 0 | 100 | 0 |
| *Strigiphilus* | 3 | 3 | 0 | 100 | 0 |
| *Strongylocotes* | 17 | 8 | 9 | 47,058824 | 52,9411765 |
| *Struthilipeurus* | 1 | 1 | 0 | 100 | 0 |
| *Struthiolipeurus* | 1 | 1 | 0 | 100 | 0 |
| *Sturnidoecus* | 5 | 3 | 2 | 60 | 40 |
| *Tinamicola* | 1 | 1 | 0 | 100 | 0 |
| *Tinamotaecola* | 4 | 3 | 1 | 75 | 25 |
| *Trabeculus* | 1 | 1 | 0 | 100 | 0 |
| *Traihoriella* | 6 | 5 | 1 | 83,333333 | 16,6666667 |
| *Trichodopeostus* | 2 | 2 | 0 | 100 | 0 |
| *Trichophilopterus* | 2 | 2 | 0 | 100 | 0 |
| *Trogoniella* | 1 | 0 | 1 | 0 | 100 |
| *Trogoninirmus* | 2 | 0 | 2 | 0 | 100 |
| *Turnicola* | 3 | 3 | 0 | 100 | 0 |
| *Turturicola* | 1 | 0 | 1 | 0 | 100 |
| *Upupicola* | 1 | 1 | 0 | 100 | 0 |
| *Vernoniella* | 2 | 2 | 0 | 100 | 0 |

**Table S3.** Details of the bird images used in Figure 1, including the English names, scientific names, artist or author of the illustrations, and credit and copyright information.

| **Common name** | **Scientific name** | **Artist** | **Credit and Copyright** |
|---|---|---|---|
| Spot-winged Antbird | *Myrmelastes leucostigma* | Hilary Burn | Illustration of Spot-winged Antbird (*Myrmelastes leucostigma*) by Hilary Burn © Lynx Nature Books and © Cornell Lab of Ornithology. |
| Blue-banded Toucanet | *Aulacorhynchus coeruleicinctis* | Al Gilbert | Illustration of Blue-banded Toucanet (*Aulacorhynchus coeruleicinctis*) by Al Gilbert © Lynx Nature Books and © Cornell Lab of Ornithology. |
| Red-whiskered Bulbul | *Pycnonotus jocosus* | Hilary Burn | Illustration of Red-whiskered Bulbul (*Pycnonotus jocosus*) by Hilary Burn © Lynx Nature Books and © Cornell Lab of Ornithology. |
| Indian Cuckooshrike | *Coracina macei* | Tim Worfolk | Illustration of Indian Cuckooshrike (Coracina macei) by Tim Worfolk © Lynx Nature Books and © Cornell Lab of Ornithology. |
| White-bellied Kingfisher | *Corythornis leucogaster* | Norman Arlott | Illustration of White-bellied Kingfisher (Corythornis leucogaster) by Norman Arlott © Lynx Nature Books and © Cornell Lab of Ornithology. |
| Velvet-fronted Nuthatch | *Sitta frontalis* | Hilary Burn | Illustration of Velvet-fronted Nuthatch (Sitta frontalis) by Hilary Burn © Lynx Nature Books and © Cornell Lab of Ornithology. |
| Silver-backed Butcherbird | *Cracticus argenteus* | Norman Arlott | Illustration of Silver-backed Butcherbird (Cracticus argenteus) by Norman Arlott © Lynx Nature Books and © Cornell Lab of Ornithology. |
| Virginia Rail | *Rallus limicola* | Norman Arlott | Illustration of Virginia Rail (Rallus limicola) by Norman Arlott © Lynx Nature Books and © Cornell Lab of Ornithology. |
| Amazonian Umbrellabird | *Cephalopterus ornatus* | Chris Rose | Illustration of Amazonian Umbrellabird (Cephalopterus ornatus) by Chris Rose © Lynx Nature Books and © Cornell Lab of Ornithology. |
| Brown Falcon | *Falco berigora* | Hilary Burn | Illustration of Brown Falcon (Falco berigora) by Hilary Burn © Lynx Nature Books and © Cornell Lab of Ornithology. |
| Yellow-rumped Cacique | *Cacicus cela* | Tim Worfolk | Illustration of Yellow-rumped Cacique (Cacicus cela) by Tim Worfolk © Lynx Nature Books and © Cornell Lab of Ornithology. |
| American Avocet | *Recurvirostra americana* | Ian Willis | Illustration of American Avocet (Recurvirostra americana) by Ian Willis © Lynx Nature Books and © Cornell Lab of Ornithology. |
| Papuan King-Parrot | *Alisterus chloropterus* | Martin Woodcock | Illustration of Papuan King-Parrot (Alisterus chloropterus) by Martin Woodcock © Lynx Nature Books and © Cornell Lab of Ornithology. |
| Australian King-Parrot | *Alisterus scapularis* | Martin Woodcock | Illustration of Australian King-Parrot (Alisterus scapularis) by Martin Woodcock © Lynx Nature Books and © Cornell Lab of Ornithology. |
| Wilson's Storm-Petrel | *Oceanites oceanicus* | Juan Varela | Illustration of Wilson's Storm-Petrel (Oceanites oceanicus) by Juan Varela © Lynx Nature Books and © Cornell Lab of Ornithology. |
| Yellow-billed Spoonbill | *Platalea flavipes* | Francesc Jutglar | Illustration of Yellow-billed Spoonbill (Platalea flavipes) by Francesc Jutglar © Lynx Nature Books and © Cornell Lab of Ornithology. |
| Common Bronzewing | *Phaps chalcoptera* | Chris Rose | Illustration of Common Bronzewing (Phaps chalcoptera) by Chris Rose © Lynx Nature Books and © Cornell Lab of Ornithology. |

| Common name | Scientific name | Artist | Credit and Copyright |
|---|---|---|---|
| Brush Bronzewing | *Phaps elegans* | Chris Rose | Illustration of Brush Bronzewing (Phaps elegans) by Chris Rose © Lynx Nature Books and © Cornell Lab of Ornithology. |
| Picui Dove | *Columbina picui* | Martin Elliott | Illustration of Picui Dove (Columbina picui) by Martin Elliott © Lynx Nature Books and © Cornell Lab of Ornithology. |
| Croaking Ground Dove | *Columbina cruziana* | Martin Elliott | Illustration of Croaking Ground Dove (Columbina cruziana) by Martin Elliott © Lynx Nature Books and © Cornell Lab of Ornithology. |
| Band-tailed Pigeon | *Patagioenas fasciata* | Jan Wilczur | Illustration of Band-tailed Pigeon (Patagioenas fasciata) by Jan Wilczur © Lynx Nature Books and © Cornell Lab of Ornithology. |
| Little Tinamou | *Crypturellus soui* | Lluís Sanz | Illustration of Little Tinamou (Crypturellus soui) by Lluís Sanz © Lynx Nature Books and © Cornell Lab of Ornithology. |
| Crested Auklet | *Aethia cristatella* | Chris Rose | Illustration of Crested Auklet (Aethia cristatella) by Chris Rose © Lynx Nature Books and © Cornell Lab of Ornithology. |

**Figure S1.** Phylogenetic tree of feather-feeding lice (Ischnocera) based on a partitioned IQ-TREE maximum likelihood (ML) analysis of a concatenated matrix of 2,359 single-copy ortholog genes. Bootstrap values are 100%, except where indicated. Hash marks on branch leading to *Quadraceps hospes* shorten the visually long branch resulting from missing data for this species. The tree topology is rotated to match the structure of the louse tree in the tanglegram shown in Figure 2. Tree rooted on Anoplura (*Proechinophthirus fluctus*), not shown.

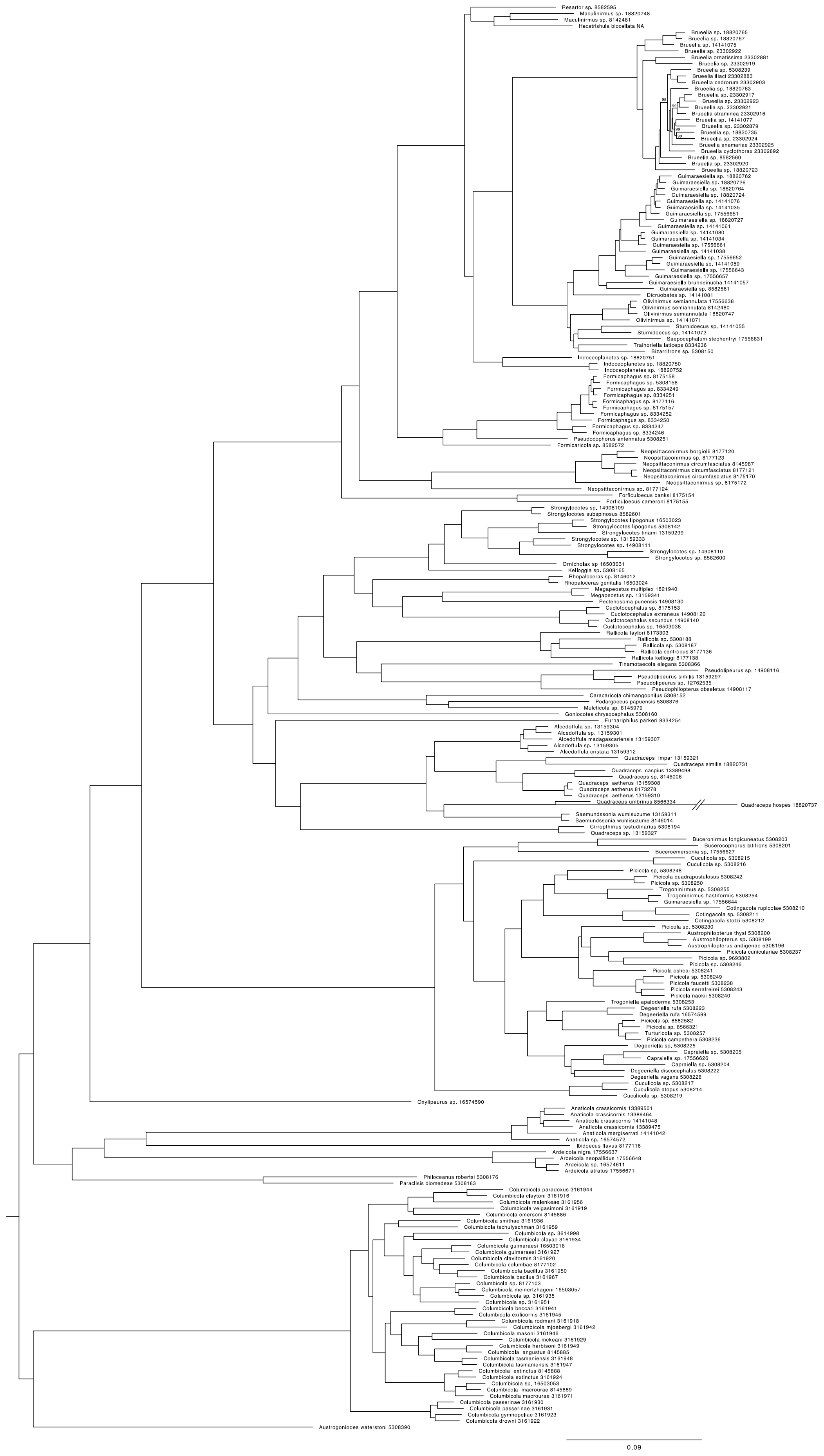
